# A joint learning approach for genomic prediction in polyploid grasses

**DOI:** 10.1101/2022.04.13.488210

**Authors:** Alexandre Hild Aono, Rebecca Caroline Ulbricht Ferreira, Aline da Costa Lima Moraes, Letícia Aparecida de Castro Lara, Ricardo José Gonzaga Pimenta, Estela Araujo Costa, Luciana Rossini Pinto, Marcos Guimarães de Andrade Landell, Mateus Figueiredo Santos, Liana Jank, Sanzio Carvalho Lima Barrios, Cacilda Borges do Valle, Lucimara Chiari, Antonio Augusto Franco Garcia, Reginaldo Massanobu Kuroshu, Ana Carolina Lorena, Gregor Gorjanc, Anete Pereira de Souza

**Affiliations:** Molecular Biology and Genetic Engineering Center (CBMEG), University of Campinas (UNICAMP), Campinas, SP, Brazil; The Roslin Institute, The University of Edinburgh, Midlothian, United Kingdom (UK); Genetics Department, Luiz de Queiroz College of Agriculture (ESALQ), University of São Paulo (USP), Piracicaba, SP, Brazil; Instituto de Ciência e Tecnologia (ICT), Universidade Federal de São Paulo (UNIFESP), São José dos Campos, SP, Brazil; Advanced Center of Sugarcane Agrobusiness Technological Research, Agronomic Institute of Campinas (IAC), Ribeirão Preto, SP, Brazil; Embrapa Beef Cattle, Brazilian Agricultural Research Corporation, Campo Grande, MS, Brazil; Technological Institute of Aeronautics, São José dos Campos, SP, Brazil; Department of Plant Biology, Institute of Biology (IB), University of Campinas (UNICAMP), Campinas, SP, Brazil

## Abstract

Poaceae, among the most abundant plant families, includes many economically important polyploid species, such as forage grasses and sugarcane (*Saccharum* spp.). These species have elevated genomic complexities and limited genetic resources, hindering the application of marker-assisted selection strategies. Currently, the most promising approach for increasing genetic gains in plant breeding is genomic selection. However, due to the polyploidy nature of these polyploid species, more accurate models for incorporating genomic selection into breeding schemes are needed. This study aims to develop a machine learning method by using a joint learning approach to predict complex traits from genotypic data. Biparental populations of sugarcane and two species of forage grasses (*Urochloa decumbens, Megathyrsus maximus*) were genotyped, and several quantitative traits were measured. High-quality markers were used to predict several traits in different cross-validation scenarios. By combining classification and regression strategies, we developed a predictive system with promising results. Compared with traditional genomic prediction methods, the proposed strategy achieved accuracy improvements exceeding 50%. Our results suggest that the developed methodology could be implemented in breeding programs, helping reduce breeding cycles and increase genetic gains.

## Introduction

Poaceae (Gramineae) is a very diverse family of flowering plants known as grasses with unquestionable economic importance, as they are sources of food, fuel and fodder. Among Poaceae crop species, sugarcane (*Saccharum* spp., SU) stands out as responsible for most of the global sugar production^1,2^ and for its high energy potential^3^, while tropical forage grasses (FGs), including *Urochloa* spp. and *Megathyrsus maximus* (syn. *Panicum maximum*), are the main food source for beef and dairy cattle in tropical and subtropical regions of the world^4,5^. In addition to their agricultural relevance, SU and FGs have complex and large polyploid genomes, making the application of marker-assisted selection more challenging than in diploid grasses such as rice, maize and sorghum^6–8^.

As with all crops, many traits of great agricultural importance are highly quantitative and controlled by many loci of very small effect and thus are easily affected by environmental conditions^9^. However, in these polyploid species, the effect of individual loci tends to be even smaller, as it is distributed among several alleles with different possible combinations, making the genotype-phenotype correspondence uncertain^10–12^. This fact, combined with other characteristics, such as the large and heterozygous nature of their genomes, which have a high content of repetitive regions^7,13^, explains the shortage of genomic resources for answering relevant biological questions and enabling the advance of the genetic improvement of these crops.

With recent advances in high-throughput sequencing and phenotyping technologies, a compelling approach known as genomic selection (GS) has been used to accelerate selection cycles through the application of statistical methods to predict the breeding value of individuals by using molecular markers, mainly single nucleotide polymorphisms (SNPs)^14^. First proposed by Meuwissen et al.^15^, this approach has shown great potential in studies on polyploid species of economic importance, as it provides relevant genetic gains considering complex traits controlled by several genes of minor effect^16–20^. However, there is still a gap in effectively applying GS in SU and FG breeding programs because no available strategy has shown enough improvement in accuracy to significantly increase yearly genetic gains^21–23^.

Among other factors, the accuracies of predictive models vary depending on the statistical and computational methods employed^24^. However, the several methods available for GS do not work satisfactorily for complex polyploids such as the species mentioned here, and as a consequence, the prediction of marker-trait associations is not accurate. Although representing a valuable alternative to improve predictive accuracy in GS, traditional machine learning (ML) methods have also been shown to be inefficient in increasing genetic gains and, thus, solving the existing limitations to incorporating GS into complex polyploid breeding schemes^25,26^. Therefore, there is a need to incorporate different learning strategies and algorithms into GS to improve predictive performance for quantitative traits in genetically complex species such as SU and FGs.

In this context, this study is aimed at the development of a predictive system for genomic prediction (GP) of complex traits in polyploid grasses, especially species with incomplete reference genomes and scarce genetic resources. Considering the difficulty of capturing species’ genomic singularities through linear regression techniques and creating predictive models with high accuracies, we incorporated different ML methodologies into an ensemble system composed of classification and regression models. The applicability and potential of this joint learning approach were assessed with several complex traits from three different biparental populations of SU and FGs (*U. decumbens* and *M. maximus*) and contrasted with those of traditional linear regression models for GS. Our study provides a novel methodology with strong potential to be implemented in polyploid grass breeding programs, assisting in reducing breeding cycles and increasing genetic gain.

## Material and Methods

All the predictive models used in this study were applied to three different biparental populations: (Pop1) 219 individuals generated from a cross between SU commercial varieties, (Pop2) 239 individuals from a cross between a commercial variety and a genotype of *U. decumbens* (2n = 4x = 36), and (Pop3) 136 individuals from a cross between a commercial variety and a genotype of *M. maximus* (2n = 4x = 32). The same data analysis procedures were applied for all populations, with specific differences regarding the genotyping and bioinformatics procedures. Different traits were evaluated for these species, and therefore, specific mixed effect models were estimated for each scenario.

### Grass Populations

Pop1 hybrids were generated from a cross between the elite clone IACSP95-3018 (female parent) and the commercial variety IACSP93-3046 (male parent). This population was developed by the Sugarcane Breeding Program of the Agronomic Institute of Campinas^27^. This population was planted in three locations: (i) Ribeirão Preto, São Paulo, Brazil (21°12^*′*^ 0.6^*′′*^ S, 47°52^′^ 21.5^′′^ W, 546.8 m), in 2005; (ii) Sales de Oliveira, São Paulo, Brazil (20°46^′^ 19^′′^ S, 47°50^′^ 16^′′^ W, 730 m), in 2007; and (iii) Piracicaba, São Paulo, Brazil (22°42^′^ 36.6^′′^ S, 47°37^′^ 58.3^′′^ W, 554 m), in 2011. Experimental fields were established following an augmented block design with five blocks (44 individuals per block) and plots with 1-meter rows and 1.5 m plant spacing. Each individual was evaluated for stalk diameter (SD) and stalk height (SH) in a sample with 10 stalks from individual plots^28^. These evaluations were performed in 2008 (plant cane) and 2009 (ratoon cane) in location (ii) and (iii), and in 2012 (plant cane), 2013 (ratoon cane) and 2014 (ratoon cane) in location (i).

Pop2 and Pop3 were generated and evaluated by the Embrapa Beef Cattle (Brazilian Agricultural Research Corporation), located in Campo Grande, Mato Grosso do Sul, Brazil (20°27^′^ S, 54°37^′^ W, 530 m). Pop2 originated from a cross between *U. decumbens* D62 (cv. Basilisk) as the male apomictic parent and *U. decumbens* D24/27 as the female sexual parent obtained by tetraploidization with colchicine^29^. Plants were evaluated for the following agronomic traits: regrowth capacity (RC), field green weight in kg/ha (FGW), total dry matter (TDM) in kg/ha, leaf dry matter (LDM) in kg/ha, leaf percentage (LP), and the leaf stem ratio (LSR), as described by Mateus et al.^30^. Briefly, the field experimental design followed a simple lattice of 18 × 18 meters with two replications, seven controls and five plants per plot. These traits were evaluated for seven harvests in 2012: two in the dry season and five in the rainy season.

Pop3 was obtained from a cross between *M. maximus* cv. Mombaça, the apomictic parent, and an obligate sexual parent, *M. maximus* S10. According to Deo et al.^31^, this population was evaluated in an augmented block design with 160 regular treatments distributed in eight blocks with two replicates. Each block was composed of 22 plots with 20 individuals and two controls. Genotypes were evaluated for several agronomic traits: green matter (GM), stem dry matter (SDM), percentage of leaf blade (PLB), TDM, LDM, and RC. Phenotyping was performed in six harvests (three in 2013 and three in 2014), with the exception of phenotyping of RC, which was performed for three harvests (one in 2013 and two in 2014).

### Phenotypic Data Analyses

Raw phenotypic data were processed according to the mixed linear models described by Aono et al.^27^, Ferreira et al.^32^, and Deo et al.^31^ for Pop1, Pop2 and Pop3 respectively. Trait heritabilities were calculated based on model variances considering 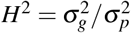, where 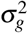 and 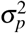 are the genetic and phenotypic variances, respectively. Pop1 data were evaluated in R statistical software^33^ using breedR v.0.12^34^ and bestNormalize^35^ for data normalization. We modeled each trait considering the following statistical model:

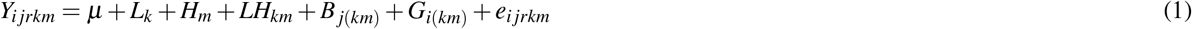

where each trait measure *Y*_*ijrkm*_ of the *r*th replicate of the *i*th genotype was evaluated at the *k*th location of the *j*th block in the *m*th year. The trait mean is given by *µ*, and the contributions of the *k*th location (*L*_*k*_), the *m*th harvest (*H*_*m*_), the *j*th block at the *k*th location in the *m*th year (***B***_*j*(*km*)_), and the interaction between the *k*th location and *m*th harvest (***LH***_*km*_) were modeled as fixed effects. The effects of the genotype *G*_*i*(*km*)_ and the residual error *e*_*i jrkm*_ were included as random effects.

Pop2 phenotypic evaluations were performed using ASReml-R v3^36,37^ and ASRemlPlus^38^ software. The traits were modeled with the following model:

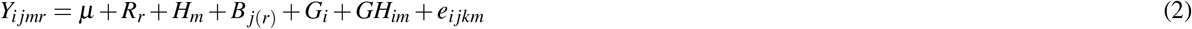

where each trait measure *Y*_*ijmr*_ of the *i*th genotype was evaluated in the *j*th block in the *m*th harvest considering the *r*th replicate. The trait mean is given by *µ*, and the contributions of the *r*th replicate (***R***_*r*_) and the *m*th harvest (***H***_*m*_) were modeled as fixed effects. The contributions of the *i*th genotype (***G***_*i*_), the *j*th block in the *r*th replicate (***B***_*j*(*r*)_), and the interaction between the *i*th genotype and the *m*th harvest (***GH***_*im*_) were included as random effects, as well as the residual error *e*_*ijkm*_.

For Pop3, a longitudinal linear mixed model was fit using ASReml-R^36^, with Box-Cox transformation^39^ for data normalization. The following model was employed:

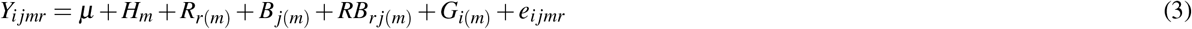

where each trait measure *Y*_*ijmr*_ of the *i*th genotype was evaluated in the *j*th block in the *m*th harvest considering the *r*th replicate. The trait mean is given by *µ*, and the contributions of the *m*th harvest (***H***_*m*_) and the *r*th replicate in the *m*th harvest (*R*_*r*(*m*)_) were modeled as fixed effects. The contributions of the *j*th block in the *m*th harvest (***B***_*j*(*m*)_), the interaction of the *r*th replicate and *j*th block in the *m*th harvest (***RB***_*r j*(*m*)_), the *i*th genotype (or control) in the *m*th harvest (*G*_*i*(*m*)_) and the environmental error (*e*_*ijmr*_) were modeled as random effects.

The best linear unbiased predictors (BLUPs) (Pop1) and corrected phenotypes (Pop2 and Pop3) were estimated, rescaled to a new range of values between 0 and 1 using min-max normalization and used for phenotypic predictions. For a set of values *v*, the transformed values *f* (*v*) are:

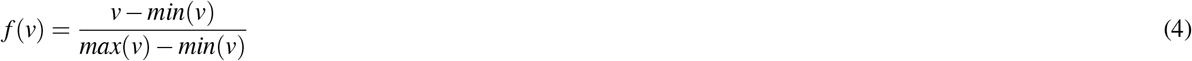

Visual inspections of phenotypic data distributions were performed using the ggplot2 package^40^ in R. Trait correlations were estimated with R Pearson correlation coefficients by using the PerformanceAnalytics package^41^ in R.

### Genotyping Procedures

As the genotyping approach, we employed a genotyping-by-sequencing (GBS) strategy. GBS libraries were prepared following the protocol of Elshire et al.^42^ for Pop1 and Pop2 and the protocol of Poland et al.^43^ for Pop3. DNA extraction from Pop1 was performed using a cetyltrimethylammonium bromide (CTAB)-based protocol^44^, and two 96-plex GBS libraries were constructed using PstI, with two samples per parent. Five sequencing runs were performed with the Illumina GAIIx and Illumina NextSeq systems^27^. Genomic DNA of Pop2 was extracted using the DNeasy Plant Kit (QIAGEN, Hilden, Germany). Two 96-plex GBS libraries were constructed using NsiI and five replicates per parent and sequenced on the NextSeq 500 platform (Illumina, San Diego, CA, USA)^32^. DNA extraction from Pop3 was performed following a CTAB-based method^45^ with slight modifications. Then, 96-plex GBS libraries were constructed using a combination of a rarely cutting enzyme (PstI) and a frequently cutting enzyme (MspI), with 12 replicates for each parent. Pop3 libraries were also sequenced on the NextSeq 500 platform^31^.

SNP calling in SU was performed by coupling several bioinformatics pipelines and selecting the intersecting markers by the TASSEL-GBS v.4 pipeline^46^ modified for polyploids^47^ and at least one other SNP caller, including SAMtools version 1.6^48^, Stacks version 2.3^49^, Genome Analysis Toolkit (GATK) version 3.7 with the Haplotype Caller algorithm^50^, and FreeBayes version 1.1.0-3^51^. GBS reads were filtered and preprocessed using FastX-Toolkit scripts^52^, and comparative alignments were performed against the PhiX genome using BLASTn^53^, as described by Aono et al.^27^. Read alignment was performed with BWA-MEM version 0.7.12^54^ and the methyl-filtered genome of the sugarcane cultivar SP70-1143^55^, retaining only uniquely mapped reads. For GATK, SAMtools and FreeBayes, a common pipeline was established.

For Pop2 and Pop3, SNP calling was performed using TASSEL-4-POLY. Because this pipeline requires a reference genome for SNP calling, we used the *Setaria viridis* v1.0 and *Panicum virgatum* v1.0 genomes as references to align the GBS reads of Pop2 and Pop3, respectively, in Bowtie2 v.2.3.1^56^. Both reference genomes were produced by the US Department of Energy Joint Genome Institute and are available from the Phytozome database (http://phytozome.jgi.doe.gov/)^57^. In this step, the following settings were used for both populations: very-sensitive-location, a limit of 20 dynamic programming problems and a maximum of 4 times to align a read.

After selecting the SNP sets for each population, we kept only biallelic variants with a minimum depth of 50 reads per individual and a maximum of 25% missing data. These markers were represented as allele proportions by calculating the ratio between the number of reads for the reference allele and the total number of reads^27^. Missing data were imputed as means.

### Multivariate Analyses

To evaluate genotypic and phenotypic data profiles across populations, we performed principal component analyses (PCAs), t-distributed stochastic neighbor embedding (t-SNE) analyses^58^, and redundancy analyses (RDAs)^59^. Scatter plots were constructed using the ggplot2 R package. We evaluated phenotypic data using PCA and genotypic data with t-SNE in the Rtsne R package^60^, considering a perplexity of 30. We combined both datasets with RDA, which was implemented in the vegan R package^61^. RDA constrained axes were evaluated with the analysis of variance (ANOVA) F-statistic, and to identify SNPs putatively under association with the phenotypes, we used a three-standard deviation cut-off (p-value of 0.0027).

Additionally, we performed a set of clustering analyses for each trait, considering hierarchical (Ward’s minimum variance with and without squared dissimilarities, herein referred to as WardWSD and WardWoSD, respectively), single, complete, unweighted pair-group method with arithmetic averaging (UPGMA), weighted pair-group method with arithmetic averaging (WPGMA), weighted pair-group method with centroid averaging (WPGMC), unweighted pair group method with centroid averaging (UPGMC) and nonhierarchical (K-means) strategies together with pairwise Euclidean distances. For each of the clustering methods and traits, we calculated the clustering index by using the NbClust R package^62^, defining the best clustering scheme as that most indicated by the indexes in a cluster number range of 2 to 10. Cluster visualization was performed with the ggtree R package^63^ and used to create intervals for each phenotype. The best method was defined through visual inspections considering the trait distribution and the cluster configurations on the heatmaps created.

### Single Regression Genomic Prediction

We evaluated the prediction accuracies of traditional models for GP and ML-based approaches. As traditional approaches, we selected (a) Bayesian ridge regression (BRR)^64^; (b) Bayesian reproducing kernel Hilbert spaces regression with kernel averaging (RKHS-KA)^65^, considering different kernels with bandwidth parameters of 15*M*, 5*M*, and 1*M*, where *M* represents the median squared Euclidean distance across genotypes; and (c) a single-environment, main genotypic effect model with a Gaussian kernel (SM-GK)^66^. BRR and RKHS-KA models were implemented in the BGLR R package^67^, and the SM-GK model was implemented in the BGGE R package^66^. Given a genotype matrix (SNP data), codified using allele proportions, with *p* loci and *n* individuals, the BRR models were estimated considering the following equation:

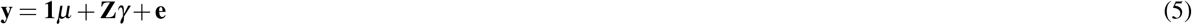

where **y** represents the response measures, *µ* the overall population mean, **e** the residuals 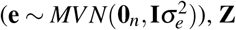, **Z** the genotype matrix, and *γ* the vector of SNP effects with the same normal prior distribution for all loci 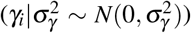 and the hyperparameter following a scaled inverse chi-squared hyperprior distribution 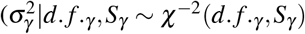, where 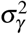 is the genetic variance, *d. f*. the degrees of freedom and *S*_*γ*_ the scale parameter). In SM-GK, considering the same equation, **Z** is the incidence matrix for random genetic effects, and *γ* the genetic effects considering 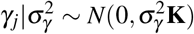, where the matrix **K** is computed through a Gaussian kernel (GK) function. For pairwise **K** calculations between two genotypes *x*_*i*_ and *x*_*i*′_, 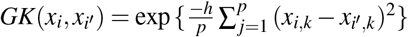, where *h* is the bandwidth parameter (herein considered 1 for SM-GK). In RKHS-KA, a multikernel model is fitted with the same GK but considering separate random effects for three established kernels. The posterior distribution of the models was assessed with the Gibbs sampler, using 20,000 interactions and discarding 2,000 cycles (burn-in).

For ML-based approaches, we selected regression methods based on the following algorithms: (i) k-nearest neighbors (KNN)^68^, (ii) support vector machine (SVM)^69^, (iii) random forest (RF)^70^, and (iv) AdaBoost^71^. All of these algorithms were implemented with the scikit-learn Python v.3 module^72^. For KNN (*k* = 5), pairwise Euclidean distances with uniform weights were considered. In SVM regression, a radial basis function kernel (*K*_*SVM*_) was used. Considering two genotypes *x*_*i*_ and *x*_*i*_′, *K*_*SVM*_(*x*_*i*_, *x*_*i*_′) = exp (−*θ* ||*x* − *x*′||^2^), with the free parameter *θ* defined as 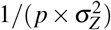, where *p* represents the number of loci and 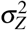 the variance of the genotype matrix **Z**. The RF meta-estimator was built considering 100 decision trees for random data splits evaluated through mean squared errors (MSEs) calculated as 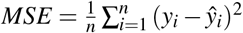, where *y*_*i*_ and ŷ_*i*_ are the real and the predicted values, respectively, across *n* individuals in data split *k* (from 1 to 100). Tree induction was performed considering a maximum tree height of *p* (the number of loci). For AdaBoost, the base estimator was a decision tree regressor with a linear loss function for weighting boosting interactions.

All these models were evaluated under a cross-validation (CV) scenario with 50 10-fold repeats and the leave-one-out (LOO) approach. For each trait, predictive accuracies were estimated as MSEs and R Pearson correlations and visualized with the ggplot2 R package. The models’ performances were compared with ANOVA followed by Tukey’s multiple comparisons test (p-value of 0.05), implemented in the agricolae R package^73^.

### Feature Selection and Marker Annotation

In this work, we used feature selection (FS) techniques to extract loci with putative phenotypic associations considering higher predictive accuracies. For this task, we tested the performance of ML-based regression and classification models for predicting the phenotypes and their interval values considering scenarios with and without FS. The predictive capabilities were estimated using 10-fold CV repeated 50 times, with stratification in the interval classification method. For regression, the models were created using the KNN, SVM, RF, and AdaBoost approaches. For classification, the (i) SVM^69^, (ii) RF^70^, (iii) multilayer perceptron (MLP) neural network^74^, and (iv) Gaussian naive Bayes (GNB)^75^ ML algorithms were used.

For classification FS, we employed the approach described by Aono et al.^27^. As an FS system, we combined the results of (i) L1-based FS with a linear support vector classification system^69^, (ii) ANOVA-based univariate FS (p-value of 0.05), and (iii) gradient tree boosting (GTB)^76^. In this approach, a locus is retrieved by the system if identified as relevant for classification in at least two out of the three FS techniques employed. This methodology was extended for regression FS, considering models adapted for a continuous response variable. Techniques (i) and (iii) were changed, considering SVM and GTB for regression, and approach (ii) was replaced with Pearson correlation (p-value cut-off of 0.05).

For each phenotype, we retrieved the selected markers and obtained a genomic window of 1,000 base pairs upstream and downstream in the reference genome. With these sequences, we performed comparative alignments against coding DNA sequences (CDSs) of fourteen different species from the *Poaceae* family (*Brachypodium distachyon, B. hybridum, B. silvatium, Hordeum vulgare, Oryza sativa, Oropetium thomaeum, Panicum hallii, P. virgatum, Sorghum bicolor, Setaria italica, S. viridis, Triticum aestivum, Thinopyrum intermedium* and *Zea mays*) and *Arabidopsis thaliana* extracted from the Phytozome v.13 database^57^. All the correspondences were used to summarize related Gene Ontology (GO) terms using the Revigo tool^77^ and Cytoscape software^78^.

### Joint Learning

With the selected traits, we contrasted the GP methods using traditional statistical methodologies (BRR, RKHS-KA, and SM-GK) and the entire SNP dataset, with the construction of SM-GK models based on the SNPs selected from the FS approaches. We examined FS-selected SNPs considering the established classification (C) and regression (R) techniques, and used the SNPs identified in the intersection of at least two (C2 or R2) or three (C3 or R3) FS methodologies. The tested combinations of models were (i) SM-GK estimated based on C2; (ii) C3; (iii) R2; (iv) R3; (v) the intersection of C2 and R2 (ICR2); (vi) the union of C2 and R2 (CR2); (vii) the union of C3 and R3 (CR3); (viii) the entire set of markers including C3 as fixed effects (C3F); and (ix) the entire set of markers including R3 as fixed effects (R3F).

The inclusion of markers as fixed effects in SM-GK models was performed with the equation:

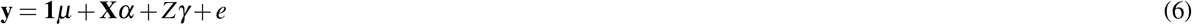

where *y* represents the response measures, *µ* the overall population mean, *X* the genotype matrix with the FS-associated markers, *Z* the incidence matrix for random genetic effects, and **e** the residuals. The marker effects in the FS set are represented by *α*, and the genetic effects of *γ* follow the distribution described above.

To compare predictive performance between the developed models and the other techniques, we employed the same CV scenarios described above (LOO and 10-fold), retrieving the MSE measures and the Pearson correlation coefficients. Correlation plots were constructed with the ggplot2 R package.

### Simulation Approaches

To evaluate the proposed approach in larger populations in a controlled scenario, we performed simulations using AlphaSimR software^79^. We selected an entire wheat breeding program^80,81^ for creating genotypic (21 chromosomes with 1,000 SNPs each and coded as 0 (ancestral homozygote), 1 (heterozygote) and 2 (derived homozygote)) and phenotypic data (yield). For simulation scenarios, we considered different quantities of QTLs per chromosome (2, 10, 100 and 1,000) with additive effects on yield sampled from a normal distribution (mean of 4 and variance of 0.1). Using such a configuration, 70 inbred lines were created and used for 20 years of phenotypic selection, with the first ten years discarded as burn-in. The environmental variance was set to 0.4 and the genotype-environment interaction variance was set to 0.2.

The inbred lines were used to create 100 biparental populations with 100 individuals each, for a total of 10,000 genotypes. These were selected using (i) a preliminary yield trial (PYT), (ii) an advanced yield trial (AYT), and (iii) an elite yield trial (EYT). To advance a genotype to the PYT, we created double haploids of the 10,000 lines, visually evaluated in a head row nursery (heritability of 0.1) to select the best 1,000 genotypes (10 per family). The best 100 lines in the PYT were selected for the AYT considering an unreplicated trial (heritability of 0.2), and the best 10 in the AYT were advanced to the EYT using a small multilocation replicated trial (heritability of 0.5). From the 10 lines in the EYT, the released variety was selected using a large multilocation replicated trial in 2 consecutive years (heritability of 0.67).

From the last ten years of phenotypic selection, we selected the genotypic data of the PYT with the corresponding phenotypes simulated to create datasets for evaluating the proposed approach. From a total of 10,000 lines, we sampled different numbers of individuals (100, 250, 500, 1,000, 2,000, 5,000 and 10,000) and performed 10-fold CV evaluations for GP.

## Results

### Phenotypic and Genotypic Data Analyses

From the defined mixed models, we could estimate, for each trait and population, the corresponding BLUPs and corrected phenotypes (Supplementary Figs. 1-3), which were used for GP. Although there were some evident correlations between traits (Supplementary Figs. 4-6), most of the evaluated phenotypes clearly represented a different prediction task for the development of models. There was a variable range of heritabilities associated with each trait, ranging from 0.49 to 0.51 in Pop1, from 0.52 to 0.81 in Pop2, and from 0.19 to 0.64 in Pop3 (Table 1).

**Table 1.**
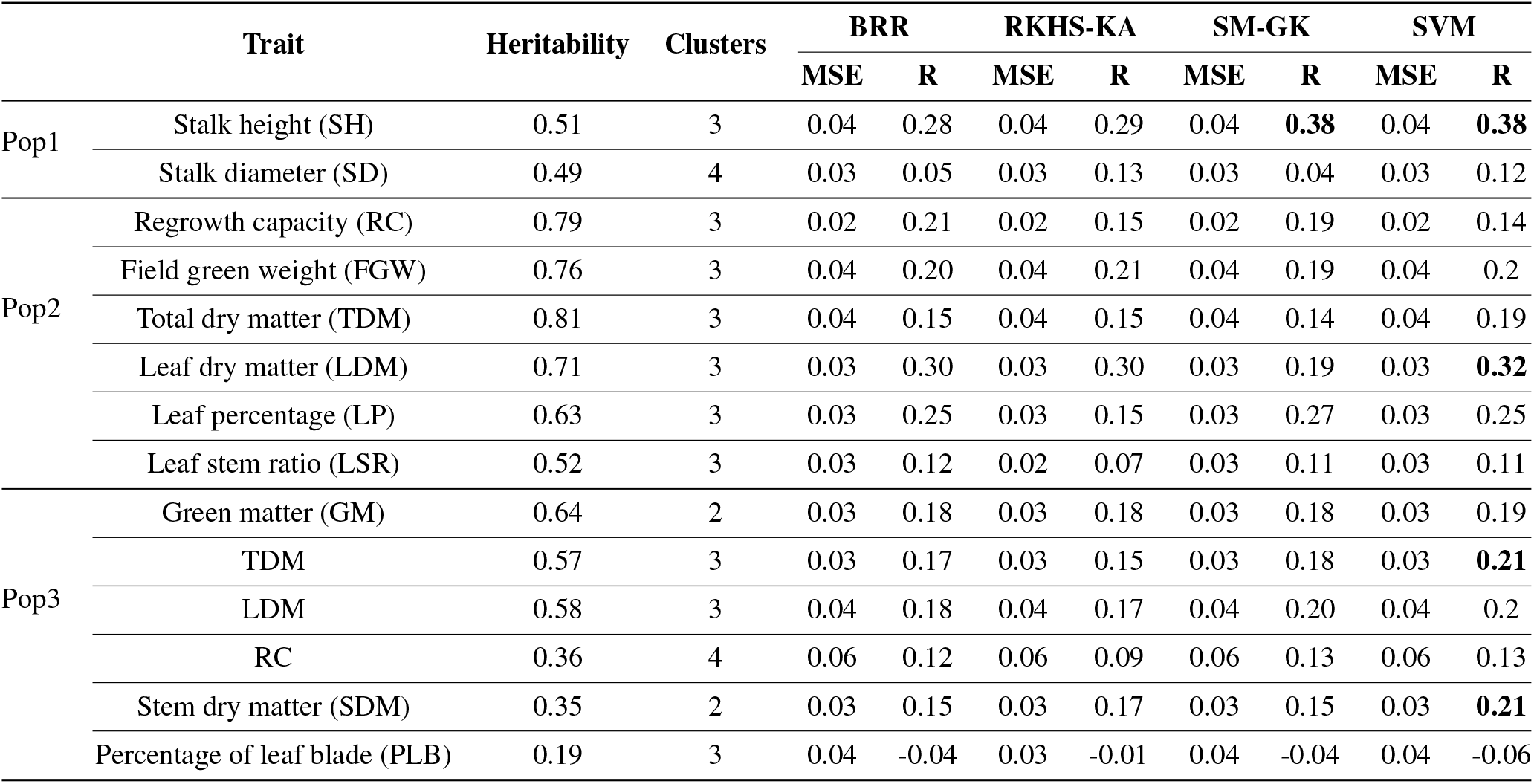
Accuracies for Bayesian ridge regression (BRR), the reproducing kernel Hilbert space with kernel averaging (RKHS-KA) model, the single-environment, main genotypic effect model with a Gaussian kernel (SM-GK), and support vector machine (SVM)-based regression across the different traits separated according to the populations: (Pop1) cross between sugarcane commercial varieties, (Pop2) cross between a commercial variety and a genotype of *Urochloa decumbens*, and (Pop3) cross between a commercial variety and a genotype of *Megathyrsus maximus*. The accuracies are estimated through mean squared errors (MSE) and Pearson correlation coefficients (R). The heritability and cluster configuration of each trait are also presented. The values in bold represent the top 5 accuracies observed.

A total of 182 (out of 219), 219 (out of 239) and 138 (out of 138) individuals were genotyped for Pop1, Pop2, and Pop3, respectively. For each population, we generated the following number of filtered biallelic SNP markers: (i) 14,540 in Pop1 (from a total of 137,757 raw SNPs), (ii) 4,548 in Pop2 (from 9,354), and (iii) 5,137 in Pop3 (from 23,637). For all these SNP sets, we observed a significant loss in the number of filtered markers (90% for Pop1, 51% for Pop2, and 78% for Pop3), which was expected due to the large quantity of missing data in GBS experiments and the pseudoreference genomes employed in the SNP calling step.

For each of the traits evaluated, we used nine different clustering methods and considered the group configuration as that most indicated by clustering indexes calculated with the NbClust package (with the cluster number ranging between 2 and 10). All nine clustering schemes identified for each trait were compared through visual inspections using dendrograms and heatmaps (Supplementary Figs. 7-9). The most suitable group configuration for separating the phenotypic values was assigned for each trait, resulting in different cluster quantities (Table 1). In Pop1, the WardWoSD method was the best clustering strategy. For Pop2, the WardWoSD method was also most appropriate for most of the traits, except for LSR and TDM, which were evaluated using hierarchical complete clustering and the WardWSD approach, respectively. For Pop3, all the traits were evaluated with the WardWSD approach.

To provide an overview of the genotypic and phenotypic data, we performed different multivariate analyses. For FG populations, contaminating individuals might be introduced during crosses, which could be detected by such analyses^82^. As expected, we found putative contaminants in Pop2 though t-SNE analysis performed on genotypic data (Supplementary Fig. 10). The individuals with distinct profiles across the Cartesian plane were excluded from the dataset and not considered in further analyses (18 genotypes located apart from most of the population in the t-SNE biplot were discarded).

With the final phenotypic datasets, we performed PCAs (Fig. 1), which were used to color and size the t-SNE results from genotypic data. These plots demonstrate the lack of correspondence between the genotypic and phenotypic datasets, corroborating the difficulty of developing predictive models for such associations in the populations under analysis. By considering linear dependency across SNPs and traits, RDAs were performed, allowing the identification of a small number of putative linear associations (233 in Pop1, 37 in Pop2, and 21 in Pop3), evidencing the need for alternative approaches for categorizing the genomic regions linked to the phenotypes analyzed.

**Figure 1.**
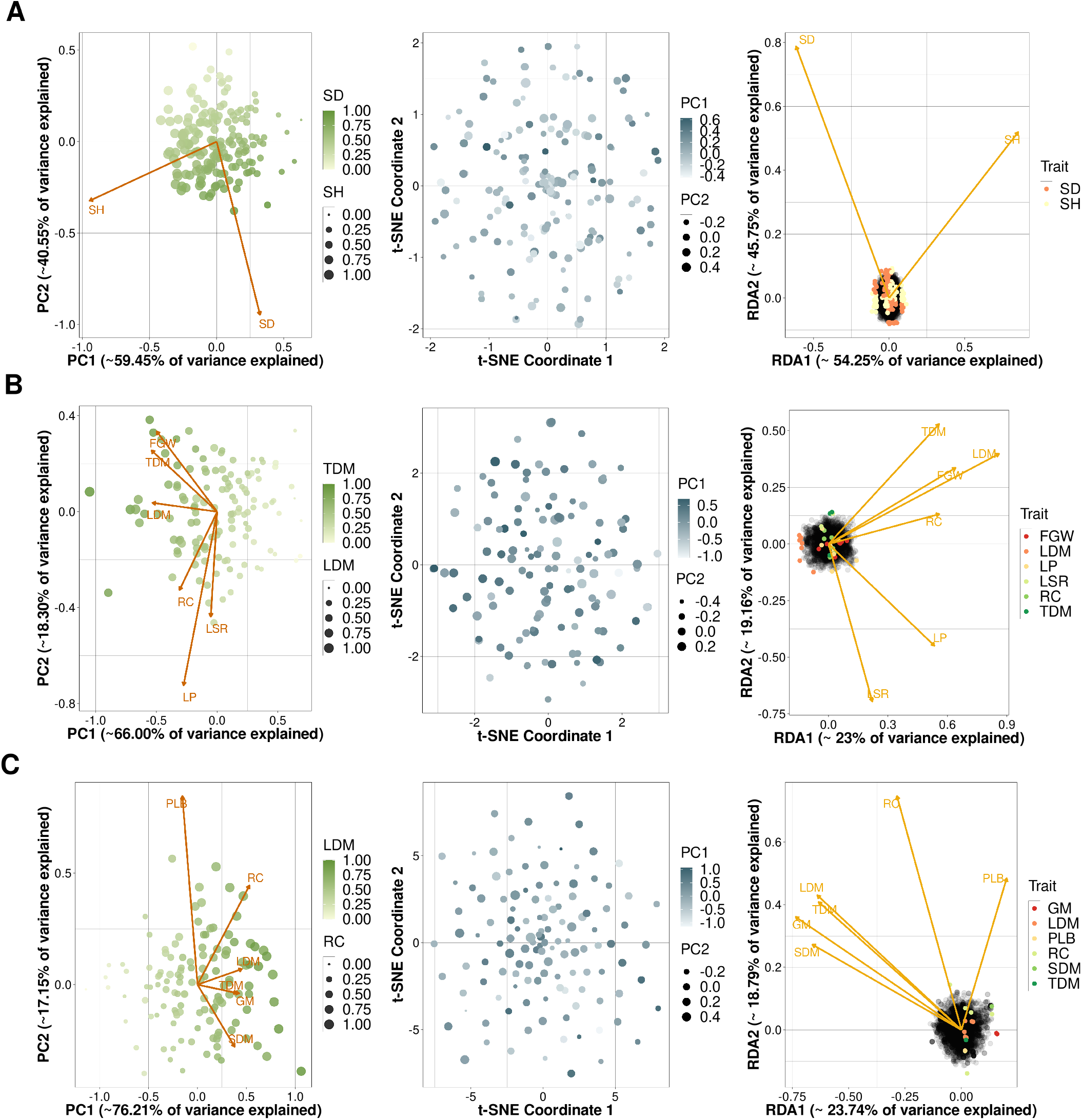
Multivariate analyses performed on both genotypic and phenotypic data for the populations of (A) sugarcane, (B) *Urochloa decumbens*, and (C) *Megathyrsus maximus*. Principal component analyses (PCAs) were performed with phenotypic data; t-distributed stochastic neighbor embedding (t-SNE), with genotypic data; and redundancy analyses (RDAs), with both datasets.

A total of 182 (out of 219), 219 (out of 239) and 138 (out of 138) individuals were genotyped for Pop1, Pop2, and Pop3, respectively. For each population, we generated the following number of filtered biallelic SNP markers: (i) 14,540 in Pop1 (from a total of 137,757 raw SNPs), (ii) 4,548 in Pop2 (from 9,354), and (iii) 5,137 in Pop3 (from 23,637). For all these SNP sets, we observed a significant loss in the number of filtered markers (90% for Pop1, 51% for Pop2, and 78% for Pop3), which was expected due to the large quantity of missing data in GBS experiments and the pseudoreference genomes employed in the SNP calling step.

For each of the traits evaluated, we used nine different clustering methods and considered the group configuration as that most indicated by clustering indexes calculated with the NbClust package (with the cluster number ranging between 2 and 10). All nine clustering schemes identified for each trait were compared through visual inspections using dendrograms and heatmaps (Supplementary Figs. 7-9). The most suitable group configuration for separating the phenotypic values was assigned for each trait, resulting in different cluster quantities (Table 1). In Pop1, the WardWoSD method was the best clustering strategy. For Pop2, the WardWoSD method was also most appropriate for most of the traits, except for LSR and TDM, which were evaluated using hierarchical complete clustering and the WardWSD approach, respectively. For Pop3, all the traits were evaluated with the WardWSD approach.

To provide an overview of the genotypic and phenotypic data, we performed different multivariate analyses. For FG populations, contaminating individuals might be introduced during crosses, which could be detected by such analyses^82^. As expected, we found putative contaminants in Pop2 though t-SNE analysis performed on genotypic data (Supplementary Fig. 10). The individuals with distinct profiles across the Cartesian plane were excluded from the dataset and not considered in further analyses (18 genotypes located apart from most of the population in the t-SNE biplot were discarded).

With the final phenotypic datasets, we performed PCAs (Fig. 1), which were used to color and size the t-SNE results from genotypic data. These plots demonstrate the lack of correspondence between the genotypic and phenotypic datasets, corroborating the difficulty of developing predictive models for such associations in the populations under analysis. By considering linear dependency across SNPs and traits, RDAs were performed, allowing the identification of a small number of putative linear associations (233 in Pop1, 37 in Pop2, and 21 in Pop3), evidencing the need for alternative approaches for categorizing the genomic regions linked to the phenotypes analyzed.

### Single Regression Genomic Prediction

The predictive performance for each trait was evaluated with seven different approaches for the regression task: BRR, RKHS-KA, SM-GK, AdaBoost, KNN, RF, and SVM. By employing 10-fold CV 50 times, we observed superior results for BRR, RKHS-KA, SM-GK and SVM (Supplementary Figs. 11-13, Supplementary Table 1). Although not all four methods ranked best in terms of predictive performance for SH (Pop1), RC (Pop2), and PLB (Pop3), there were no pronounced differences between their accuracies and those of the highest ranking models for these traits.

For BRR, RKHS-KA, SM-GK and SVM, we also performed LOO evaluations (Table 1). The predictive performances were small even in a CV scenario with a larger training dataset. The highest observed R values were 0.38, 0.32 and 0.21 for SH in Pop1 (SM-GK and SVM), LDM in Pop2 (SVM), and SDM in Pop3 (SVM), respectively. Additionally, we did not observe clearly superior performance for any model, which would have allowed us to select a specific approach for fitting the regression model. For each trait, one of the four selected methods presented a slightly superior accuracy compared to that of the other three, as already observed in the 10-fold CV evaluations (Supplementary Figs. 11-13). Even with the best accuracy values observed, SH in Pop1, LDM in Pop2 and SDM in Pop3 could not be efficiently predicted, once again demonstrating the need for the development of suitable regression methods for complex polyploid grasses.

### Feature Selection

To improve predictive accuracies, we combined different FS methods for both classification (C) and regression (R) techniques. The regression problem was configured considering trait value prediction, and classification, considering the prediction of the clusters defined for each trait. To supply a more restricted group of markers, we used two different approaches to marker selection: the intersection between the three FS techniques used in classification (C3) or regression (R3) and the intersection of at least two out of the three FS methods (C2 and R2). Each approach generated a different quantity of markers when considering the regression and classification problems separately (Supplementary Table 2).

Regression-based FS generated larger quantities of markers, ranging from 898 to 980 in Pop1, from 242 to 367 in Pop2, and from 266 to 373 in Pop3. Classification-based FS, on the other hand, enabled the identification of more restricted marker subsets (from 208 to 211 in Pop1, from 86 to 117 in Pop2, and from 58 to 133 in Pop3). Interestingly, approximately half of the markers found for classification were also found for regression, showing the compatibility of these approaches. As expected, such FS methods enabled higher accuracies for ML prediction and the categorization of phenotype-associated genomic regions with diverse biological functions (Supplementary Figs. 14-41).

Even though the ML models for predicting the different traits differed in suitability, more pronounced differences were observed when changing the marker dataset and not the ML algorithm. For some trait-model combinations, the more effective strategy was employing C2/R2, and for others, C3/R3. Regarding the biological process GO terms associated with the genes found in these marker regions, there was a clear core set of common interactions for each trait when analyzing the regression- and classification-based approaches, with differences regarding the inclusion of novel connections and categories.

### Joint Learning

For each trait, we compared the predictive performances of the BRR, RKHS-KA and SM-GK models using the entire set of markers with the performances of SM-GK coupled with the FS approaches considering classification (C) and regression (R) algorithms. In addition to testing the intersection of the three FS techniques tested for both C and R (C3/R3) and the intersection of at least two techniques out of the three (C2/R2), we also evaluated (i) the union of C2 and R2 (CR2), (ii) the union of C3 and R3 (CR3), and (iii) the intersection of C2 and R2 (ICR2). Finally, we tested the addition of C3 and R3 as fixed effects (C3F and R3F) with the maintenance of the remaining markers for estimating the relationship matrix *K*. All these models were evaluated using R Pearson correlation coefficients together with MSE measures in the 10-fold CV scenario established; these measures were contrasted with Tukey’s multiple comparisons test (Supplementary Figs. 42-55).

The correlation values calculated when subsetting the marker datasets clearly increased due to decreases in MSE values. The addition of markers as fixed effects, on the other hand, increased the R Pearson coefficients together with the MSEs; this indicates that such an increase in the correlation is derived from higher variability in the dataset rather than from better predictive performance. By selecting the best results according to Tukey’s test for both evaluated metrics (Supplementary Table 3), we could establish the best models as SM-GK/R2 and SM-GK/CR2 for Pop1, SM-GK/R2, SM-GK/R3, and SM-GK/CR3 for Pop2, and SM-GK/R2, SM-GK/R3, SM-GK/CR2, and SM-GK/CR3 for Pop3.

Interestingly, we did not observe differences when applying SM-GK with R2 or CR2 and with R3 or CR3. In fact, although presenting intersections with R2 and R3 markers, C2 and C3 markers by themselves did not perform well in regression model estimation (Supplementary Figs. 42-55). To further analyze these results, we employed LOO CV with the SM-GK regression model and the GNB classification algorithm for marker datasets R2, R3, CR2 and CR3 (Supplementary Table 4). We selected the GNB model due to the promising results observed in previous analyses (Supplementary Figs. 14-41).

Similar to the results observed for the regression techniques, the classification algorithms performed better with the inclusion of markers selected for classification. However, as previously pointed out, the use of both strategies combined (CR2 and CR3) produced satisfactory results for both classification and regression, supplying a powerful set of markers to be considered for prediction (Fig. 2).

**Figure 2.**
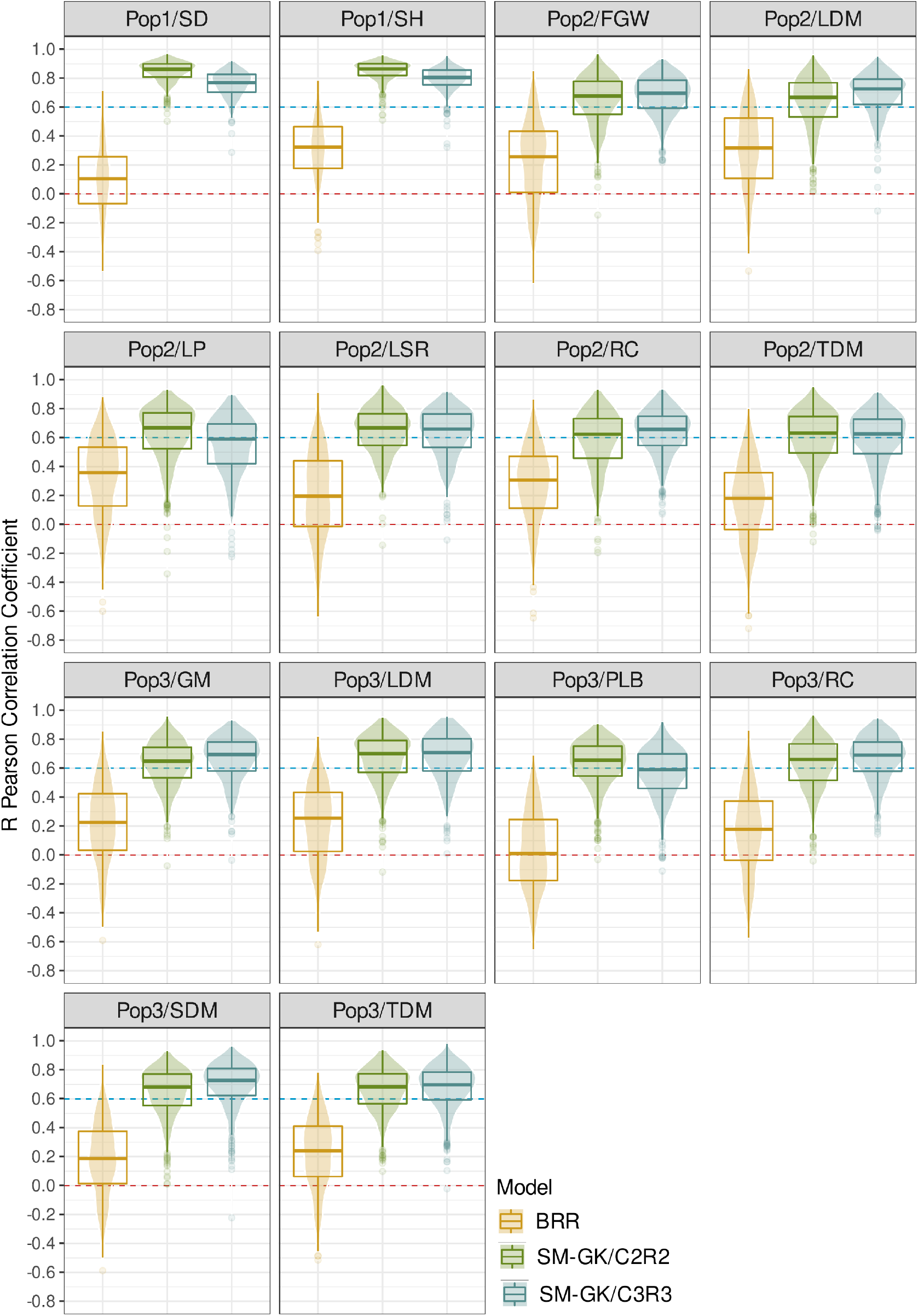
Genomic prediction models’ performances for the populations of (Pop1) sugarcane (stalk diameter (SD) and stalk height (SH)); (Pop2) *Urochloa decumbens* (field green weight (FGW), leaf dry matter (LDM), leaf percentage (LP), leaf stem ratio (LSR), regrowth capacity (RC), and total dry matter (TDM)); and (Pop3) *Megathyrsus maximus* (green matter (GM), LDM, percentage of leaf blade (PLB), RC, stem dry matter (SDM), and TDM). The Bayesian ridge regression (BRR) and single-environment main genotypic effect model with a Gaussian kernel (SM-GK) approaches with the inclusion of different feature selection methods (CR2 and CR3) were evaluated.

As the final approach suggested for constructing these GP models for complex polyploids, we highlight the approach of subsetting the marker dataset using the combined FS technique CR3, mainly because of (i) the high predictive accuracies observed for all traits, (ii) its ability to accommodate both classification and regression models, and (iii) the small number of markers compared to that in the CR2 dataset (for all traits, the number of markers selected for CR3 was approximately 10% of the number of markers selected for CR2). Although in some scenarios the prediction accuracies were higher for CR2, even when using CR3, they were significantly better than those observed with the traditional approaches.

Finally, by combining such an FS system based on the joint use of classification and regression ML strategies (CR3) with a popular statistical model for GP (SM-GK), we were able to suggest an ensemble method for predicting quantitative phenotypes in highly complex polyploid species. The increases in accuracy are remarkable (Fig. 3), and such a strategy also enables the construction of classification models for assisting in breeding selection strategies.

**Figure 3.**
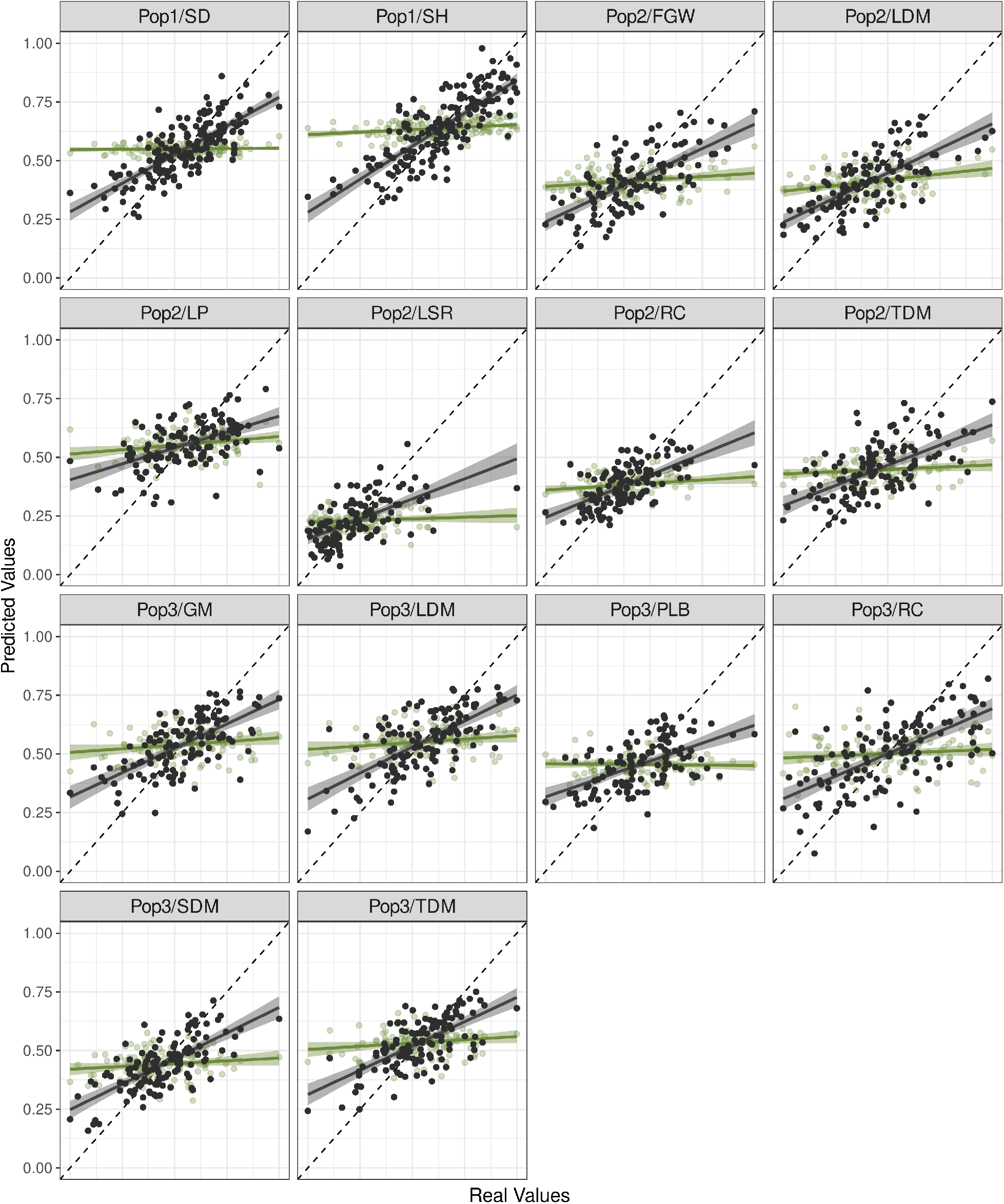
Correlation plots for the predicted and real values in the populations: (Pop1) sugarcane (stalk diameter (SD) and stalk height (SH)); (Pop2) *Urochloa decumbens* (field green weight (FGW), leaf dry matter (LDM), leaf percentage (LP), leaf stem ratio (LSR), regrowth capacity (RC), and total dry matter (TDM)); and (Pop3) *Megathyrsus maximus* (green matter (GM), LDM, percentage of leaf blade (PLB), RC, stem dry matter (SDM), and TDM). The approaches evaluated were Bayesian ridge regression (BRR), colored green, and the single-environment main genotypic effect model with a Gaussian kernel (SM-GK) with the inclusion of the established feature selection machinery, colored black.

### Simulation Approaches

To evaluate the feasibility of using the proposed approach for wider datasets, we performed simulations of a wheat breeding program and selected lines from the PYT for use in CV strategies. Using ten years of phenotypic selection, we were able to create 4 different datasets of 10,000 rows corresponding to genotypic data created based on 2, 10, 100 and 1,000 quantitative trait loci (QTLs). From these sets, different quantities of rows were sampled for independent evaluations (100, 250, 500, 1,000, 2,000, 5,000, and 10,000). We contrasted the performance of the SM-GK using the entire marker dataset with the performance considering marker selection with CR3 (Fig. 4). For all the configurations tested, the prediction accuracies of SM-GK/CR3 were better than those of SM-GK, with differences in the percentage of increase depending on the configuration established.

**Figure 4.**
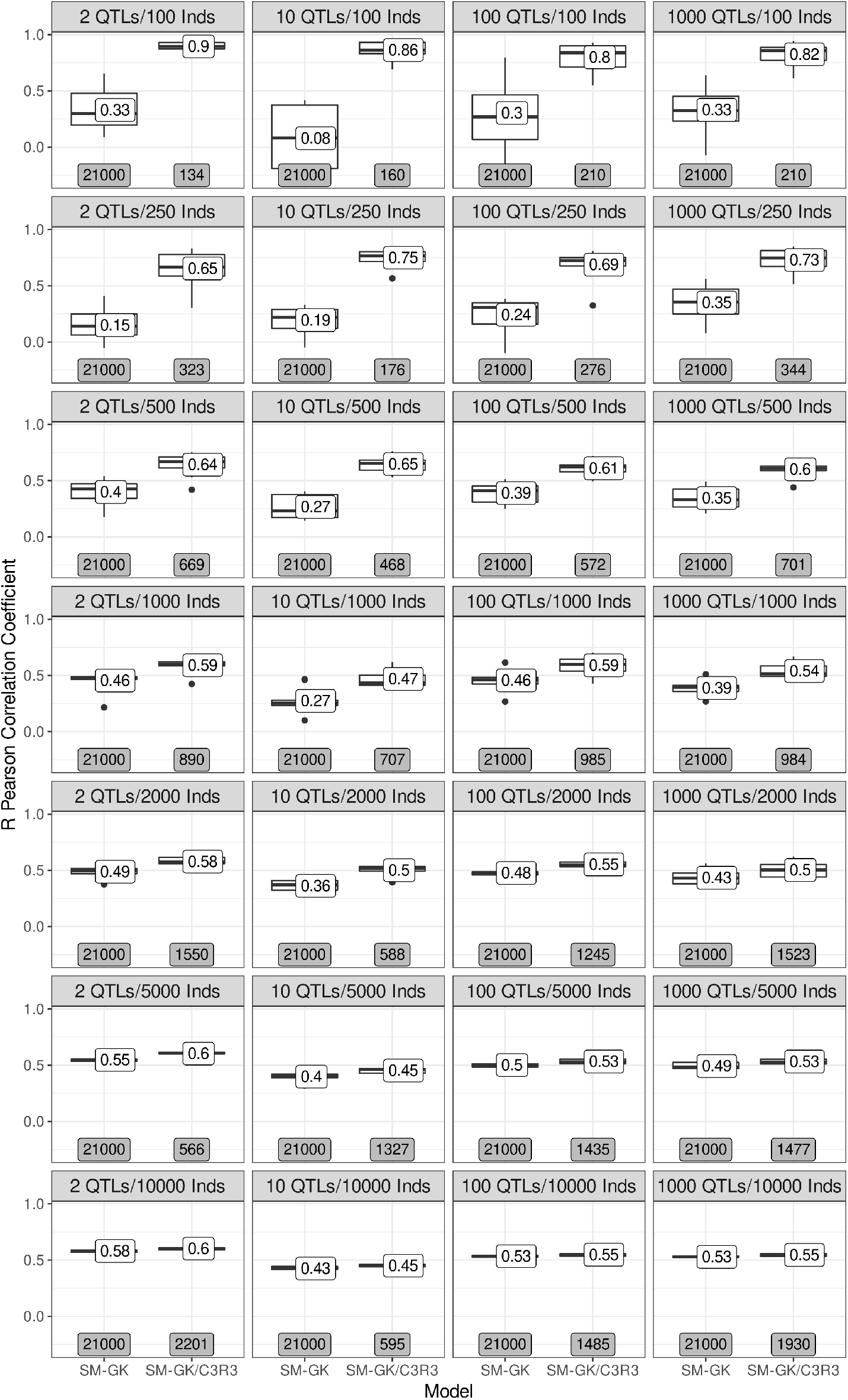
Accuracies of simulation scenarios created using a wheat breeding program and different quantities of QTLs per chromosome (2, 10, 100, and 1,000). From 20 years of phenotypic selection (10 years of burn-in), preliminary yield trials (1,000 lines) were selected by year, forming a dataset of 10,000 genotypes, which was sampled for different quantities of individuals (100, 250, 500, 1,000, 2,000, 5,000, and 10,000) to perform 10-fold cross-validation using a single-environment main genotypic effect model with a Gaussian kernel (SM-GK) with and without the inclusion of the established feature selection approach (SM-GK/C3R3). The values in the boxplot labels represent the mean accuracy, and in the gray boxes the quantity of markers used to create the predictive model.

Even though we observed slight differences in model performance when using different QTL quantities for the creation of genotypic datasets, the most pronounced increases in performance for SM-GK/CR3 compared with SM-GK were based on the quantity of individuals sampled from the data simulated and used for CV. In the smallest datasets (100 individuals), the accuracy for SM-GK/CR3 was approximately three times that of SM-GK, with a marker reduction exceeding 99%. The same increases in accuracy and decreases in marker data were also observed for 250 individuals. For 500 individuals, the increase was approximately 2 times, and the marker reduction was 98%. For more than 1,000 individuals, the reductions in the marker dataset were also large (approximately 95%, 95%, 94%, and 90% for 1,000, 2,000, 5,000, and 10,000 individuals, respectively); however, the increases in accuracy were more modest, being approximately 50%, 20%, 10%, and 5% for 1,000, 2,000, 5,000, and 10,000 individuals, respectively.

## Discussion

SU and FGs have considerable global economic importance, and their production demand is constantly rising^1,2^. In the current scenario of climate change and food insecurity, the development of novel and more productive cultivars by breeding programs represents the most effective and sustainable strategy for increasing production^83,84^. Although genomic approaches have revolutionized plant breeding with large associated genetic gains^85^, the inclusion of such techniques for the species employed in this work is still in its infancy^86,87^. The genomic complexity of these crops has hindered the development and acquisition of breeding genomic resources for many years^7,8^. In such a scenario, different molecular strategies have been tested for reducing such genomic complexity through methylation-sensitivity enzymes and facilitating the genotyping process while reducing associated costs, including GBS^43,88^. In this work, different populations genotyped with the GBS strategy were used to evaluate the estimation performance of GP models.

Even though GBS has been suitable for several studies of QTL mapping in these species^31,32,89^, developed GP strategies have presented low prediction accuracies^20,22,90,91^. Therefore, for practical incorporation of such techniques into breeding programs through GS, improved GP strategies are required, especially for most traits of interest, which are quantitative and highly polygenic and show phenotypic effects spread across unknown QTLs organized in a very complex genetic architecture^92^. Motivated by these facts, we suggested the incorporation of FS into GP as a means of circumventing the difficulty of estimating genotype and phenotype interactions and improving the low accuracies observed.

The traditional approaches employed for GP are based on the creation of regression models^25^, which estimate marker or genotype effects for the phenotype of interest, providing a means of predicting plant performances in the field^25^. In recent years, many studies have evaluated different types of statistical paradigms for such model estimation; however, most of the results show little/no changes in accuracy^93–95^. In contrast to the model efficiency observed for species with well-established genomic resources^96,97^, these strategies do not work properly in our scenario, inaccurately estimating the genetic effects of a specific trait and causing a significant loss of accuracy. We hypothesize that this deficiency is mainly caused by the difficulty of properly genotyping the individuals, which is due to several factors, including a lack of appropriate genomic references, the frequent occurrence of duplicated regions^7,13^, aneuploidy at different loci^98^, the difficulty of estimating correct allele dosages^99^, and the large amount of missing data^100^. Our first attempt to circumvent some of these limitations involved employing allele proportions instead of dosages, expanding the marker dataset and removing statistical assumptions regarding dosage attributions^27^.

The inclusion of ML approaches in GP has been controversial, being described as beneficial^95,101–103^ or not advantageous^18,104,105^. In the first scenarios that we evaluated in our work, we did not observe large increases when incorporating ML into prediction. Most of the ML algorithms performed worse than the traditional modeling approaches. Although we observed reasonable accuracies for some specific k-fold configurations (Supplementary Figs. 11-13), this was not uniform across the CV scenarios established, which corroborates the strong impact of genotyping results on model creation. Beneficial k-fold splits for prediction represent similar training and test datasets (i.e., individuals with similar polymorphic profiles), and in such cases, GP models work accurately. Therefore, we consider that, in addition to our hypothesis that the marker datasets employed were not accurate enough for creating such models, the lack of satisfactory accuracies might be related to an absence of markers linked to several trait-associated QTLs, probably due to the GBS protocol and bioinformatics procedures employed. To evaluate these factors, we decided to check whether the exclusion of nonbeneficial markers for prediction through FS could improve model performance based on (i) possible incorrect marker estimation, (ii) the principle that not all SNPs influence the phenotype^106^, and (iii) our observation that the models were not able to automatically separate the markers most important for prediction from the rest of the SNPs.

This strategy of FS has already demonstrated promising results across the literature^27,107–110^. Subsetting datasets through principles of FS is a common strategy in data science for datasets with a large number of variables^111^. Considering the principle underlying GS that all markers can be used in model construction with different effects estimated for the traits of interest^15^, FS methods have not been required in the construction of these predictive models. However, this is not the scenario for several complex datasets, including those used in this work. An FS system in ML aims at removing redundant, incorrect and irrelevant data from the dataset prior to prediction^112^, and for a GP problem, this strategy supplies a means of selecting the most promising set of markers for prediction, which might also be indicative of QTL associations^113^.

The reduction of markers through the different FS strategies employed was effective, dramatically reducing the datasets and increasing the prediction accuracies. For the final strategy considered, the intersection of the three FS methods selected enabled the establishment of a very restricted dataset with insights into biological functions potentially associated with different trait configurations. Interestingly, all of the traits were associated with genes having biological functions related to regulatory mechanisms (e.g., regulation of transcription, translation, and DNA duplication), which is in accordance with the control mechanism impacting protein-DNA interactions expected for QTLs^114^. Additionally, important categories could be retrieved for the evaluated agronomic traits, such as carbohydrate metabolism, defense response, cellulose biosynthetic process, and response to stress. All of these GO terms have been previously associated with important biological processes regulating plant growth performance^31,115,116^. Although a detailed analysis should be performed to provide in-depth inferences about these mechanisms underlying the trait configurations, this preliminary investigation highlights the potential of the methodology to reveal such important biological features associated with target traits for breeding programs.

Considering that the final purpose of breeding programs is selecting the most promising genotypes for advancing breeding cycles and producing novel cultivars, we also considered for each individual a related phenotypic group, supplying an indicator of its overall performance. Through the initial phenotypic clustering analyses performed (Supplementary Figs. 7-9), it was possible to observe distinct groups of individuals with similar phenotypic values, which were considered classes in an ML-based classification scenario. Such interval prediction can be considered a complementary strategy for selecting the best potential candidates for a specific trait because of its capability of excluding a large number of candidates that have fewer chances to be among the top genotypes in terms of performance. Such interval prediction follows the same principle of ranking approaches, which have already been demonstrated as a potential tool in selective breeding^117^. Different from traditional GP models based on regression, this classification task can take advantage of a vast set of ML algorithms. Together with the marker dataset selected through FS methods applied to the quantitative traits, we coupled an FS system based on classification, selecting the markers related to these defined intervals and forming a joint strategy for prediction. Therefore, in addition to creating a prediction model for GP of quantitative traits, the same marker dataset can be applied to an independent system estimated for interval prediction.

Regarding the performance comparison of the best models (BRR, RKHS-KA, SM-GK, and SVM), we did not observe significant differences between the accuracy results. For this reason, we selected only one of them, SM-GK, for combination with the FS system. In addition to clear improvements in accuracy by such SNP subsetting, we also evaluated whether such selection could serve as a means of defining fixed effects in GP models. Although we also observed increases in the measurements of accuracy through Pearson correlation coefficients, the same beneficial aspect was not observed when evaluating the models’ performance through MSEs. Instead, we observed higher predictive variability with stronger correlations and larger associated errors. Such a conclusion was previously reached when using peaks from a genome-wide association study (GWAS) as fixed effects in GP^118^. For this reason, we considered the subsetting strategy as the most promising method for boosting accuracies in our scenarios.

On the basis of the improvements observed (Fig. 3), we concluded that such an FS system was essential for showing the inefficiency of the models tested in terms of capturing real marker effects. Including a larger number of SNPs in GP may introduce background noise^108^, and in our scenario, such noise is expected to be found due to the elevated genomic complexity of these species. Interestingly, the GP models for SU presented better results than those for FGs. For SU SNP data, an SU pseudoreference genome was used for SNP calling^55^, different from the approach applied to FGs, where SNPs were estimated using genomic references from closely related species. Using *Sorghum bicolor* as a genomic reference for SU, a close relative, also decreases the quantity of reliable markers^27,110^ and, consecutively, the chances of finding SNPs surrounding QTL regions. For this reason, we believe that the increases in accuracy in FG results were more modest than those in SU.

As the last evaluations performed, we created simulated populations using a wheat breeding program to check the feasibility of the approach in scenarios with a broader set of individuals and not only single biparental populations. One of the main limitations that we found was related to the number of individuals used for performing k-fold validation. Increases in the predictive accuracies were modest when very large populations were analysed, but even in these cases our approach still represents an important strategy to reduce marker datasets needed for prediction. This might contribute with cutting down genotyping costs – a constraint that remains highly relevant in breeding programs of the species under study. Increasing the number of observations in the dataset and using the same quantity of folds for CV introduce a larger number of samples to be predicted, which may correspond to a more difficult prediction task if the individuals in the training and test sets are more distantly genetically related^119^. This fact corroborates the need to develop appropriate populations for the proper application of GS models^120^. Additionally, using a small number of individuals leads to a restricted set of markers linked to the phenotypic variation being examined, which is not expected for a larger number of genotypes^121^. As expected, the methodology worked efficiently for a broad number of QTLs, showing its appropriateness for different quantitative traits, as already demonstrated in this study.

In the present study, we provided a method for GP using different datasets of polyploid grasses. Combining an FS engineering system capable of retrieving markers important for classification and regression, our methodology also shows great potential for investigating marker-trait associations. Additionally, our results highlight the benefit of incorporating methodologies for marker selection into prediction, which may also be seen as a promising approach for developing targeted sequencing methodologies that can be applied to create models for GS. With a small number of markers, we could achieve high associated accuracies for quantitative traits and for predicting putative performance intervals through classification, which can be seen as a complementary breeding tool. This strategy has the potential to aid in the development of models for species with elevated genomic complexity, surpassing the limitations of available genomic resources and supplying a means of incorporating GS into their breeding programs.

## Supporting information

Supplementary Results

## Acknowledgements

The authors gratefully acknowledge the Fundação de Amparo à Pesquisa do Estado de São Paulo (FAPESP) for Ph.D. fellowships to AA (2019/03232-6) and RP (2019/21682-9), for a research internship abroad (BEPE) scholarship to AA (2019/26858-8), and for a PD scholarship to RF (2018/19219-6); the Coordenação de Aperfeiçoamento do Pessoal de Nível Superior (CAPES) for financial support (Computational Biology Program); and the Conselho Nacional de Desenvolvimento Científico e Tecnológico (CNPq).

## Author contribution statement

AA and RF performed all the analyses and wrote the manuscript; AM and RP assisted in the genotypic analyses and in manuscript writing; LL assisted in the phenotypic analyses; GG assisted in the simulation procedures; EC, LP, ML, MS, LK, SB, CV, and LC conducted the field experiments; AG, RK, AL, GG and AS conceived the project. All authors reviewed, read and approved the manuscript.

## Data Availability

Accession codes of sequencing data are available through the Sequence Read Archive (SRA) database with the accession numbers PRJNA478025, PRJNA472576 and PRJNA563938.

## References

1. Faostat, R. et al. Faostat database. Food Agric. Organ. UN (2017).

2. ISO. International sugar organization (2020).

3. Hoang, N. V., Furtado, A., Botha, F. C., Simmons, B. A. & Henry, R. J. Potential for genetic improvement of sugarcane as a source of biomass for biofuels. Front. bioengineering biotechnology 3, 182 (2015).

4. Jank, L., Barrios, S. C., do Valle, C. B., Simeão, R. M. & Alves, G. F. The value of improved pastures to brazilian beef production. Crop. Pasture Sci. 65, 1132–1137 (2014).

5. Prache, S., Martin, B. & Coppa, M. Authentication of grass-fed meat and dairy products from cattle and sheep. Animal 14, 854–863 (2020).

6. Pereira, J. F. et al. Research priorities for next-generation breeding of tropical forages in brazil. Crop. Breed. Appl. Biotechnol. 18, 314–319 (2018).

7. Thirugnanasambandam, P. P., Hoang, N. V. & Henry, R. J. The challenge of analyzing the sugarcane genome. Front. plant science 9, 616 (2018).

8. Schiessl, S.-V., Katche, E., Ihien, E., Chawla, H. S. & Mason, A. S. The role of genomic structural variation in the genetic improvement of polyploid crops. The Crop. J. 7, 127–140 (2019).

9. Zhang, M. et al. Analysis of the genes controlling three quantitative traits in three diverse plant species reveals the molecular basis of quantitative traits. Sci. reports 10, 1–14 (2020).

10. Comai, L. The advantages and disadvantages of being polyploid. Nat. reviews genetics 6, 836–846 (2005).

11. Fu, D., Mason, A. S., Xiao, M. & Yan, H. Effects of genome structure variation, homeologous genes and repetitive dna on polyploid crop research in the age of genomics. Plant Sci. 242, 37–46 (2016).

12. Bourke, P. M., Voorrips, R. E., Visser, R. G. & Maliepaard, C. Tools for genetic studies in experimental populations of polyploids. Front. plant science 9, 513 (2018).

13. Worthington, M. et al. A new brachiaria reference genome and its application in identifying genes associated with natural variation in tolerance to acidic soil conditions among brachiaria grasses. bioRxiv 843870 (2019).

14. Bhat, J. A. et al. Genomic selection in the era of next generation sequencing for complex traits in plant breeding. Front. genetics 7, 221 (2016).

15. Meuwissen, T. H., Hayes, B. J. & Goddard, M. E. Prediction of total genetic value using genome-wide dense marker maps. Genetics 157, 1819–1829 (2001).

16. Amadeu, R. R. et al. Impact of dominance effects on autotetraploid genomic prediction. Crop. Sci. 60, 656–665 (2020).

17. Juliana, P. et al. Improving grain yield, stress resilience and quality of bread wheat using large-scale genomics. Nat. genetics 51, 1530–1539 (2019).

18. Zingaretti, L. M. et al. Exploring deep learning for complex trait genomic prediction in polyploid outcrossing species. Front. plant science 11, 25 (2020).

19. Ferrão, L. F. V., Amadeu, R. R., Benevenuto, J., de Bem Oliveira, I. & Munoz, P. R. Genomic selection in an outcrossing autotetraploid fruit crop: lessons from blueberry breeding. Front. plant science 1075 (2021).

20. Batista, L. G., Mello, V. H., Souza, A. P. & Margarido, G. R. Genomic prediction with allele dosage information in highly polyploid species. Theor. Appl. Genet. 1–17 (2021).

21. Simeão Resende, R. M., Casler, M. D. & de Resende, M. D. V. Genomic selection in forage breeding: accuracy and methods. Crop. Sci. 54, 143–156 (2014).

22. de C. Lara, L. A. et al. Genomic selection with allele dosage in panicum maximum jacq. G3: Genes, Genomes, Genet. 9, 2463–2475 (2019).

23. Deomano, E. et al. Genomic prediction of sugar content and cane yield in sugar cane clones in different stages of selection in a breeding program, with and without pedigree information. Mol. Breed. 40, 1–12 (2020).

24. Lozada, D. N., Mason, R. E., Sarinelli, J. M. & Brown-Guedira, G. Accuracy of genomic selection for grain yield and agronomic traits in soft red winter wheat. BMC genetics 20, 1–12 (2019).

25. Azodi, C. B. et al. Benchmarking parametric and machine learning models for genomic prediction of complex traits. G3: Genes, Genomes, Genet. 9, 3691–3702 (2019).

26. Abdollahi-Arpanahi, R., Gianola, D. & Peñagaricano, F. Deep learning versus parametric and ensemble methods for genomic prediction of complex phenotypes. Genet. Sel. Evol. 52, 1–15 (2020).

27. Aono, A. H. et al. Machine learning approaches reveal genomic regions associated with sugarcane brown rust resistance. Sci. reports 10, 1–16 (2020).

28. CONSECANA-CONSELHO, D. P. D. C. & DE-AÇÚCAR, A. E. Á. D. Manual de instruções. CONSECANA-SP, Piracicaba,.

29. Simioni, C. & do Valle, C. B. Chromosome duplication in brachiaria (a. rich.) stapf allows intraspecific crosses. Crop. Breed. & Appl. Biotechnol. 9 (2009).

30. Mateus, R. G. et al. Genetic parameters and selection of brachiaria decumbens hybrids for agronomic traits and resistance to spittlebugs. Crop. Breed. Appl. Biotechnol. 15, 227–234 (2015).

31. Deo, T. G. et al. High-resolution linkage map with allele dosage allows the identification of regions governing complex traits and apospory in guinea grass (megathyrsus maximus). Front. plant science 11, 15 (2020).

32. Ferreira, R. C. U. et al. Genetic mapping with allele dosage information in tetraploid urochloa decumbens (stapf) rd webster reveals insights into spittlebug (notozulia entreriana berg) resistance. Front. plant science 10, 92 (2019).

33. Team, R. C. et al. R: A language and environment for statistical computing. (2013).

34. Munoz, F. & Rodriguez, L. S. breedr: Statistical methods for forest genetic resources analysis. In Trees for the future: plant material in a changing climate, 13–p (2014).

35. Peterson, R. A. bestnormalize: normalizing transformation functions. R package version 1, 573 (2018).

36. Butler, D., Cullis, B. R., Gilmour, A. & Gogel, B. Asreml-r reference manual. The State Queensland, Dep. Prim. Ind. Fish. Brisb. (2009).

37. Gilmour, A. R., Gogel, B. J., Cullis, B. R., Welham, S. & Thompson, R. Asreml user guide release 1.0. (2002).

38. Brien, C. asremlplus: Augments the use of asreml-r in fitting mixed models. R package version 2 (2016).

39. Box, G. E. & Cox, D. R. An analysis of transformations. J. Royal Stat. Soc. Ser. B (Methodological) 26, 211–243 (1964).

40. Wickham, H., Chang, W. & Wickham, M. H. Package ‘ggplot2’. Creat. Elegant Data Vis. Using Gramm. Graph. Version 2, 1–189 (2016).

41. Peterson, B. G. et al. Package ‘performanceanalytics’. R Team Coop. 3, 13–14 (2018).

42. Elshire, R. J. et al. A robust, simple genotyping-by-sequencing (gbs) approach for high diversity species. PloS one 6, e19379 (2011).

43. Poland, J. A. & Rife, T. W. Genotyping-by-sequencing for plant breeding and genetics. The Plant Genome 5 (2012).

44. Aljanabi, S. M., Forget, L. & Dookun, A. An improved and rapid protocol for the isolation of polysaccharide-and polyphenol-free sugarcane dna. Plant Mol. Biol. Report. 17, 281–282 (1999).

45. Doyle, J. J. & Doyle, J. L. A rapid dna isolation procedure for small quantities of fresh leaf tissue. Tech. Rep. (1987).

46. Glaubitz, J. C. et al. Tassel-gbs: a high capacity genotyping by sequencing analysis pipeline. PloS one 9, e90346 (2014).

47. Pereira, G. S., Garcia, A. A. F. & Margarido, G. R. A fully automated pipeline for quantitative genotype calling from next generation sequencing data in autopolyploids. BMC bioinformatics 19, 1–10 (2018).

48. Li, H. et al. The sequence alignment/map format and samtools. Bioinformatics 25, 2078–2079 (2009).

49. Catchen, J., Hohenlohe, P. A., Bassham, S., Amores, A. & Cresko, W. A. Stacks: an analysis tool set for population genomics. Mol. ecology 22, 3124–3140 (2013).

50. McKenna, A. et al. The genome analysis toolkit: a mapreduce framework for analyzing next-generation dna sequencing data. Genome research 20, 1297–1303 (2010).

51. Garrison, E. & Marth, G. Haplotype-based variant detection from short-read sequencing. arXiv preprint 1207.3907 (2012).

52. Gordon, A., Hannon, G. et al. Fastx-toolkit. FASTQ/A short-reads preprocessing tools (unpublished) 5 (2010).

53. Altschul, S. F., Gish, W., Miller, W., Myers, E. W. & Lipman, D. J. Basic local alignment search tool. J. molecular biology 215, 403–410 (1990).

54. Li, H. Aligning sequence reads, clone sequences and assembly contigs with bwa-mem. arXiv preprint 1303.3997 (2013).

55. Grativol, C. et al. Sugarcane genome sequencing by methylation filtration provides tools for genomic research in the genus s accharum. The Plant J. 79, 162–172 (2014).

56. Langmead, B., Trapnell, C., Pop, M. & Salzberg, S. L. Ultrafast and memory-efficient alignment of short dna sequences to the human genome. Genome biology 10, 1–10 (2009).

57. Goodstein, D. M. et al. Phytozome: a comparative platform for green plant genomics. Nucleic acids research 40, D1178–D1186 (2012).

58. Van der Maaten, L. & Hinton, G. Visualizing data using t-sne. J. machine learning research 9 (2008).

59. Van Den Wollenberg, A. L. Redundancy analysis an alternative for canonical correlation analysis. Psychometrika 42, 207–219 (1977).

60. Krijthe, J., van der Maaten, L. & Krijthe, M. J. Package ‘rtsne’ (2018).

61. Oksanen, J. et al. Package ‘vegan’. Community ecology package, version 2, 1–295 (2013).

62. Charrad, M., Ghazzali, N., Boiteau, V. & Niknafs, A. Nbclust: an r package for determining the relevant number of clusters in a data set. J. statistical software 61, 1–36 (2014).

63. Yu, G., Smith, D. K., Zhu, H., Guan, Y. & Lam, T. T.-Y. ggtree: an r package for visualization and annotation of phylogenetic trees with their covariates and other associated data. Methods Ecol. Evol. 8, 28–36 (2017).

64. Gianola, D. Priors in whole-genome regression: the bayesian alphabet returns. Genetics 194, 573–596 (2013).

65. Gianola, D. & Van Kaam, J. B. Reproducing kernel hilbert spaces regression methods for genomic assisted prediction of quantitative traits. Genetics 178, 2289–2303 (2008).

66. Granato, I. et al. Bgge: a new package for genomic-enabled prediction incorporating genotype× environment interaction models. G3: Genes, Genomes, Genet. 8, 3039–3047 (2018).

67. Pérez, P. & de Los Campos, G. Genome-wide regression and prediction with the bglr statistical package. Genetics 198, 483–495 (2014).

68. Cover, T. & Hart, P. Nearest neighbor pattern classification. IEEE transactions on information theory 13, 21–27 (1967).

69. Cristianini, N., Shawe-Taylor, J. et al. An introduction to support vector machines and other kernel-based learning methods (Cambridge university press, 2000).

70. Breiman, L. Random forests. Mach. learning 45, 5–32 (2001).

71. Freund, Y. & Schapire, R. E. A decision-theoretic generalization of on-line learning and an application to boosting. J. computer system sciences 55, 119–139 (1997).

72. Pedregosa, F. et al. Scikit-learn: Machine learning in python. J. machine Learn. research 12, 2825–2830 (2011).

73. de Mendiburu, F. & de Mendiburu, M. F. Package ‘agricolae’. R Packag. Version 1–2 (2019).

74. Popescu, M.-C., Balas, V. E., Perescu-Popescu, L. & Mastorakis, N. Multilayer perceptron and neural networks. WSEAS Transactions on Circuits Syst. 8, 579–588 (2009).

75. Friedman, N., Geiger, D. & Goldszmidt, M. Bayesian network classifiers. Mach. learning 29, 131–163 (1997).

76. Chen, T. & Guestrin, C. Xgboost: A scalable tree boosting system. In Proceedings of the 22nd acm sigkdd international conference on knowledge discovery and data mining, 785–794 (2016).

77. Supek, F., Bošnjak, M., Škunca, N. & Šmuc, T. Revigo summarizes and visualizes long lists of gene ontology terms. PloS one 6, e21800 (2011).

78. Shannon, P. et al. Cytoscape: a software environment for integrated models of biomolecular interaction networks. Genome research 13, 2498–2504 (2003).

79. Gaynor, R. C., Gorjanc, G. & Hickey, J. M. Alphasimr: an r package for breeding program simulations. G3 11, jkaa017 (2021).

80. Gaynor, R. C. et al. A two-part strategy for using genomic selection to develop inbred lines. Crop. Sci. 57, 2372–2386 (2017).

81. de C Lara, L. A., Pocrnic, I., de P Oliveira, T., Gaynor, R. C. & Gorjanc, G. Temporal and genomic analysis of additive genetic variance in breeding programmes. Heredity (2021).

82. Martins, F. B. et al. A semi-automated snp-based approach for contaminant identification in biparental polyploid populations of tropical forage grasses. Front. plant science 12 (2021).

83. Lenaerts, B., Collard, B. C. & Demont, M. Improving global food security through accelerated plant breeding. Plant Sci. 287, 110207 (2019).

84. Qaim, M. Role of new plant breeding technologies for food security and sustainable agricultural development. Appl. Econ. Perspectives Policy 42, 129–150 (2020).

85. Poland, J. Breeding-assisted genomics. Curr. opinion plant biology 24, 119–124 (2015).

86. Yadav, S. et al. Accelerating genetic gain in sugarcane breeding using genomic selection. Agronomy 10, 585 (2020).

87. Simeão, R. M. et al. Genomic selection in tropical forage grasses: Current status and future applications. Front. Plant Sci. 12, 761 (2021).

88. Scheben, A., Batley, J. & Edwards, D. Genotyping-by-sequencing approaches to characterize crop genomes: choosing the right tool for the right application. Plant biotechnology journal 15, 149–161 (2017).

89. Balsalobre, T. W. A. et al. Gbs-based single dosage markers for linkage and qtl mapping allow gene mining for yield-related traits in sugarcane. BMC genomics 18, 1–19 (2017).

90. Matias, F. I. et al. On the accuracy of genomic prediction models considering multi-trait and allele dosage in urochloa spp. interspecific tetraploid hybrids. Mol. Breed. 39, 1–16 (2019).

91. Hayes, B. J. et al. Accuracy of genomic prediction of complex traits in sugarcane. Theor. Appl. Genet. 134, 1455–1462 (2021).

92. Mauricio, R. Mapping quantitative trait loci in plants: uses and caveats for evolutionary biology. Nat. Rev. Genet. 2, 370–381 (2001).

93. Roorkiwal, M. et al. Genome-enabled prediction models for yield related traits in chickpea. Front. plant science 7, 1666 (2016).

94. Varshney, R. K. Exciting journey of 10 years from genomes to fields and markets: some success stories of genomics-assisted breeding in chickpea, pigeonpea and groundnut. Plant Sci. 242, 98–107 (2016).

95. Ma, W. et al. A deep convolutional neural network approach for predicting phenotypes from genotypes. Planta 248, 1307–1318 (2018).

96. Crossa, J. et al. Genomic prediction in cimmyt maize and wheat breeding programs. Heredity 112, 48–60 (2014).

97. Millet, E. J. et al. Genomic prediction of maize yield across european environmental conditions. Nat. genetics 51, 952–956 (2019).

98. Sforça, D. A. et al. Gene duplication in the sugarcane genome: a case study of allele interactions and evolutionary patterns in two genic regions. Front. plant science 10, 553 (2019).

99. Garcia, A. A. et al. Snp genotyping allows an in-depth characterisation of the genome of sugarcane and other complex autopolyploids. Sci. reports 3, 1–10 (2013).

100. Torkamaneh, D., Laroche, J. & Belzile, F. Genome-wide snp calling from genotyping by sequencing (gbs) data: a comparison of seven pipelines and two sequencing technologies. PloS one 11, e0161333 (2016).

101. Bellot, P., de los Campos, G. & Pérez-Enciso, M. Can deep learning improve genomic prediction of complex human traits? Genetics 210, 809–819 (2018).

102. Waldmann, P., Pfeiffer, C. & Mészáros, G. Sparse convolutional neural networks for genome-wide prediction. Front. Genet. 11 (2020).

103. Liu, Y. et al. Phenotype prediction and genome-wide association study using deep convolutional neural network of soybean. Front. Genet. 10, 1091 (2019).

104. Montesinos-López, O. A. et al. Multi-trait, multi-environment genomic prediction of durum wheat with genomic best linear unbiased predictor and deep learning methods. Front. Plant Sci. 10 (2019).

105. Crossa, J. et al. Deep kernel and deep learning for genome-based prediction of single traits in multienvironment breeding trials. Front. Genet. 10 (2019).

106. Orgogozo, V., Morizot, B. & Martin, A. The differential view of genotype–phenotype relationships. Front. genetics 6, 179 (2015).

107. Bermingham, M. L. et al. Application of high-dimensional feature selection: evaluation for genomic prediction in man. Sci. reports 5, 1–12 (2015).

108. Li, B. et al. Genomic prediction of breeding values using a subset of snps identified by three machine learning methods. Front. genetics 9, 237 (2018).

109. Luo, Z., Yu, Y., Xiang, J. & Li, F. Genomic selection using a subset of snps identified by genome-wide association analysis for disease resistance traits in aquaculture species. Aquaculture 539, 736620 (2021).

110. Pimenta, R. J. G. et al. Genome-wide approaches for the identification of markers and genes associated with sugarcane yellow leaf virus resistance. Sci. Reports 11, 1–18 (2021).

111. Miao, J. & Niu, L. A survey on feature selection. Procedia Comput. Sci. 91, 919–926 (2016).

112. Cai, J., Luo, J., Wang, S. & Yang, S. Feature selection in machine learning: A new perspective. Neurocomputing 300, 70–79 (2018).

113. Jeong, S., Kim, J.-Y. & Kim, N. Gmstool: Gwas-based marker selection tool for genomic prediction from genomic data. Sci. reports 10, 1–12 (2020).

114. Gaffney, D. J. et al. Dissecting the regulatory architecture of gene expression qtls. Genome biology 13, 1–15 (2012).

115. Kasirajan, L., Hoang, N. V., Furtado, A., Botha, F. C. & Henry, R. J. Transcriptome analysis highlights key differentially expressed genes involved in cellulose and lignin biosynthesis of sugarcane genotypes varying in fiber content. Sci. reports 8, 1–16 (2018).

116. Volaire, F. et al. The resilience of perennial grasses under two climate scenarios is correlated with carbohydrate metabolism in meristems. J. experimental botany 71, 370–385 (2020).

117. Blondel, M., Onogi, A., Iwata, H. & Ueda, N. A ranking approach to genomic selection. PloS one 10, e0128570 (2015).

118. Rice, B. & Lipka, A. E. Evaluation of rr-blup genomic selection models that incorporate peak genome-wide association study signals in maize and sorghum. The Plant Genome 12 (2019).

119. Berro, I., Lado, B., Nalin, R. S., Quincke, M. & Gutiérrez, L. Training population optimization for genomic selection. The Plant Genome 12, 190028 (2019).

120. Isidro, J. et al. Training set optimization under population structure in genomic selection. Theor. applied genetics 128, 145–158 (2015).

121. Larkin, D. L., Lozada, D. N. & Mason, R. E. Genomic selection—considerations for successful implementation in wheat breeding programs. Agronomy 9, 479 (2019).

