## Supplementary Results for "A joint learning approach for genomic prediction in polyploid grasses"

**SUPPLEMENTARY TABLES**

**Supplementary Table S1**. Best strategies in terms of predictive performance as determined using a Tukey multiple comparisons test (p value of 0.05): Bayesian ridge regression (BRR), Bayesian reproducing kernel Hilbert spaces regression with kernel averaging (RKHS-KA), the single-environment main genotypic effect model with a Gaussian kernel (SM-GK), the k-nearest neighbors (KNN) algorithm, the support vector machine (SVM) strategy, AdaBoost, and random forest (RF).

| **Population** | **Trait** | **Best Models** |
|---|---|---|
| Sugarcane | Stalk diameter | BRR, RKHS-KA, and SVM |
| | Stalk height | SM-GK and SVM |
| *Urochloa decumbens* | Regrowth capacity | BRR, RKHS-KA, and SM-GK |
| | Field green weight | BRR, RKHS-KA, SM-GK, and SVM |
| | Total dry matter | BRR, RKHS-KA, SM-GK, and SVM |
| | Leaf dry matter | BRR, RKHS-KA, SM-GK, and SVM |
| | Leaf percentage | BRR, RKHS-KA, SM-GK, and SVM |
| | Leaf stem ratio | BRR, RKHS-KA, SM-GK, and SVM |
| *Megathyrsus maximus* | Green matter | BRR, RKHS-KA, SM-GK, and SVM |
| | Total dry matter | BRR, RKHS-KA, SM-GK, RF and SVM |
| | Leaf dry matter | BRR, RKHS-KA, SM-GK, and SVM |
| | Regrowth capacity | All |

| | | |
|---|---|---|
| | Stem dry matter | BRR, RKHS-KA, SM-GK, and SVM |
| | Percentage of leaf blade | RKHS-KA, KNN and RF |

**Supplementary Table S2.** SNPs selected through FS across the different traits separated according to the populations: (Pop1) cross between sugarcane commercial varieties, (Pop2) cross between genotypes of *Urochloa decumbens*, and (Pop3) cross between genotypes of *Megathyrsus maximus*. For marker selection, we employed classification (C)- and regression (R)-based approaches, selecting the intersection of the results from the three FS strategies tested (C3/I3) and the intersection of at least two out of the three FS methods (C2/I2). We also evaluated the intersection (ICR2/ICR3) and the union (CR2/CR3) between R and C in both approaches.

| | **Trait** | **C2** | **R2** | **ICR2** | **CR2** | **C3** | **R3** | **ICR3** | **CR3** |
|---|---|---|---|---|---|---|---|---|---|
| Pop1 | Stem height (SH) | 211 | 980 | 112 | 1079 | 14 | 82 | 4 | 92 |
| | Stem diameter (SD) | 208 | 898 | 70 | 1036 | 22 | 94 | 3 | 113 |
| | **Mean value** | **210** | **939** | **91** | **1058** | **18** | **88** | **4** | **103** |
| Pop2 | Regrowth capacity (RC) | 117 | 284 | 46 | 355 | 8 | 41 | 1 | 48 |
| | Field green weight (FGW) | 104 | 294 | 47 | 351 | 11 | 36 | 1 | 46 |
| | Total dry matter (TDM) | 86 | 281 | 31 | 336 | 9 | 40 | 0 | 49 |
| | Leaf dry matter (LDM) | 93 | 367 | 43 | 417 | 11 | 47 | 4 | 54 |
| | Leaf percentage (LP) | 113 | 289 | 55 | 347 | 12 | 48 | 2 | 58 |
| | Leaf stem ratio (LSR) | 115 | 242 | 59 | 298 | 16 | 39 | 5 | 50 |
| | **Mean value** | **105** | **293** | **47** | **351** | **11** | **42** | **2** | **51** |
| Pop3 | Green matter (GM) | 83 | 373 | 38 | 418 | 7 | 47 | 2 | 52 |
| | TDM | 133 | 362 | 56 | 439 | 26 | 40 | 3 | 63 |
| | LDM | 82 | 349 | 40 | 391 | 15 | 47 | 2 | 60 |
| | RC | 110 | 310 | 51 | 369 | 13 | 45 | 1 | 57 |
| | Stem dry matter (SDM) | 58 | 345 | 34 | 369 | 11 | 40 | 3 | 48 |

| | | | | | | | | | |
|---|---|---|---|---|---|---|---|---|---|
| | Percentage of leaf blade (PLB) | 115 | 266 | 43 | 338 | 21 | 23 | 7 | 37 |
| | **Mean value** | **97** | **334** | **44** | **387** | **16** | **40** | **3** | **53** |

**Supplementary Table S3.** Best models selected for the traits of the populations: (Pop1) cross between sugarcane commercial varieties, (Pop2) cross between genotypes of *Urochloa decumbens*, and (Pop3) cross between genotypes of *Megathyrsus maximus*. Prediction performances were evaluated as R Pearson correlation coefficients and mean squared errors (MSE) using the approaches employed for regression (Bayesian ridge regression (BRR), reproducing kernel Hilbert space with kernel averaging (RKHS-KA) model, and single-environment, main genotypic effect model with a Gaussian kernel (SM-GK)) and the markers selected through FS methods considering classification (C) and regression (R) algorithms and the intersection of the three tested techniques (C3/R3) or at least two of them (C2/R2). The combinations (i) union of C2 and R2 (CR2), (ii) union of C3 and R3 (CR3), and (iii) intersection of C2 and R2 (ICR2) were also evaluated, as well as the addition of C3 and R3 as fixed effects (C3F and R3F).

| | **Trait** | **Best Performance (R)** | **Best Performance (MSE)** | **Best Performance (R and MSE)** |
|---|---|---|---|---|
| Pop1 | Stem diameter (SD) | SM-GK/R2,<br>SM-GK/CR2 | SM-GK/R2,<br>SM-GK/CR2 | SM-GK/R2,<br>SM-GK/CR2 |
| | Stem height (SH) | SM-GK/R2,<br>SM-GK/CR2 | SM-GK/R2,<br>SM-GK/CR2 | SM-GK/R2,<br>SM-GK/CR2 |
| Pop2 | Field green weight (FGW) | SM-GK/R2,<br>SM-GK/R3,<br>SM-GK/CR2,<br>SM-GK/CR3,<br>SM-GK/ICR2 | SM-GK/R2,<br>SM-GK/R3,<br>SM-GK/CR3,<br>SM-GK/ICR2 | SM-GK/R2,<br>SM-GK/R3,<br>SM-GK/CR3,<br>SM-GK/ICR2 |
| | Leaf dry matter (LDM) | SM-GK/R3,<br>SM-GK/CR3 | SM-GK/R3,<br>SM-GK/CR3 | SM-GK/R3,<br>SM-GK/CR3 |
| | Leaf percentage (LP) | SM-GK/R2,<br>SM-GK/CR2 | SM-GK/R2,<br>SM-GK/CR2,<br>SM-GK/ICR2 | SM-GK/R2,<br>SM-GK/CR2 |
| | Leaf stem ratio (LSR) | SM-GK/R2,<br>SM-GK/R3, | SM-GK/R2,<br>SM-GK/R3, | SM-GK/R2,<br>SM-GK/R3, |

| | | | | |
|---|---|---|---|---|
| | | SM-GK/CR2, SM-GK/CR3, SM-GK/ICR2 | SM-GK/CR2, SM-GK/CR3, SM-GK/ICR2 | SM-GK/CR2, SM-GK/CR3, SM-GK/ICR2 |
| | Regrowth capacity (RC) | SM-GK/R3, SM-GK/CR3 | SM-GK/C2, SM-GK/R2, SM-GK/R3, SM-GK/CR2, SM-GK/CR3, SM-GK/ICR2, SM-GK/C3F | SM-GK/R3, SM-GK/CR3 |
| | Total dry matter (TDM) | SM-GK/R2, SM-GK/CR2, SM-GK/CR3 | SM-GK/R2, SM-GK/CR2, SM-GK/CR3 | SM-GK/R2, SM-GK/CR2, SM-GK/CR3 |
| Pop3 | Green matter (GM) | SM-GK/R2, SM-GK/R3, SM-GK/CR2, SM-GK/CR3 | SM-GK/R2, SM-GK/R3, SM-GK/CR2, SM-GK/CR3 | SM-GK/R2, SM-GK/R3, SM-GK/CR2, SM-GK/CR3 |
| | LDM | SM-GK/R2, SM-GK/R3, SM-GK/CR2, SM-GK/CR3 | SM-GK/R2, SM-GK/R3, SM-GK/CR2, SM-GK/CR3 | SM-GK/R2, SM-GK/R3, SM-GK/CR2, SM-GK/CR3 |
| | Percentage of leaf blade (PLB) | SM-GK/R2, SM-GK/R3, SM-GK/CR2 | SM-GK/R2, SM-GK/R3, SM-GK/CR2, SM-GK/CR3 | SM-GK/R2, SM-GK/R3, SM-GK/CR2 |
| | RC | SM-GK/R2, SM-GK/R3, SM-GK/CR2, SM-GK/CR3, SM-GK/ICR2 | All models except SM-GK/R3F | SM-GK/R2, SM-GK/R3, SM-GK/CR2, SM-GK/CR3, SM-GK/ICR2 |
| | Stem dry matter (SDM) | SM-GK/R3, SM-GK/CR3 | SM-GK/R3, SM-GK/CR3 | SM-GK/R3, SM-GK/CR3 |
| | TDM | SM-GK/R2, SM-GK/R3, SM-GK/CR2, SM-GK/CR3, SM-GK/ICR2 | All models SM-GK/C3F | SM-GK/R2, SM-GK/R3, SM-GK/CR2, SM-GK/CR3, SM-GK/ICR2 |

**Supplementary Table S4.** Leave-one-out evaluation for the traits of the populations: (Pop1) cross between sugarcane commercial varieties, (Pop2) cross between genotypes of *Urochloa decumbens*, and (Pop3) cross between genotypes of *Megathyrsus maximus*. Prediction performances were evaluated as R Pearson correlation coefficients using the single-environment, main genotypic effect model with a Gaussian kernel (SM-GK) and with the rate of the correctly classified genotypes according to group configuration using the Gaussian naive Bayes (GNB) algorithm. The markers were selected through FS methods considering classification (C) and regression (R) algorithms and the intersection of the three tested techniques (C3/R3) or at least two of them (C2/R2).

| | **Trait** | **Markers** | **R Pearson Correlation (SM-GK Regression)** | **Prediction Accuracy (GNB Classification)** | **Marker Quantity** |
|---|---|---|---|---|---|
| Pop1 | Stem diameter (SD) | R2 | **0.85** | 0.65 | 898 |
| | | CR2 | **0.85** | **0.74** | 1036 |
| | | R3 | 0.78 | 0.56 | 94 |
| | | CR3 | 0.75 | 0.67 | 113 |
| | Stem height (SH) | R2 | **0.85** | 0.68 | 980 |
| | | CR2 | **0.85** | **0.75** | 1079 |
| | | R3 | 0.80 | 0.67 | 82 |
| | | CR3 | 0.79 | 0.73 | 92 |
| Pop2 | Field green weight (FGW) | R2 | **0.69** | 0.64 | 294 |
| | | CR2 | 0.65 | **0.73** | 351 |
| | | R3 | 0.67 | 0.59 | 36 |
| | | CR3 | 0.67 | 0.57 | 46 |

| | | | | | |
|---|---|---|---|---|---|
| | Leaf dry matter (LDM) | R2 | 0.62 | 0.56 | 367 |
| | | CR2 | 0.63 | **0.66** | 417 |
| | | R3 | 0.68 | 0.64 | 47 |
| | | CR3 | **0.69** | **0.66** | 54 |
| | Leaf percentage (LP) | R2 | **0.63** | 0.57 | 289 |
| | | CR2 | 0.59 | **0.66** | 347 |
| | | R3 | 0.53 | 0.55 | 48 |
| | | CR3 | 0.51 | 0.62 | 58 |
| | Leaf stem ratio (LSR) | R2 | **0.63** | 0.66 | 242 |
| | | CR2 | 0.61 | 0.71 | 298 |
| | | R3 | **0.63** | 0.66 | 39 |
| | | CR3 | 0.59 | **0.76** | 50 |
| | Regrowth capacity (RC) | R2 | 0.56 | 0.62 | 284 |
| | | CR2 | 0.54 | **0.75** | 355 |
| | | R3 | **0.62** | 0.65 | 41 |
| | | CR3 | 0.61 | 0.69 | 48 |
| | Total dry matter (TDM) | R2 | **0.63** | 0.56 | 281 |
| | | CR2 | 0.60 | **0.72** | 336 |
| | | R3 | 0.53 | 0.58 | 40 |
| | | CR3 | 0.58 | 0.65 | 49 |
| Pop3 | Green matter (GM) | R2 | 0.62 | 0.70 | 373 |
| | | CR2 | 0.62 | **0.78** | 418 |
| | | R3 | 0.63 | 0.73 | 47 |
| | | CR3 | **0.65** | **0.78** | 52 |
| | LDM | R2 | 0.64 | 0.54 | 349 |

| | | | | | |
|---|---|---|---|---|---|
| | | CR2 | 0.65 | 0.59 | 391 |
| | | R3 | 0.65 | 0.53 | 47 |
| | | CR3 | **0.68** | **0.66** | 60 |
| | Percentage of leaf blade (PLB) | R2 | **0.61** | 0.62 | 266 |
| | | CR2 | 0.60 | **0.73** | 338 |
| | | R3 | 0.58 | 0.63 | 23 |
| | | CR3 | 0.55 | 0.67 | 37 |
| | RC | R2 | 0.63 | 0.39 | 310 |
| | | CR2 | 0.61 | 0.49 | 369 |
| | | R3 | **0.64** | 0.49 | 45 |
| | | CR3 | **0.64** | **0.59** | 57 |
| | Stem dry matter (SDM) | R2 | 0.63 | 0.68 | 345 |
| | | CR2 | 0.64 | 0.70 | 369 |
| | | R3 | 0.68 | 0.76 | 40 |
| | | CR3 | **0.69** | **0.80** | 48 |
| | TDM | R2 | 0.65 | 0.63 | 362 |
| | | CR2 | 0.65 | 0.72 | 439 |
| | | R3 | 0.66 | 0.65 | 40 |
| | | CR3 | **0.67** | **0.76** | 63 |

### SUPPLEMENTARY FIGURES

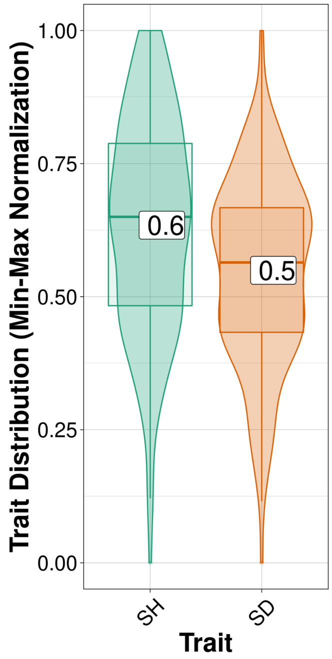


**Supplementary Fig. S1.** Best linear unbiased predictor (BLUP) distribution for individuals from the cross performed between sugarcane commercial varieties (values rescaled between 0 and 1). The evaluated traits were stalk height (SH) and stalk diameter (SD).

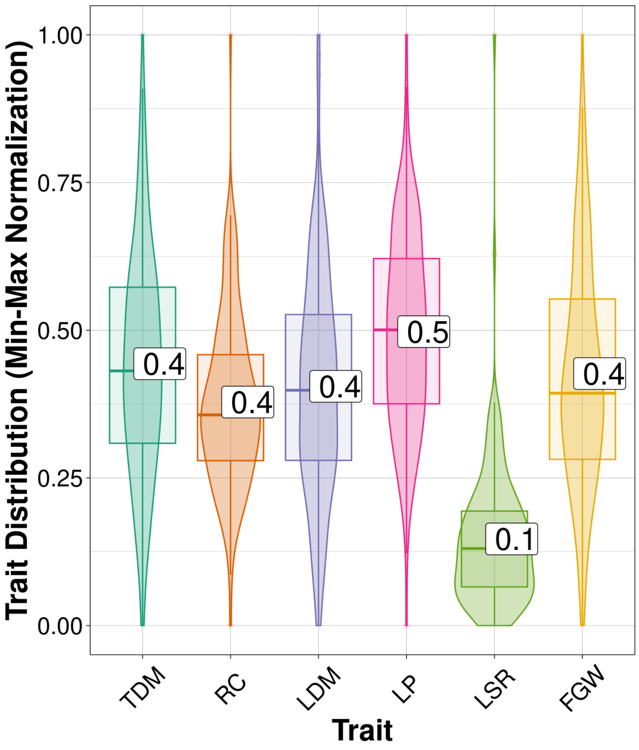


**Supplementary Fig. S2.** Corrected trait distribution for individuals from the cross performed between tetraploid genotypes of *Urochloa decumbens* (values rescaled between 0 and 1). The evaluated traits were total dry matter (TDM), regrowth capacity (RC), leaf dry matter (LDM), leaf percentage (LP), leaf stem ratio (LSR) and field green weight (FGW).

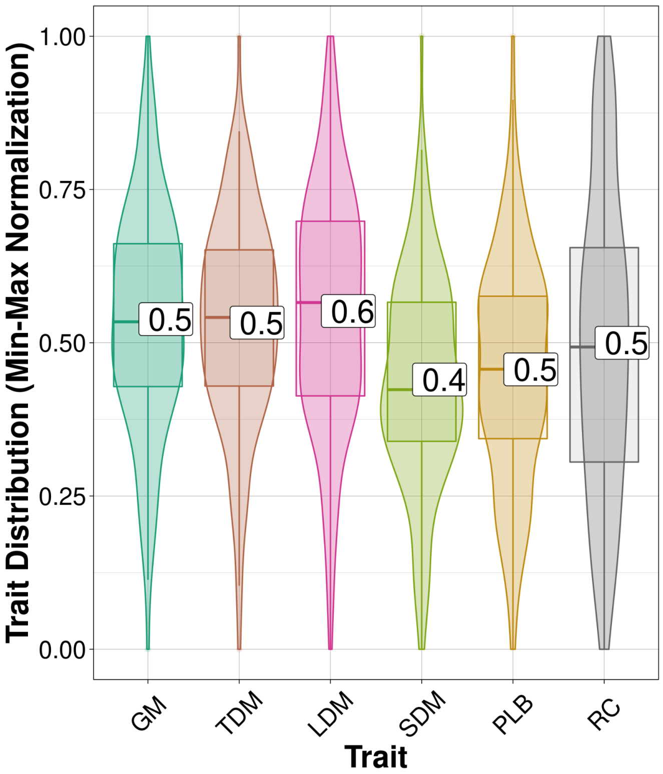


**Supplementary Fig. S3.** Corrected trait distribution for individuals from the cross between genotypes of *Megathyrsus maximus* (values rescaled between 0 and 1). The evaluated traits were green matter (GM), total dry matter (TDM), leaf dry matter (LDM), stem dry matter (SDM), percentage of leaf blade (PLB) and regrowth capacity (RC).

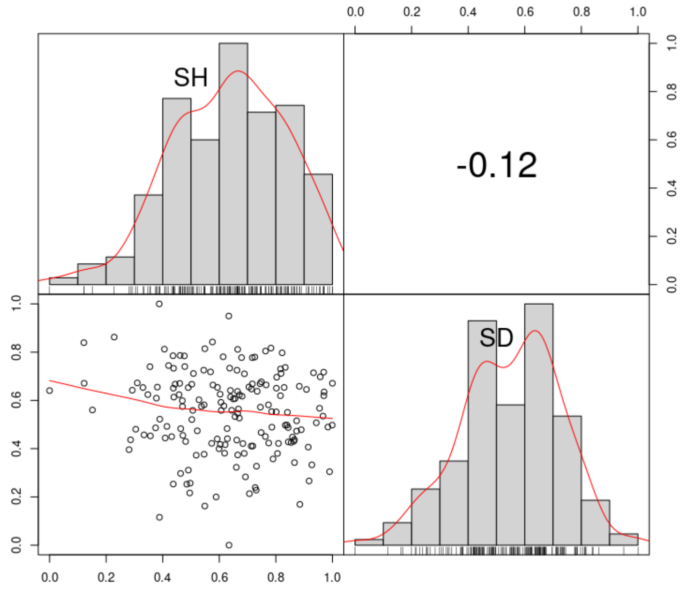


**Supplementary Fig. S4.** Correlations between phenotypic traits of individuals from the cross performed between sugarcane commercial varieties. The evaluated traits were stalk height (SH) and stalk diameter (SD).

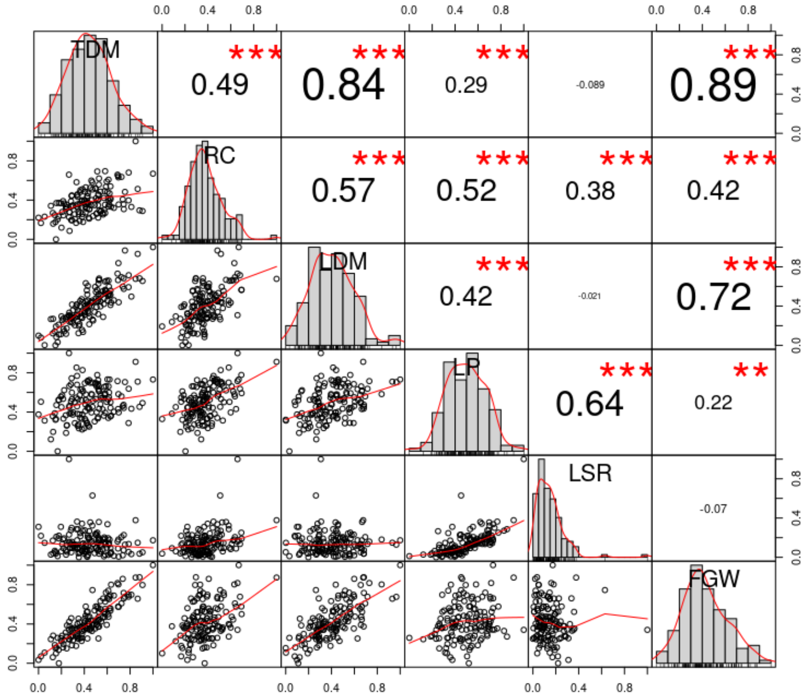


**Supplementary Fig. S5.** Correlations between phenotypic traits of individuals from the cross performed between tetraploid genotypes of *Urochloa decumbens*. The evaluated traits were total dry matter (TDM), regrowth capacity (RC), leaf dry matter (LDM), leaf percentage (LP), leaf stem ratio (LSR) and field green weight (FGW).

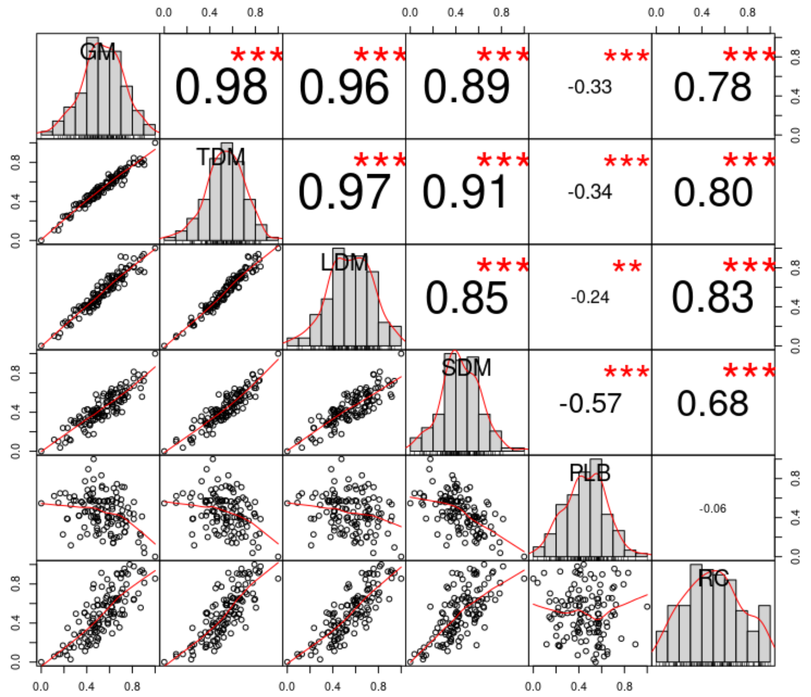


**Supplementary Fig. S6.** Correlations between phenotypic traits of individuals from the cross between genotypes of *Megathyrsus maximus*. The evaluated traits were green matter (GM), total dry matter (TDM), leaf dry matter (LDM), stem dry matter (SDM), percentage of leaf blade (PLB) and regrowth capacity (RC).

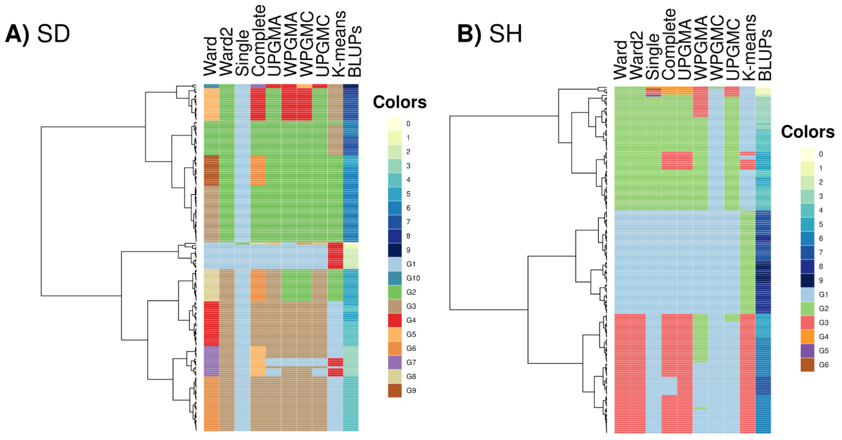


**Supplementary Fig. S7.** Heatmaps and dendrograms for the best linear unbiased predictions (BLUPs) of the evaluated traits (A – stalk diameter (SD) and B – stalk height (SH)) in the sugarcane population using the K-means algorithm and hierarchical clustering methods (Euclidean distance-based): (i) Ward's minimum variance with squared similarities, (ii) Ward's minimum variance without squared similarities, (iii) single, (iv) complete, (v) unweighted pair-group method with arithmetic averaging (UPGMA), (vi) weighted pair-group method with arithmetic averaging (WPGMA), (vii) weighted pair-group method with centroid averaging (WPGMC), and (viii) unweighted pair group method with centroid averaging (UPGMC).

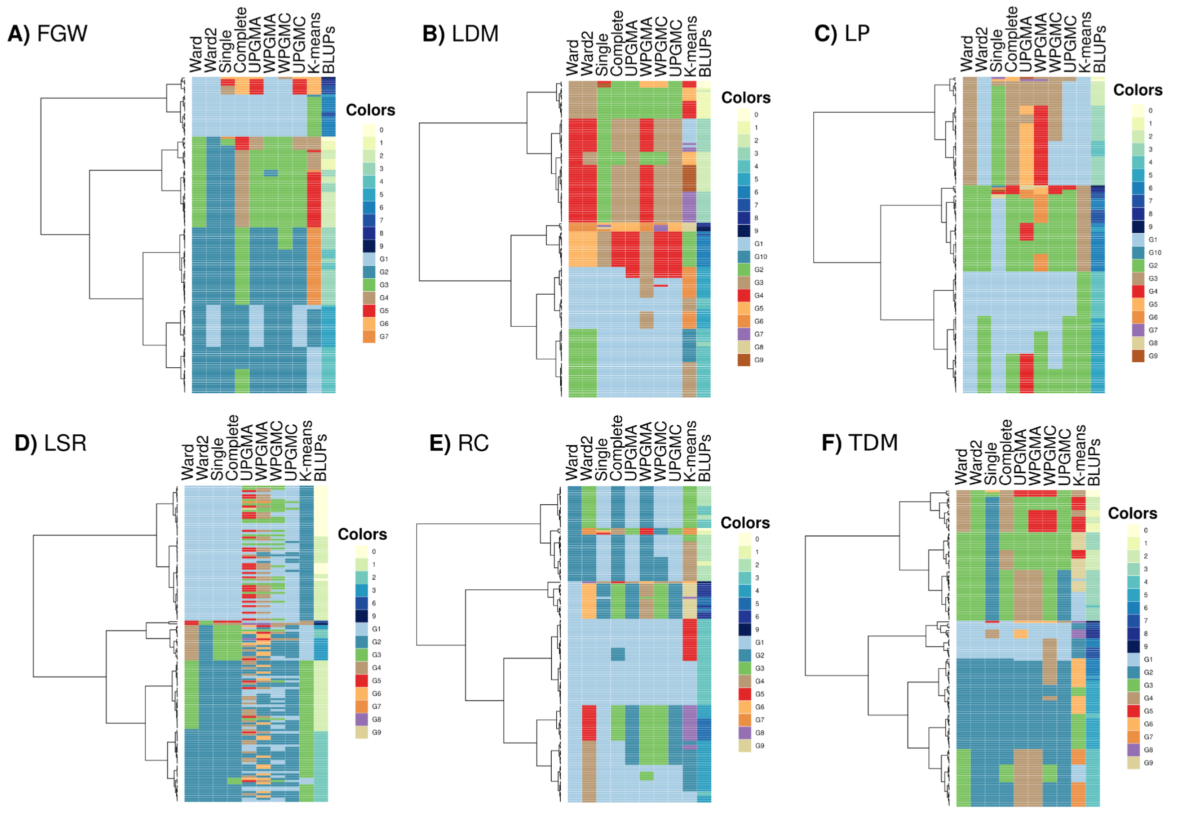


**Supplementary Fig. S8.** Heatmaps and dendrograms for the best linear unbiased predictions (BLUPs) of the evaluated traits (A – field green weight (FGW), B – leaf dry matter (LDM), C – leaf percentage (LP), D – leaf stem ratio (LSR), E – regrowth capacity (RC), and F – total dry matter (TDM)) in the population of *Urochloa decumbens* using the K-means algorithm and hierarchical clustering methods (Euclidean distance-based): (i) Ward's minimum variance with squared similarities, (ii) Ward's minimum variance without squared similarities, (iii) single, (iv) complete, (v) unweighted pair-group method with arithmetic averaging (UPGMA), (vi) weighted pair-group method with arithmetic averaging (WPGMA), (vii) weighted pair-group method with centroid averaging (WPGMC), and (viii) unweighted pair group method with centroid averaging (UPGMC).

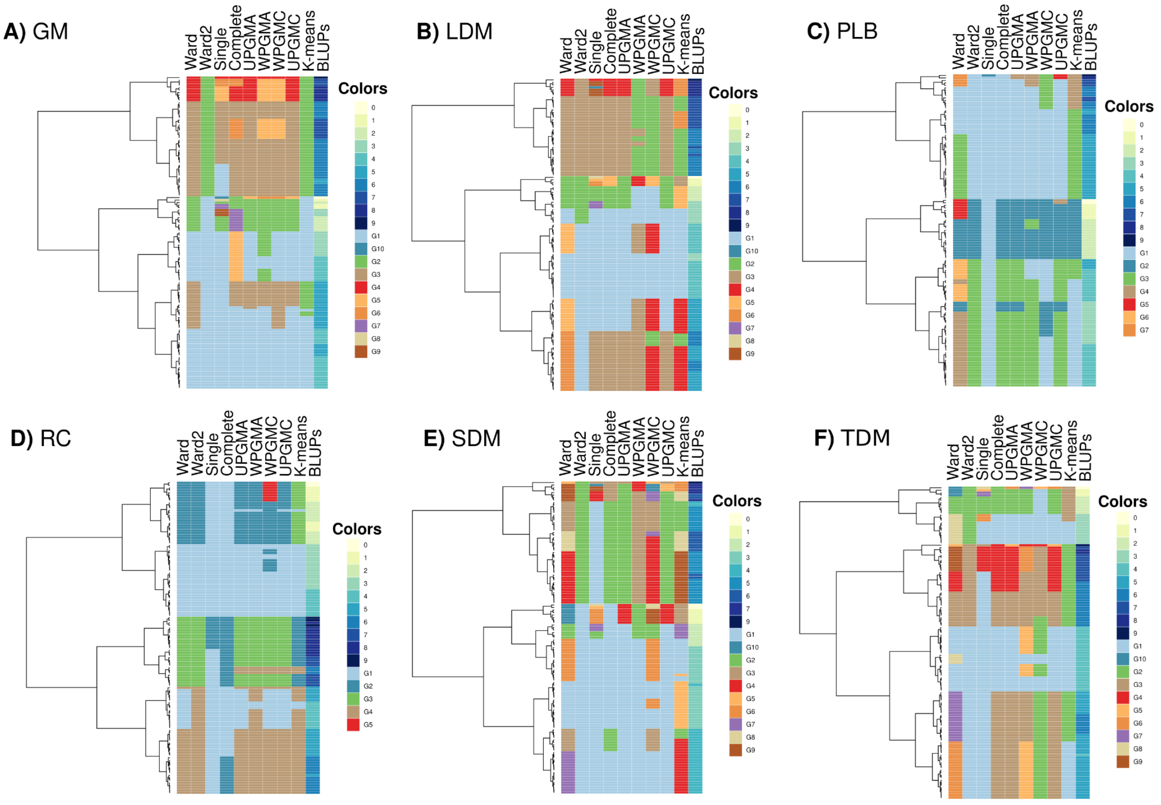


**Supplementary Fig. S9.** Heatmaps and dendrograms for the best linear unbiased predictions (BLUPs) of the evaluated traits (A – green matter (GM), B – leaf dry matter (LDM), C – percentage of leaf blade (PLB), D – regrowth capacity (RC), E – stem dry matter (SDM), and F – total dry matter (TDM)) in the population of *Megathyrsus maximus* using the K-means algorithm and hierarchical clustering methods (Euclidean distance-based): (i) Ward's minimum variance with squared similarities, (ii) Ward's minimum variance without squared similarities, (iii) single, (iv) complete, (v) unweighted pair-group method with arithmetic averaging (UPGMA), (vi) weighted pair-group method with arithmetic averaging (WPGMA), (vii) weighted pair-group method with centroid averaging (WPGMC), and (viii) unweighted pair group method with centroid averaging (UPGMC).

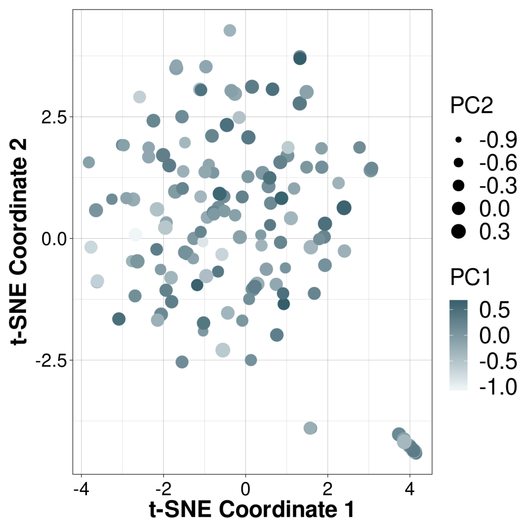


**Supplementary Fig. S10**. T-distributed stochastic neighbor embedding (t-SNE) analysis for genotypic data of *Urochloa decumbens* population, in which the genotypes were colored and sized according to the principal components (PCs) obtained through a principal component analysis (PCA) performed on phenotypic data.

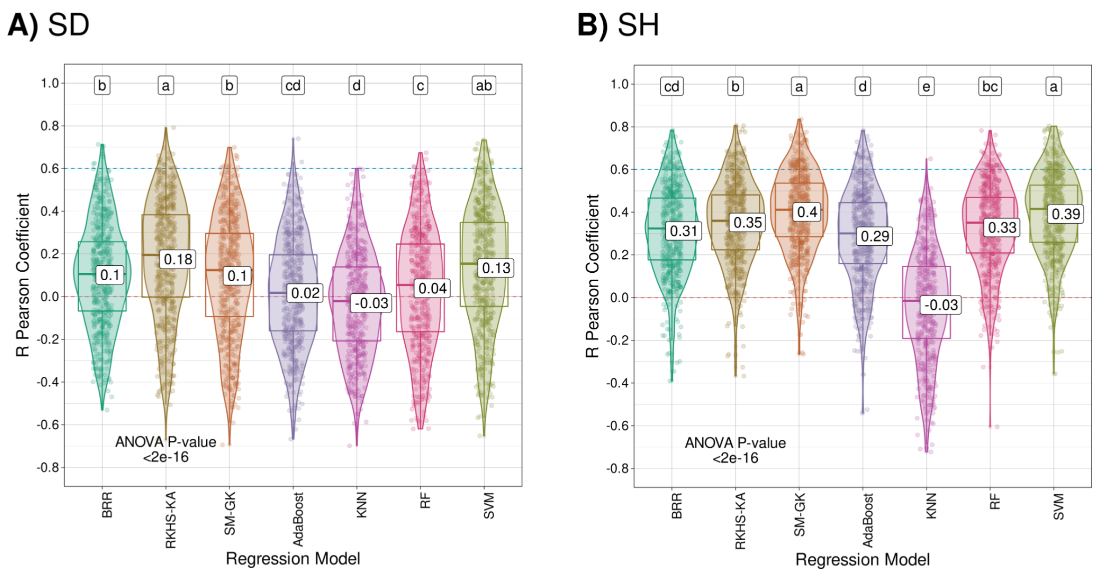


**Supplementary Fig. S11.** Model accuracy distribution using the R Pearson correlation coefficients for the evaluated traits (stalk diameter (SD) and stalk height (SH)) in the sugarcane population considering a 10-fold cross-validation strategy (repeated 50 times). The models evaluated were Bayesian ridge regression (BRR), Bayesian reproducing kernel Hilbert spaces regression with kernel averaging (RKHS-KA), the single-environment main genotypic effect model with a Gaussian kernel (SM-GK), AdaBoost, k-nearest neighbors (KNN), random forest (RF), and support vector machine (SVM). Mean values are presented in the boxplots, and the labels at the top of the plots represent the results from Tukey's multiple comparisons test (p value of 0.05).

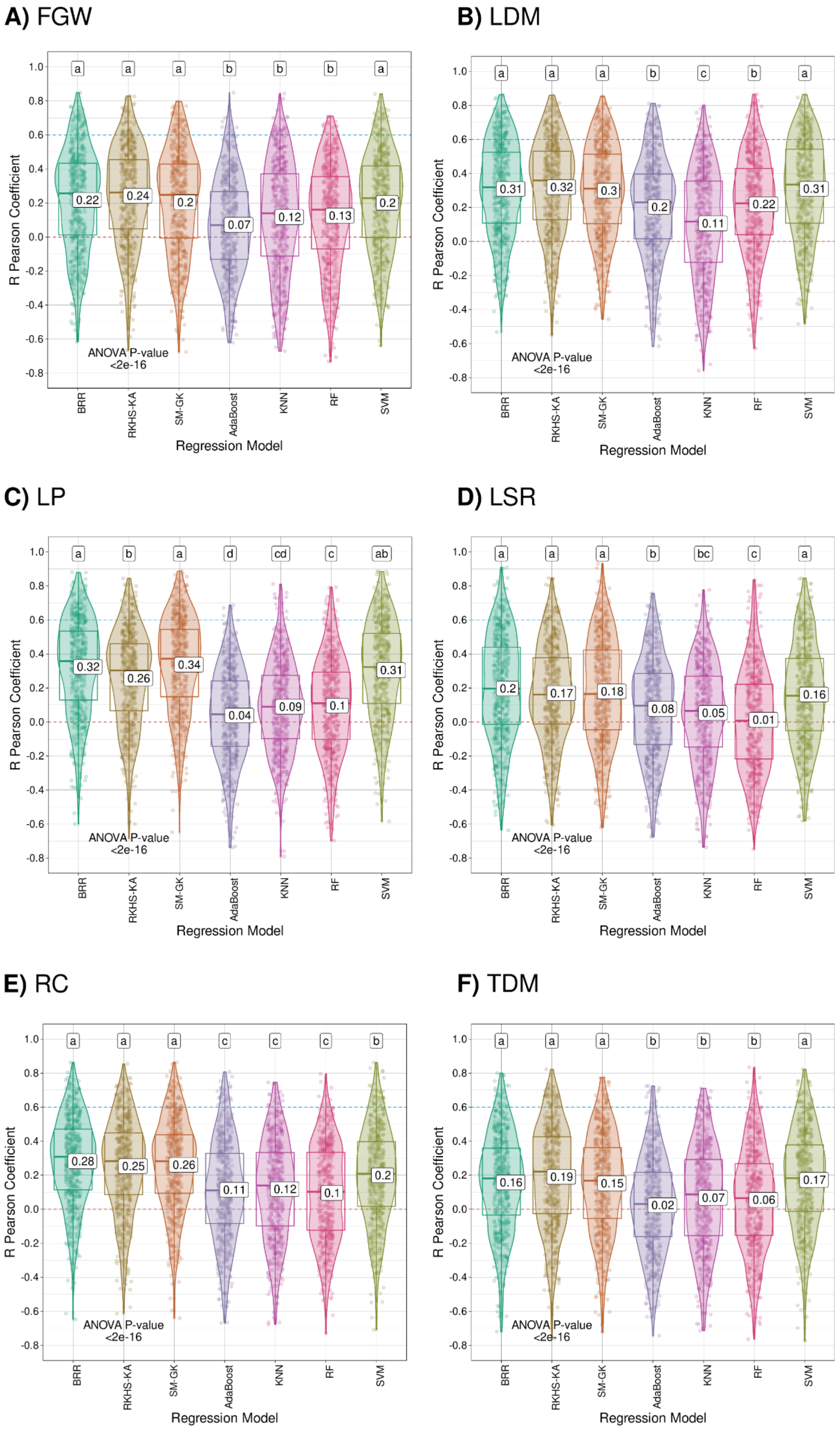
A) FGW
B) LDM
C) LP
D) LSR
E) RC
F) TDM
R Pearson Coefficient
Regression Model
BRR
RKHS-KA
SM-GK
AdaBoost
KNN
RF
SVM
1.0
0.8
0.6
0.4
0.2
0.0
-0.2
-0.4
-0.6
-0.8
ANOVA P-value
<2e-16
a a a b b b a
0.22 0.24 0.2 0.07 0.12 0.13 0.2
a a a b c b a
0.31 0.32 0.3 0.2 0.11 0.22 0.31
a b a d cd c ab
0.32 0.26 0.34 0.04 0.09 0.1 0.31
a a a b bc c a
0.2 0.17 0.18 0.08 0.05 0.01 0.16
a a a c c c b
0.28 0.25 0.26 0.11 0.12 0.1 0.2
a a a b b b a
0.16 0.19 0.15 0.02 0.07 0.06 0.17

**Supplementary Fig. S12.** Model accuracy distribution using the R Pearson correlation coefficients for the evaluated traits (A – field green weight (FGW), B – leaf dry matter (LDM), C – leaf percentage (LP), D – leaf stem ratio (LSR), E – regrowth capacity (RC), and F – total dry matter (TDM)) in the *Urochloa decumbens* population considering a 10-fold cross-validation strategy (repeated 50 times). The models evaluated were Bayesian ridge regression (BRR), Bayesian reproducing kernel Hilbert spaces regression with kernel averaging (RKHS-KA), the single-environment main genotypic effect model with a Gaussian kernel (SM-GK), AdaBoost, k-nearest neighbors (KNN), random forest (RF), and support vector machine (SVM). Mean values are presented in the boxplots, and the labels at the top of the plots represent the results from Tukey's multiple comparisons test (p value of 0.05).

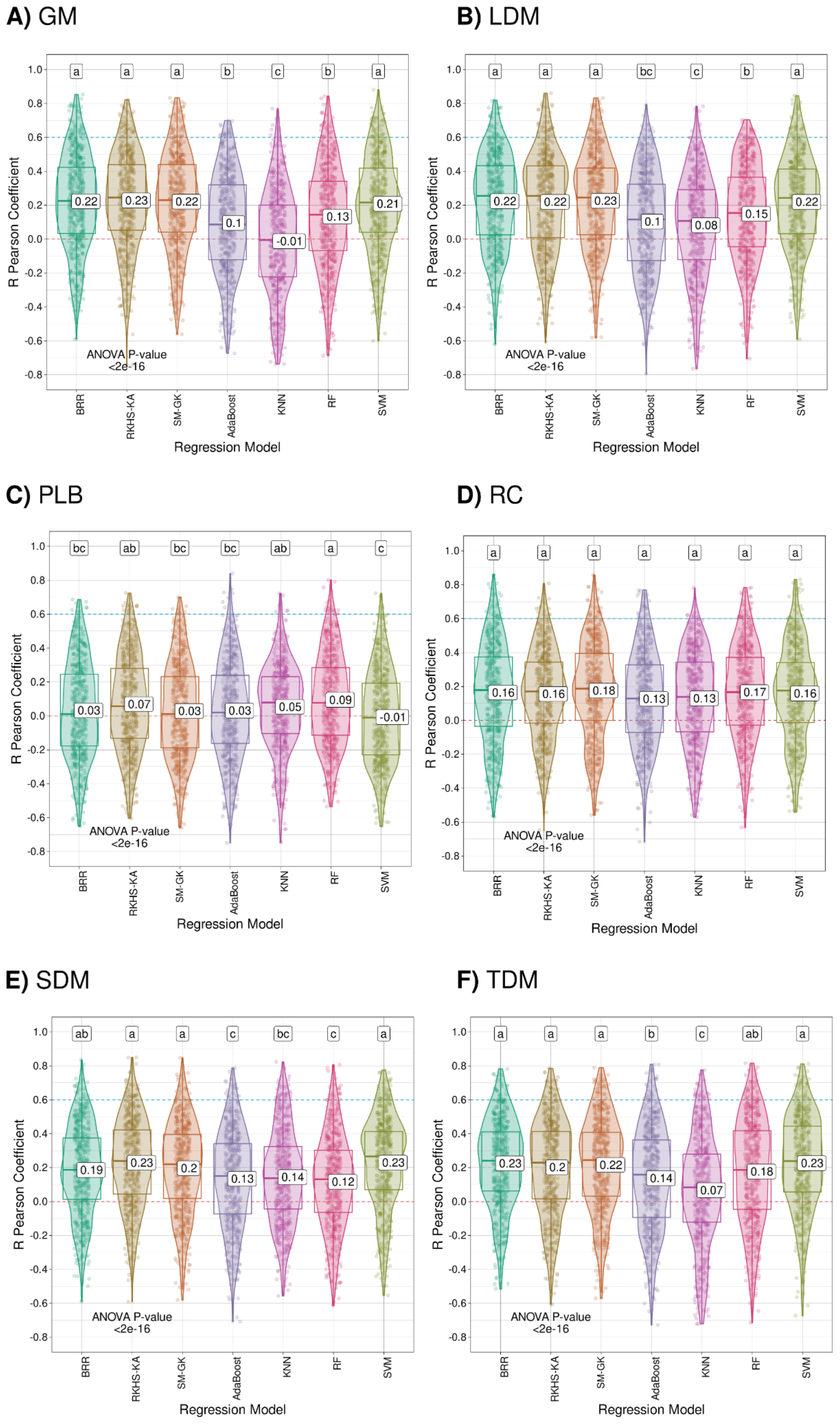
A) GM
B) LDM
C) PLB
D) RC
E) SDM
F) TDM
R Pearson Coefficient
Regression Model
BRR
RKHS-KA
SM-GK
AdaBoost
KNN
RF
SVM
ANOVA P-value
<2e-16
a a a b c b a
0.22 0.23 0.22 0.1 -0.01 0.13 0.21
a a a bc c b a
0.22 0.22 0.23 0.1 0.08 0.15 0.22
bc ab bc bc ab a c
0.03 0.07 0.03 0.03 0.05 0.09 -0.01
a a a a a a a
0.16 0.16 0.18 0.13 0.13 0.17 0.16
ab a a c bc c a
0.19 0.23 0.2 0.13 0.14 0.12 0.23
a a a b c ab a
0.23 0.2 0.22 0.14 0.07 0.18 0.23

**Supplementary Fig. S13.** Model accuracy distribution using the R Pearson correlation coefficients for the evaluated traits (A – green matter (GM), B – leaf dry matter (LDM), C – percentage of leaf blade (PLB), D – regrowth capacity (RC), E – stem dry matter (SDM), and F – total dry matter (TDM)) in the *Megathyrsus maximus* population considering a 10-fold cross-validation strategy (repeated 50 times). The models evaluated were Bayesian ridge regression (BRR), Bayesian reproducing kernel Hilbert spaces regression with kernel averaging (RKHS-KA), the single-environment main genotypic effect model with a Gaussian kernel (SM-GK), AdaBoost, k-nearest neighbors (KNN), random forest (RF), and support vector machine (SVM). Mean values are presented in the boxplots, and the labels at the top of the plots represent the results from Tukey's multiple comparisons test (p value of 0.05).

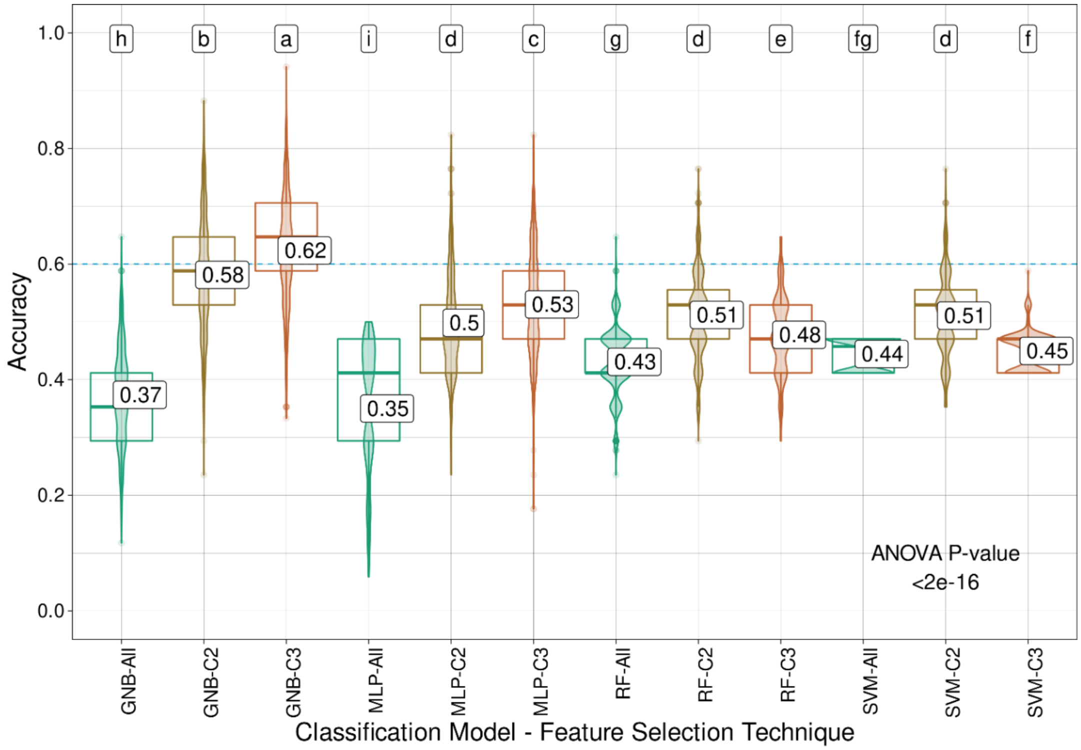
h b a i d c g d e fg d f
0.58
0.62
0.37
0.35
0.5
0.53
0.43
0.51
0.48
0.44
0.51
0.45
ANOVA P-value
<2e-16
Accuracy
GNB-All GNB-C2 GNB-C3 MLP-All MLP-C2 MLP-C3 RF-All RF-C2 RF-C3 SVM-All SVM-C2 SVM-C3
Classification Model - Feature Selection Technique

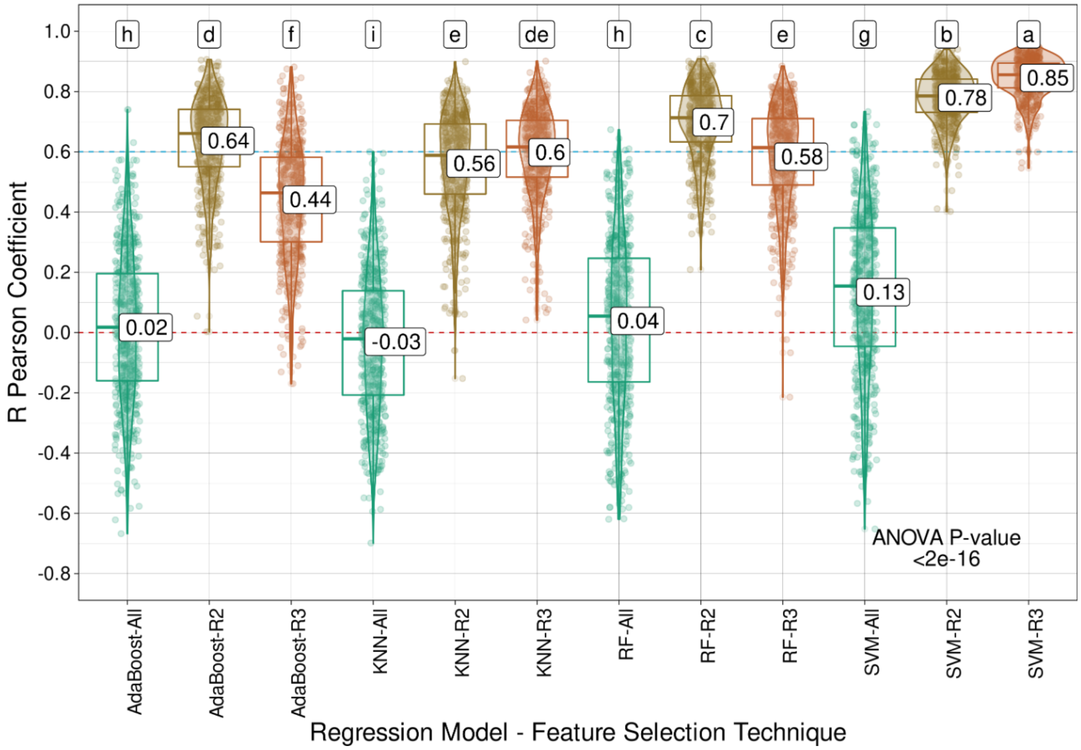
h d f i e de h c e g b a
0.02
0.64
0.44
-0.03
0.56
0.6
0.04
0.7
0.58
0.13
0.78
0.85
ANOVA P-value
<2e-16
R Pearson Coefficient
AdaBoost-All AdaBoost-R2 AdaBoost-R3 KNN-All KNN-R2 KNN-R3 RF-All RF-R2 RF-R3 SVM-All SVM-R2 SVM-R3
Regression Model - Feature Selection Technique

**Supplementary Fig. S14**. Model performance for sugarcane stem diameter (SD) using classification (C) algorithms (Gaussian naive Bayes (GNB), multilayer perceptron (MLP), random forest (RF), and support vector machine (SVM)) and regression (R) models (AdaBoost, k-nearest neighbors (KNN), RF and SVM), coupled with the intersection of the results from the three feature selection (FS) strategies tested (C3/I3), the intersection of at least two out of the three FS methods (C2/I2), or the entire set of markers (All). Mean values are presented in the boxplots, and the labels at the top of the plots represent the results from Tukey's multiple comparisons test (p value of 0.05).

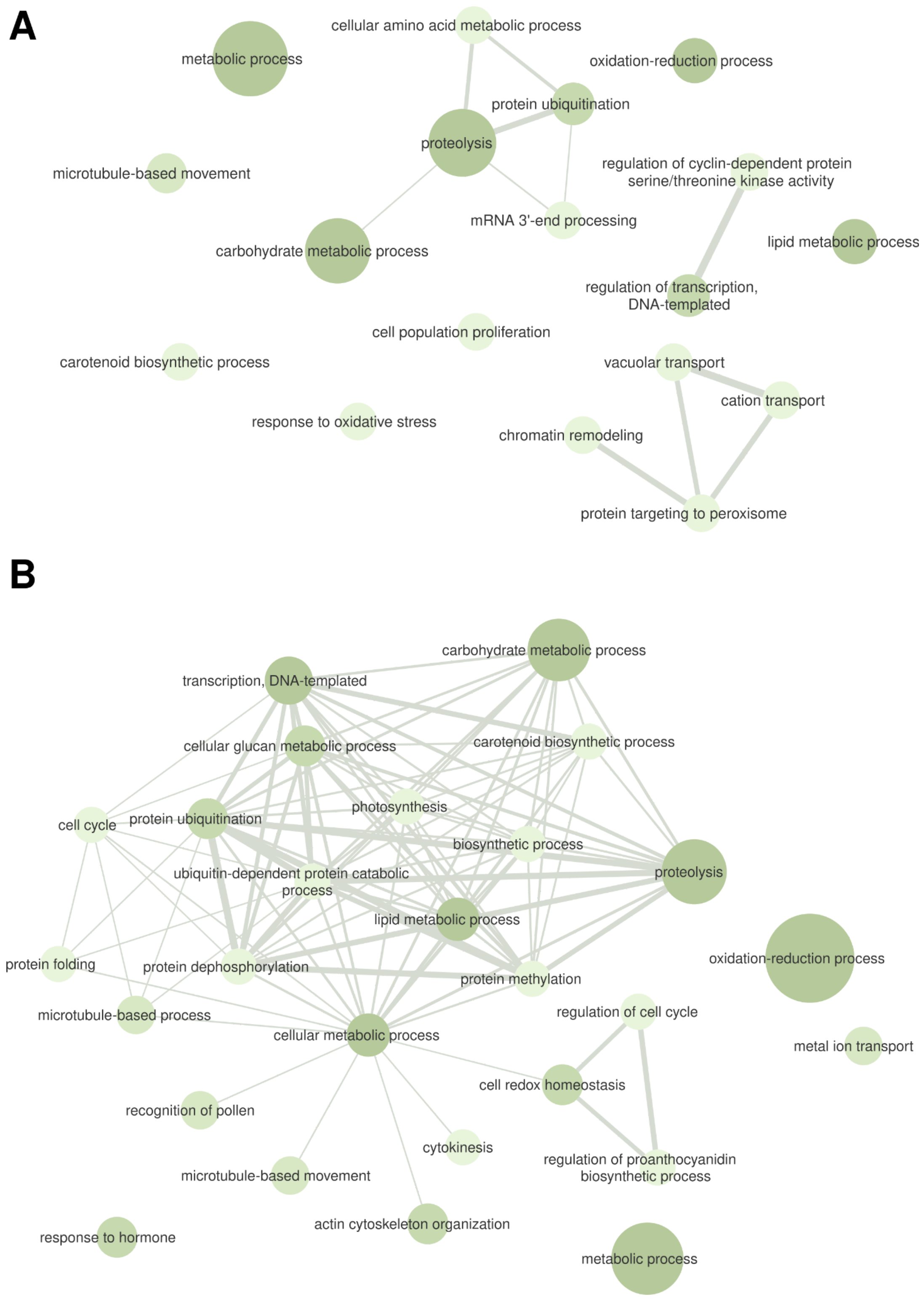
A
metabolic process
cellular amino acid metabolic process
oxidation-reduction process
protein ubiquitination
proteolysis
microtubule-based movement
regulation of cyclin-dependent protein serine/threonine kinase activity
mRNA 3'-end processing
carbohydrate metabolic process
lipid metabolic process
regulation of transcription, DNA-templated
cell population proliferation
carotenoid biosynthetic process
vacuolar transport
cation transport
response to oxidative stress
chromatin remodeling
protein targeting to peroxisome
B
carbohydrate metabolic process
transcription, DNA-templated
cellular glucan metabolic process
carotenoid biosynthetic process
photosynthesis
cell cycle
protein ubiquitination
biosynthetic process
ubiquitin-dependent protein catabolic process
proteolysis
lipid metabolic process
oxidation-reduction process
protein folding
protein dephosphorylation
protein methylation
microtubule-based process
regulation of cell cycle
cellular metabolic process
metal ion transport
cell redox homeostasis
recognition of pollen
cytokinesis
regulation of proanthocyanidin biosynthetic process
microtubule-based movement
actin cytoskeleton organization
response to hormone
metabolic process

**Supplementary Fig. S15**. Gene ontology (GO) terms from the biological process category associated with the genomic regions containing the markers selected by the intersection of the results from at least two out of the three feature selection (FS) strategies for sugarcane stem diameter (SD). These markers were selected through (**A**) classification algorithms and (**B**) regression models.

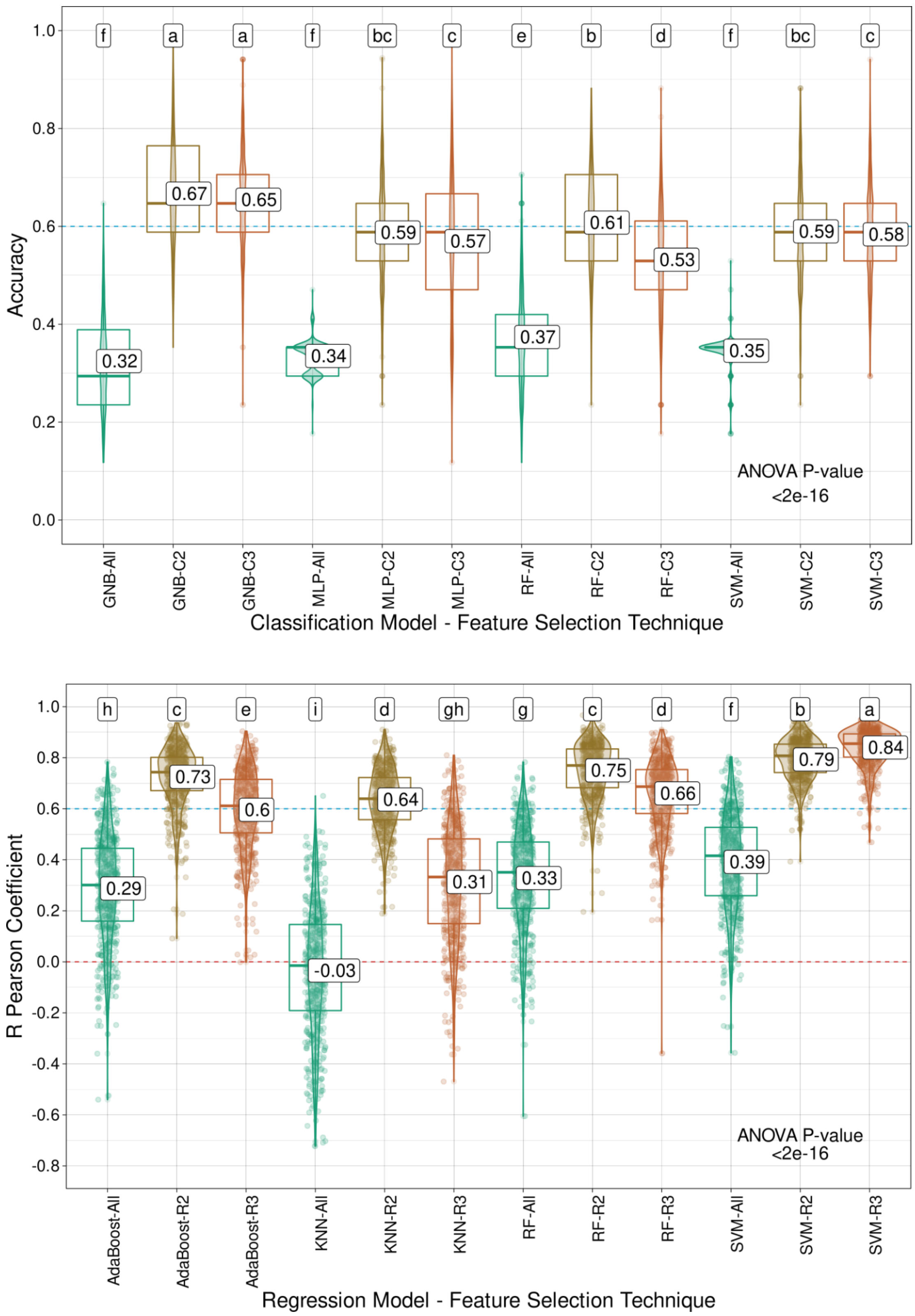

Accuracy
f a a f bc c e b d f bc c
0.32 0.67 0.65 0.34 0.59 0.57 0.37 0.61 0.53 0.35 0.59 0.58
ANOVA P-value
<2e-16
GNB-All GNB-C2 GNB-C3 MLP-All MLP-C2 MLP-C3 RF-All RF-C2 RF-C3 SVM-All SVM-C2 SVM-C3
Classification Model - Feature Selection Technique
R Pearson Coefficient
h c e i d gh g c d f b a
0.29 0.73 0.6 -0.03 0.64 0.31 0.33 0.75 0.66 0.39 0.79 0.84
ANOVA P-value
<2e-16
AdaBoost-All AdaBoost-R2 AdaBoost-R3 KNN-All KNN-R2 KNN-R3 RF-All RF-R2 RF-R3 SVM-All SVM-R2 SVM-R3
Regression Model - Feature Selection Technique

**Supplementary Fig. S16**. Model performance for sugarcane stem height (SH) using classification (C) algorithms (Gaussian naive Bayes (GNB), multilayer perceptron (MLP), random forest (RF), and support vector machine (SVM)) and regression (R) models (AdaBoost, k-nearest neighbors (KNN), RF and SVM), coupled with the intersection of the results from the three feature selection (FS) strategies tested (C3/I3), the intersection of at least two out of the three FS methods (C2/I2), or the entire set of markers (All). Mean values are presented in the boxplots, and the labels at the top of the plots represent the results from Tukey's multiple comparisons test (p value of 0.05).

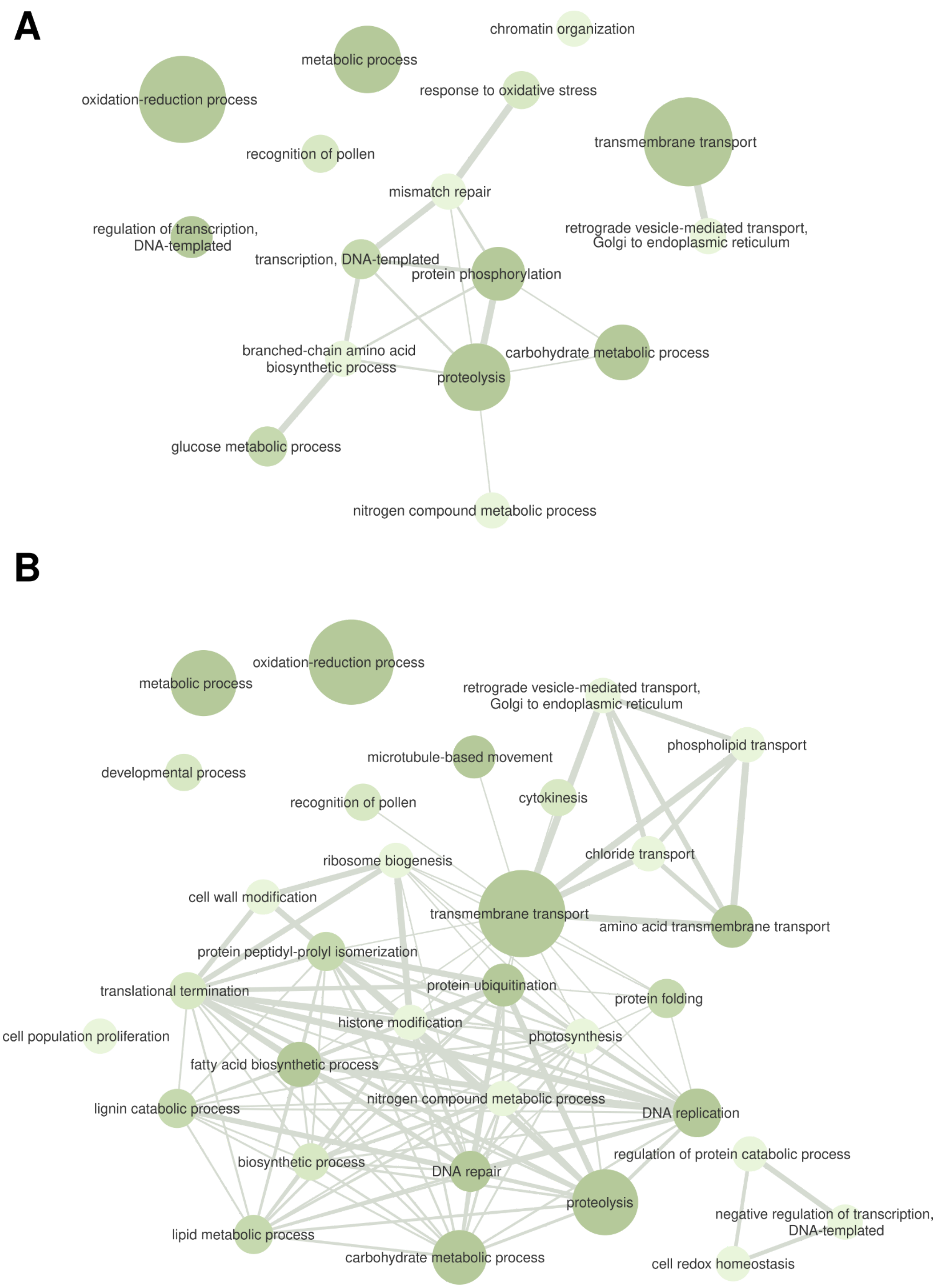


**Supplementary Fig. S17**. Gene ontology (GO) terms from the biological process category associated with the genomic regions containing the markers selected by the intersection of the

results from at least two out of the three feature selection (FS) strategies for sugarcane stem height (SH). These markers were selected through (**A**) classification algorithms and (**B**) regression models.

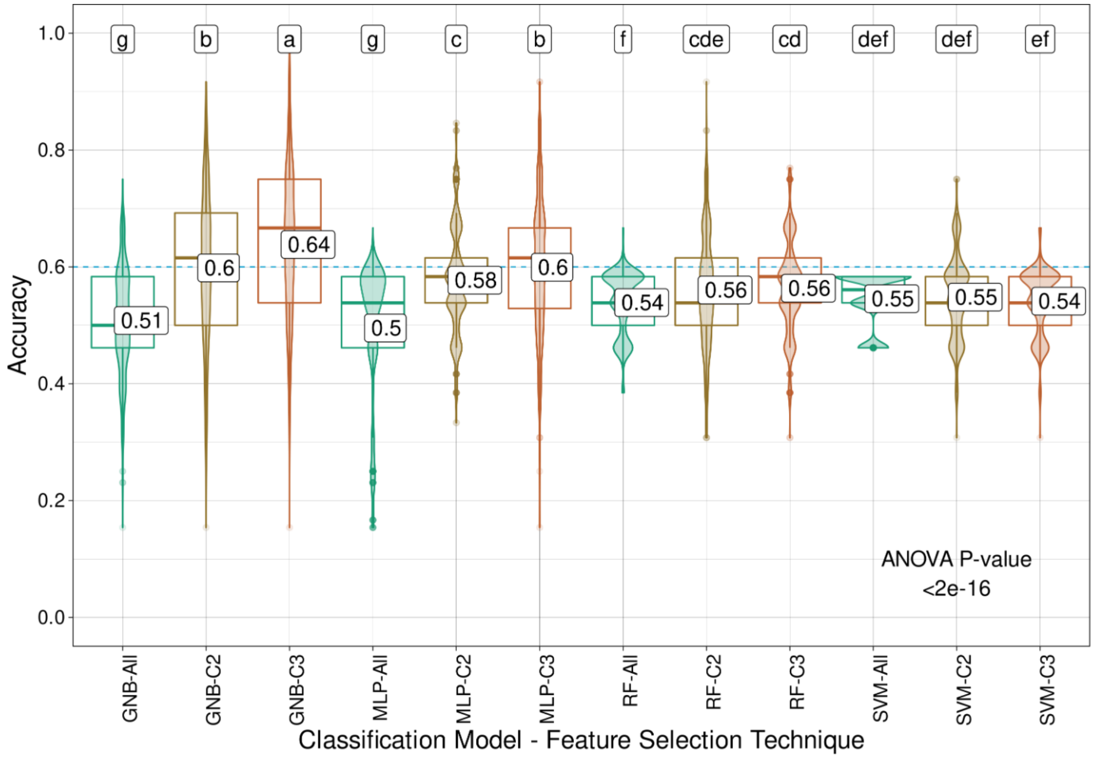
g b a g c b f cde cd def def ef
0.51 0.6 0.64 0.5 0.58 0.6 0.54 0.56 0.56 0.55 0.55 0.54
ANOVA P-value
<2e-16
Accuracy
GNB-All GNB-C2 GNB-C3 MLP-All MLP-C2 MLP-C3 RF-All RF-C2 RF-C3 SVM-All SVM-C2 SVM-C3
Classification Model - Feature Selection Technique

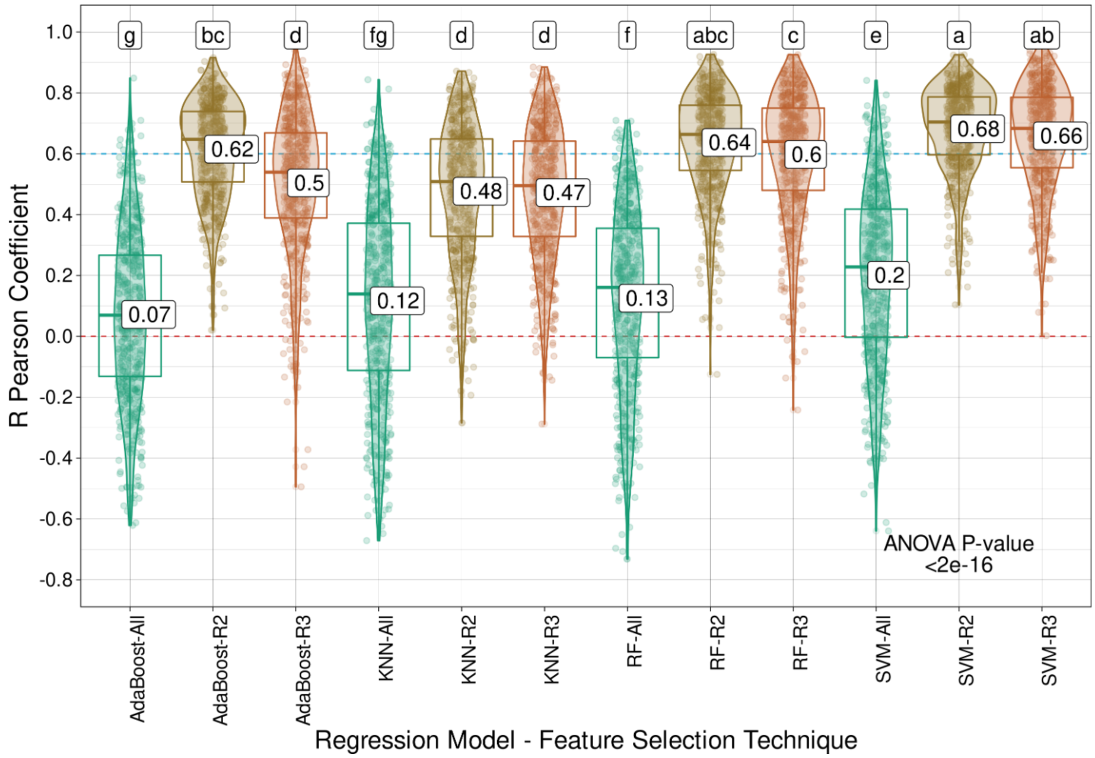
g bc d fg d d f abc c e a ab
0.07 0.62 0.5 0.12 0.48 0.47 0.13 0.64 0.6 0.2 0.68 0.66
ANOVA P-value
<2e-16
R Pearson Coefficient
AdaBoost-All AdaBoost-R2 AdaBoost-R3 KNN-All KNN-R2 KNN-R3 RF-All RF-R2 RF-R3 SVM-All SVM-R2 SVM-R3
Regression Model - Feature Selection Technique

**Supplementary Fig. S18**. Model performance for *Urochloa decumbens* field green weight (FGW) using classification (C) algorithms (Gaussian naive Bayes (GNB), multilayer perceptron (MLP), random forest (RF), and support vector machine (SVM)) and regression (R) models (AdaBoost, k-nearest neighbors (KNN), RF and SVM), coupled with the intersection of the results from the three feature selection (FS) strategies tested (C3/I3), the intersection of at least two out of the three FS methods (C2/I2), or the entire set of markers (All). Mean values are presented in the boxplots, and the labels at the top of the plots represent the results from Tukey's multiple comparisons test (p value of 0.05).

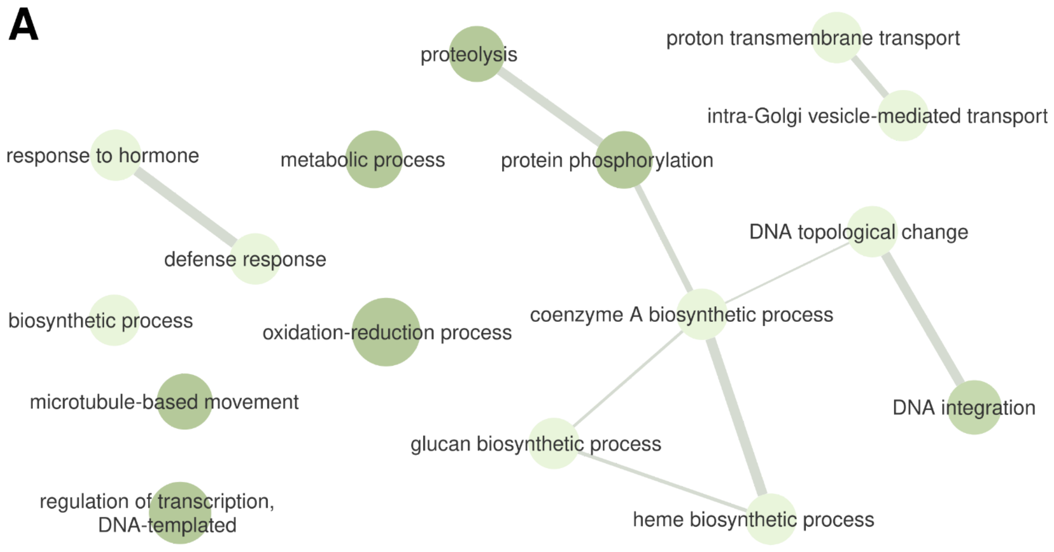


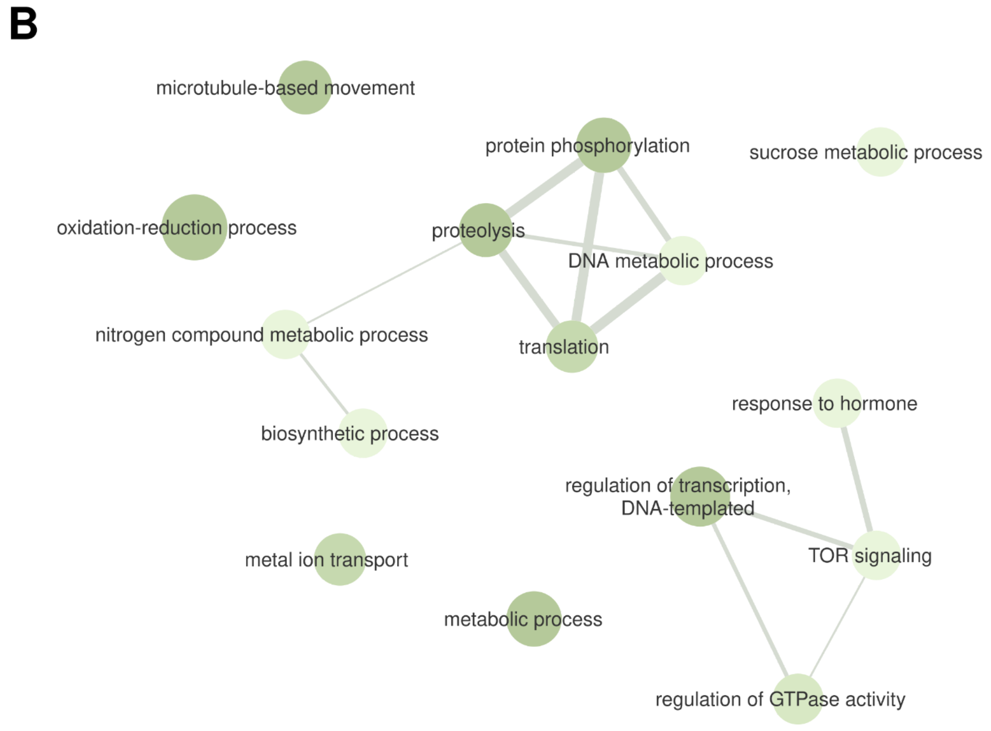


**Supplementary Fig. S19**. Gene ontology (GO) terms from the biological process category associated with the genomic regions containing the markers selected by the intersection of the results from at least two out of the three feature selection (FS) strategies for *Urochloa decumbens*

field green weight (FGW). These markers were selected through (**A**) classification algorithms and (**B**) regression models.

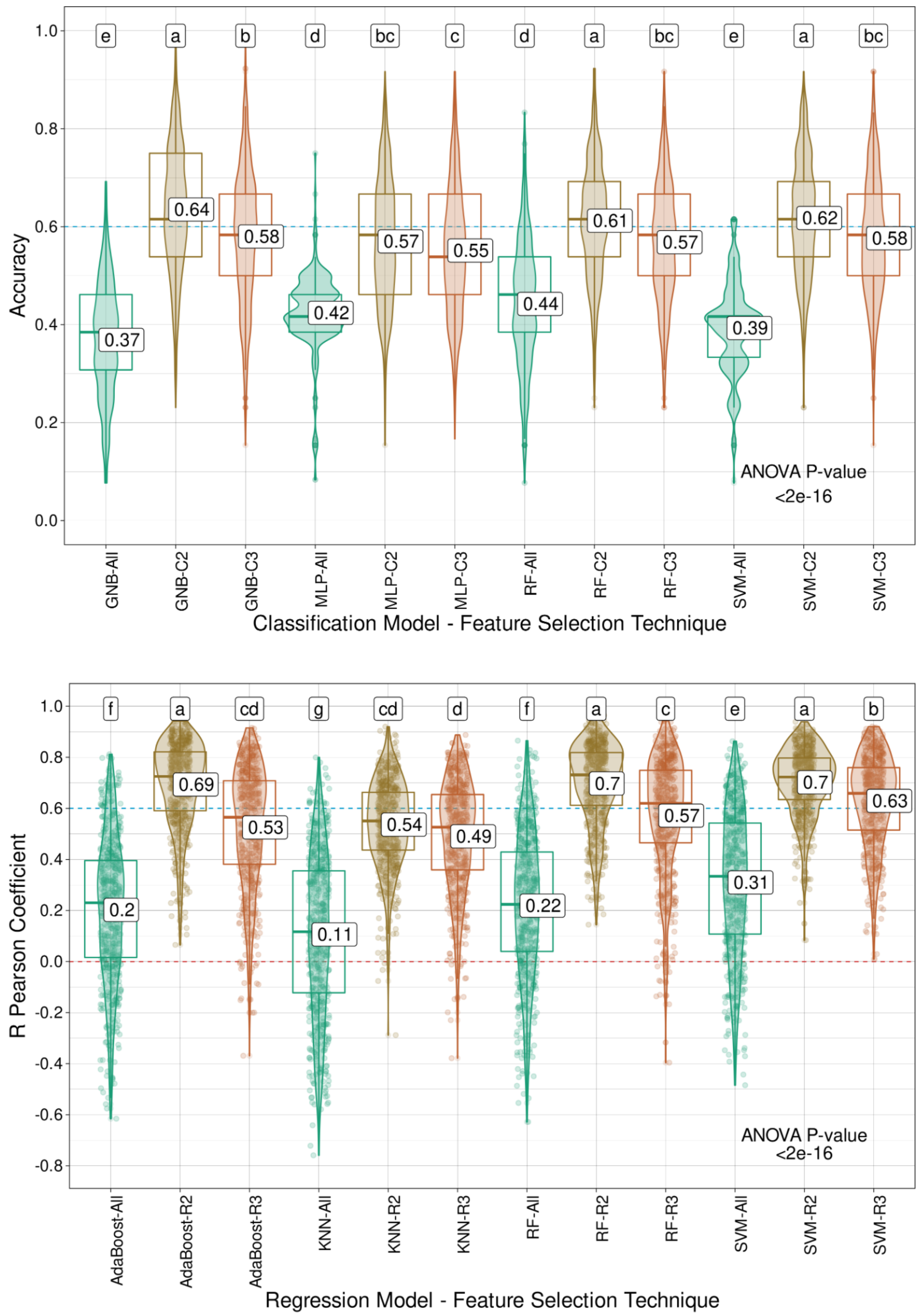
e a b d bc c d a bc e a bc
0.37 0.64 0.58 0.42 0.57 0.55 0.44 0.61 0.57 0.39 0.62 0.58
ANOVA P-value
<2e-16
Accuracy
GNB-All GNB-C2 GNB-C3 MLP-All MLP-C2 MLP-C3 RF-All RF-C2 RF-C3 SVM-All SVM-C2 SVM-C3
Classification Model - Feature Selection Technique
f a cd g cd d f a c e a b
0.2 0.69 0.53 0.11 0.54 0.49 0.22 0.7 0.57 0.31 0.7 0.63
ANOVA P-value
<2e-16
R Pearson Coefficient
AdaBoost-All AdaBoost-R2 AdaBoost-R3 KNN-All KNN-R2 KNN-R3 RF-All RF-R2 RF-R3 SVM-All SVM-R2 SVM-R3
Regression Model - Feature Selection Technique

**Supplementary Fig. S20**. Model performance for *Urochloa decumbens* leaf dry matter (LDM) using classification (C) algorithms (Gaussian naive Bayes (GNB), multilayer perceptron (MLP), random forest (RF), and support vector machine (SVM)) and regression (R) models (AdaBoost, k-nearest neighbors (KNN), RF and SVM), coupled with the intersection of the results from the three feature selection (FS) strategies tested (C3/I3), the intersection of at least two out of the three FS methods (C2/I2), or the entire set of markers (All). Mean values are presented in the boxplots, and the labels at the top of the plots represent the results from Tukey's multiple comparisons test (p value of 0.05).

**A**

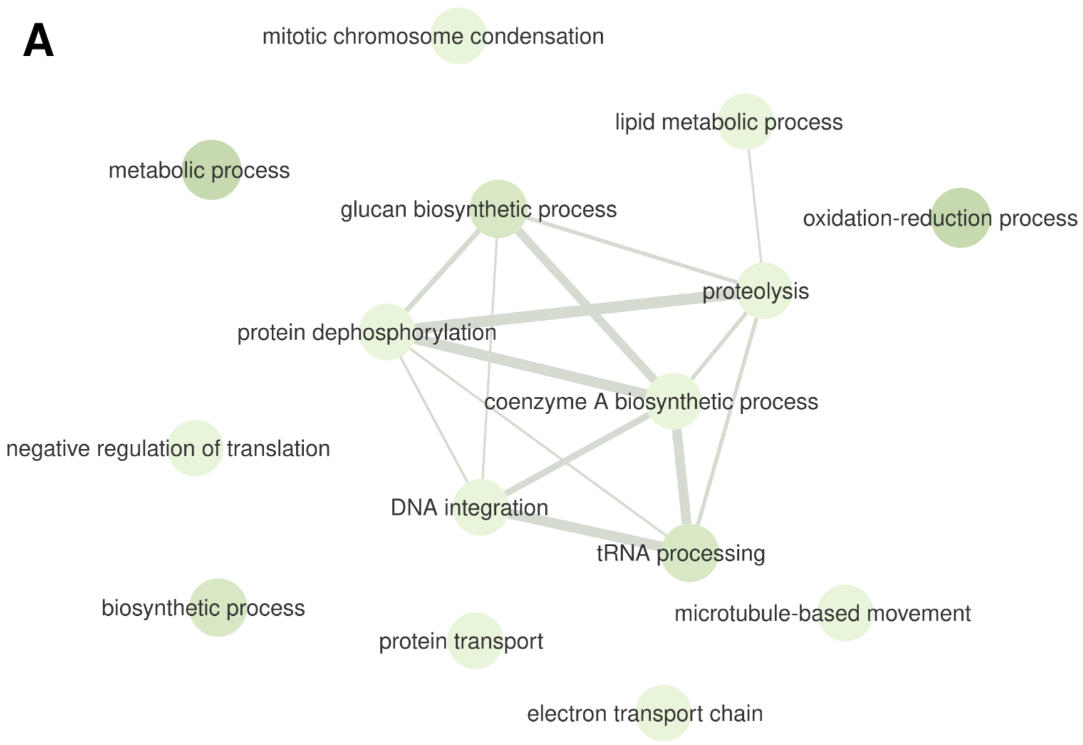


**B**

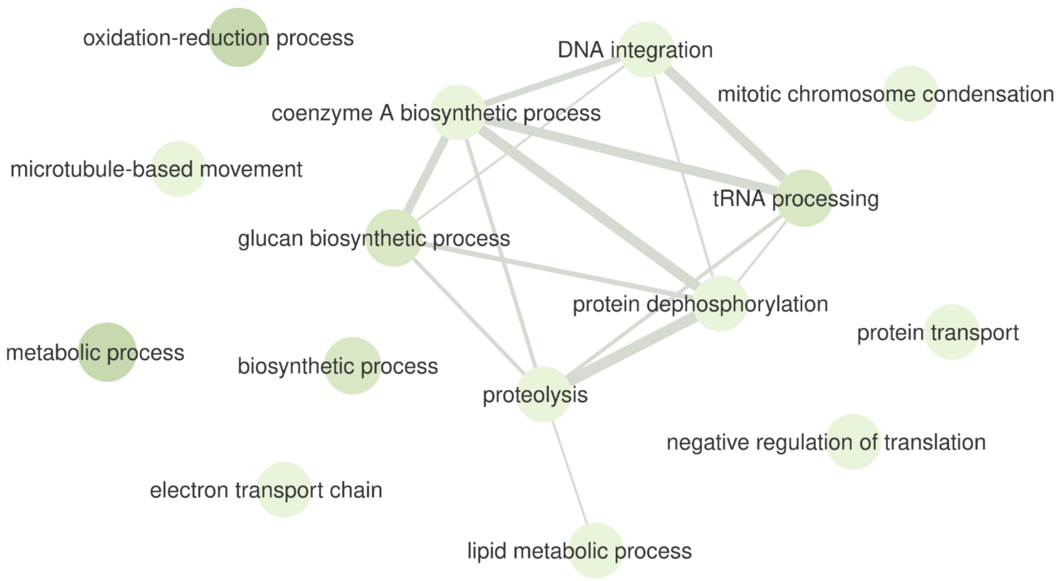


**Supplementary Fig. S21**. Gene ontology (GO) terms from the biological process category associated with the genomic regions containing the markers selected by the intersection of the

results from at least two out of the three feature selection (FS) strategies for *Urochloa decumbens* leaf dry matter (LDM). These markers were selected through (**A**) classification algorithms and (**B**) regression models.

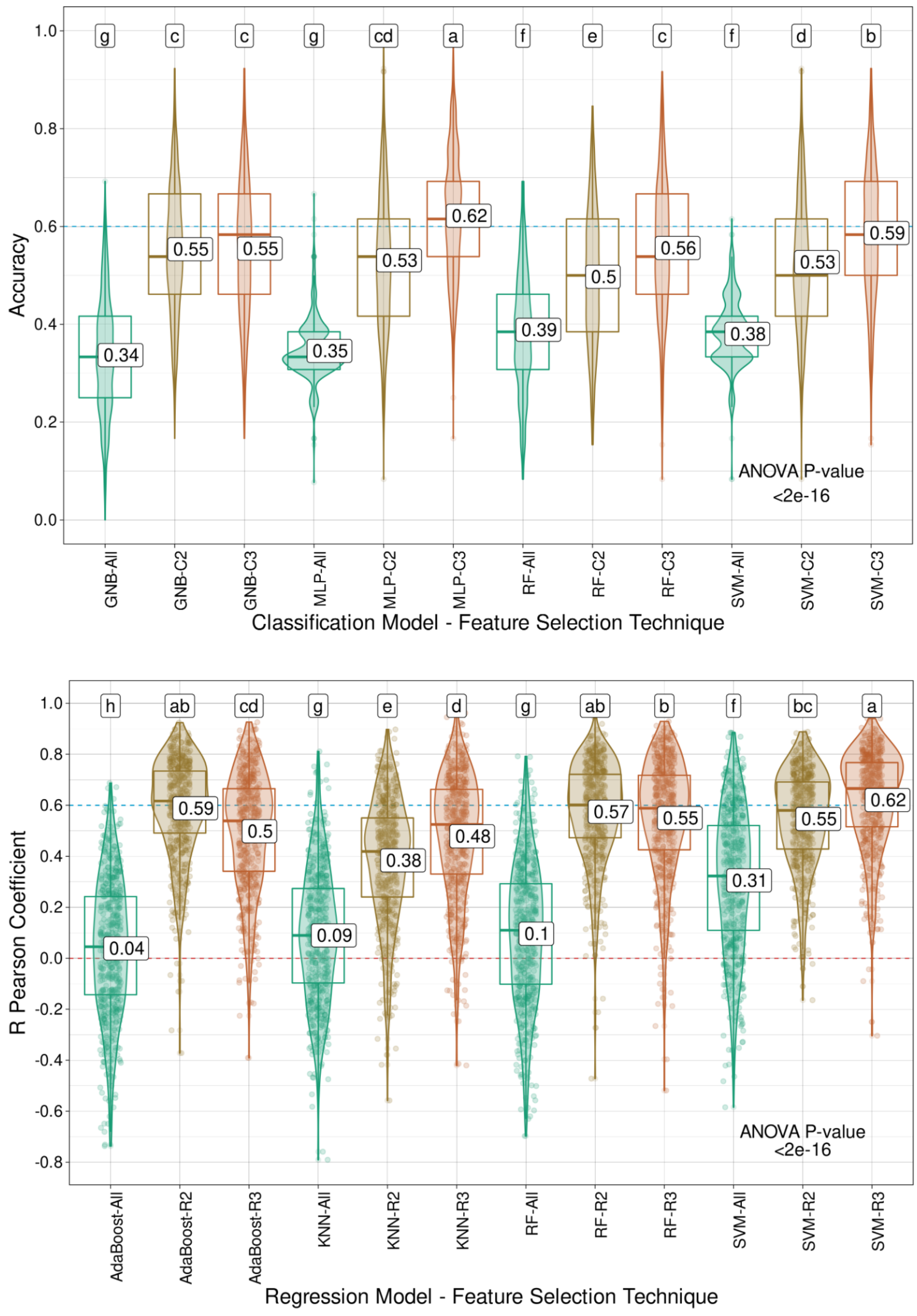
g
c
c
g
cd
a
f
e
c
f
d
b
Accuracy
0.34
0.55
0.55
0.35
0.53
0.62
0.39
0.5
0.56
0.38
0.53
0.59
ANOVA P-value
<2e-16
GNB-All
GNB-C2
GNB-C3
MLP-All
MLP-C2
MLP-C3
RF-All
RF-C2
RF-C3
SVM-All
SVM-C2
SVM-C3
Classification Model - Feature Selection Technique
h
ab
cd
g
e
d
g
ab
b
f
bc
a
R Pearson Coefficient
0.04
0.59
0.5
0.09
0.38
0.48
0.1
0.57
0.55
0.31
0.55
0.62
ANOVA P-value
<2e-16
AdaBoost-All
AdaBoost-R2
AdaBoost-R3
KNN-All
KNN-R2
KNN-R3
RF-All
RF-R2
RF-R3
SVM-All
SVM-R2
SVM-R3
Regression Model - Feature Selection Technique

**Supplementary Fig. S22**. Model performance for *Urochloa decumbens* leaf percentage (LP) using classification (C) algorithms (Gaussian naive Bayes (GNB), multilayer perceptron (MLP), random forest (RF), and support vector machine (SVM)) and regression (R) models (AdaBoost, k-nearest neighbors (KNN), RF and SVM), coupled with the intersection of the results from the three feature selection (FS) strategies tested (C3/I3), the intersection of at least two out of the three FS methods (C2/I2), or the entire set of markers (All). Mean values are presented in the boxplots, and the labels at the top of the plots represent the results from Tukey's multiple comparisons test (p value of 0.05).

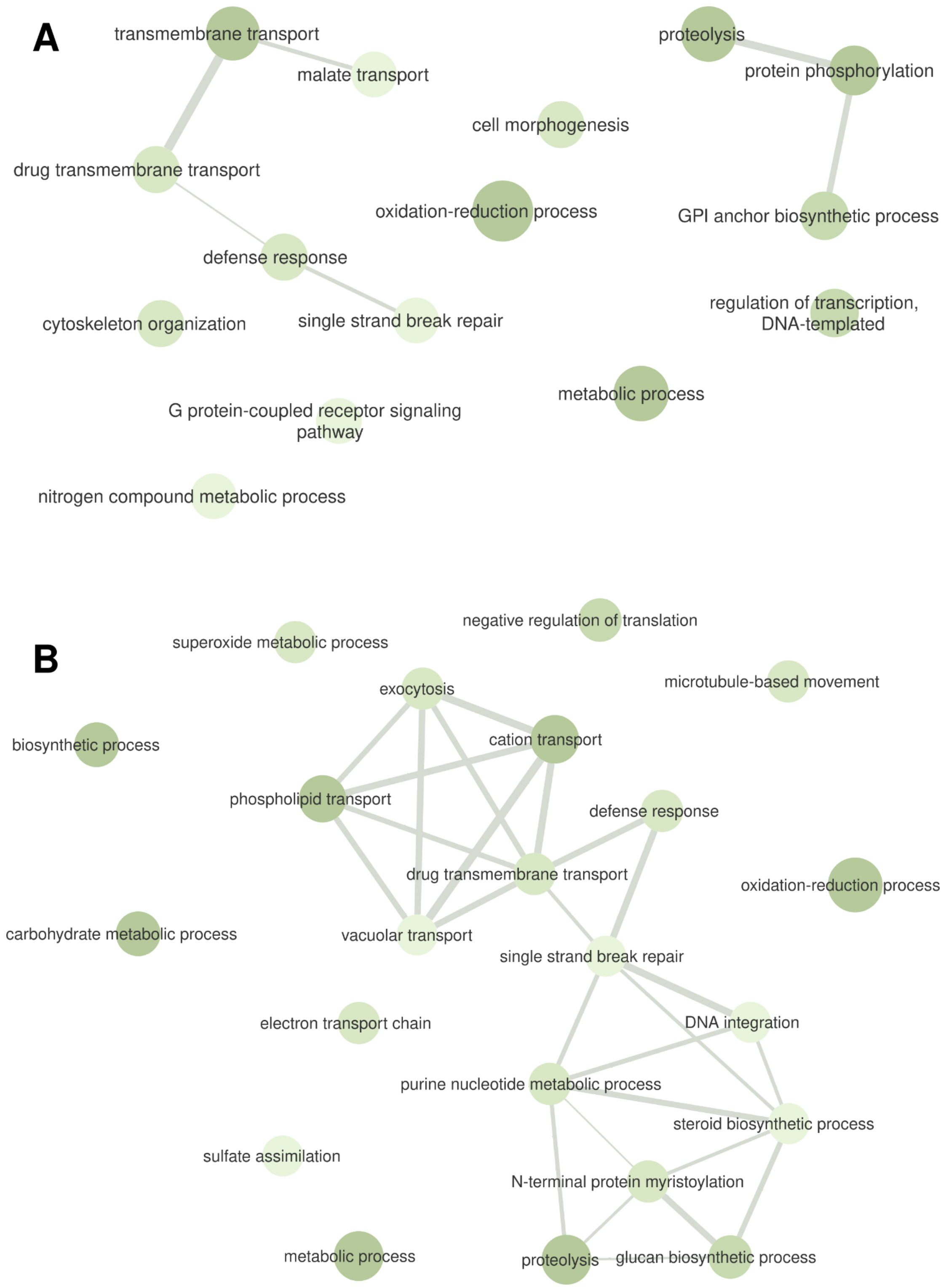


**Supplementary Fig. S23**. Gene ontology (GO) terms from the biological process category associated with the genomic regions containing the markers selected by the intersection of the

results from at least two out of the three feature selection (FS) strategies for *Urochloa decumbens* leaf percentage (LP). These markers were selected through (**A**) classification algorithms and (**B**) regression models.

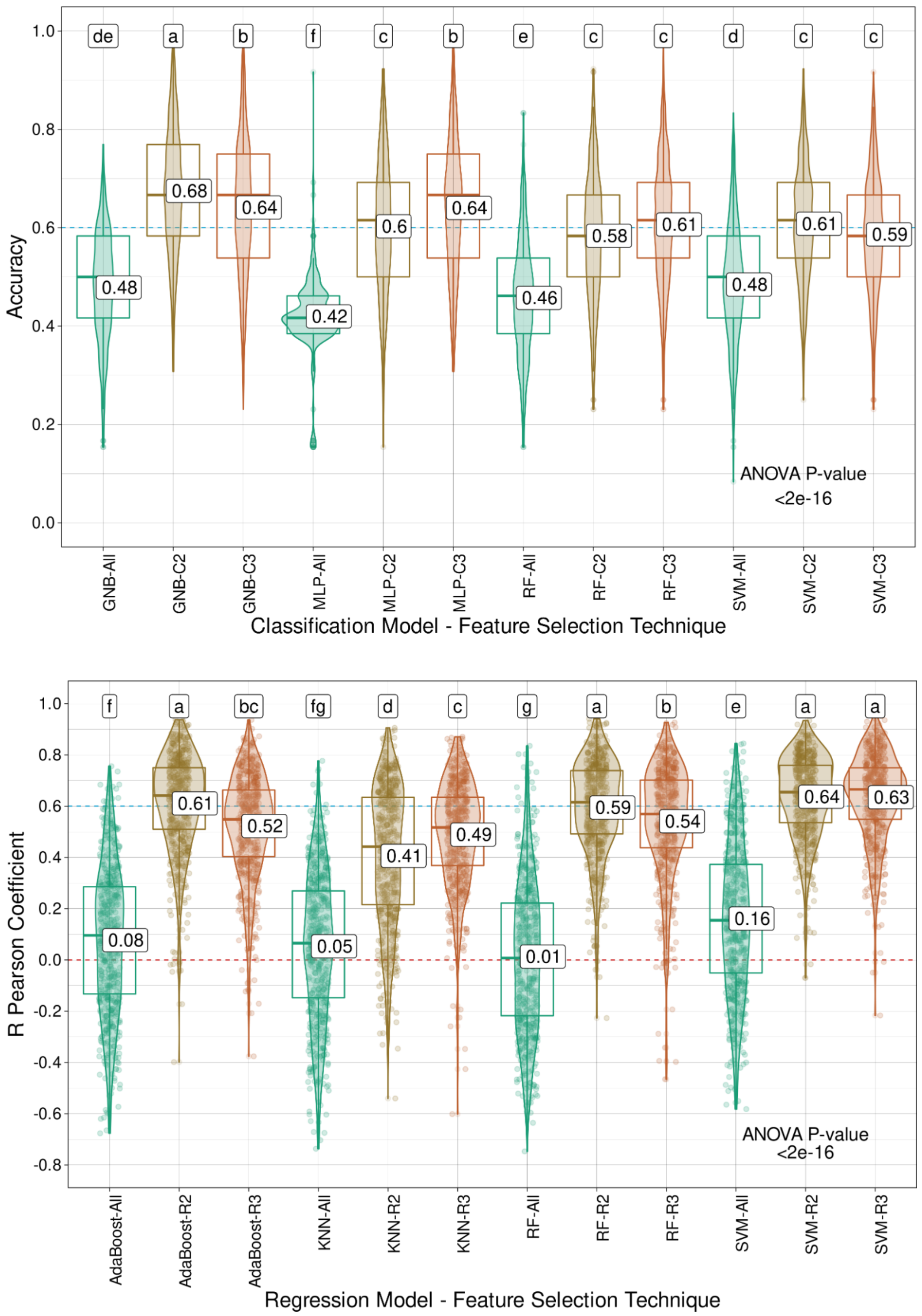


**Supplementary Fig. S24**. Model performance for the *Urochloa decumbens* leaf stem ratio (LSR) using classification (C) algorithms (Gaussian naive Bayes (GNB), multilayer perceptron

(MLP), random forest (RF), and support vector machine (SVM)) and regression (R) models (AdaBoost, k-nearest neighbors (KNN), RF and SVM), coupled with the intersection of the results from the three feature selection (FS) strategies tested (C3/I3), the intersection of at least two out of the three FS methods (C2/I2), or the entire set of markers (All). Mean values are presented in the boxplots, and the labels at the top of the plots represent the results from Tukey's multiple comparisons test (p value of 0.05).

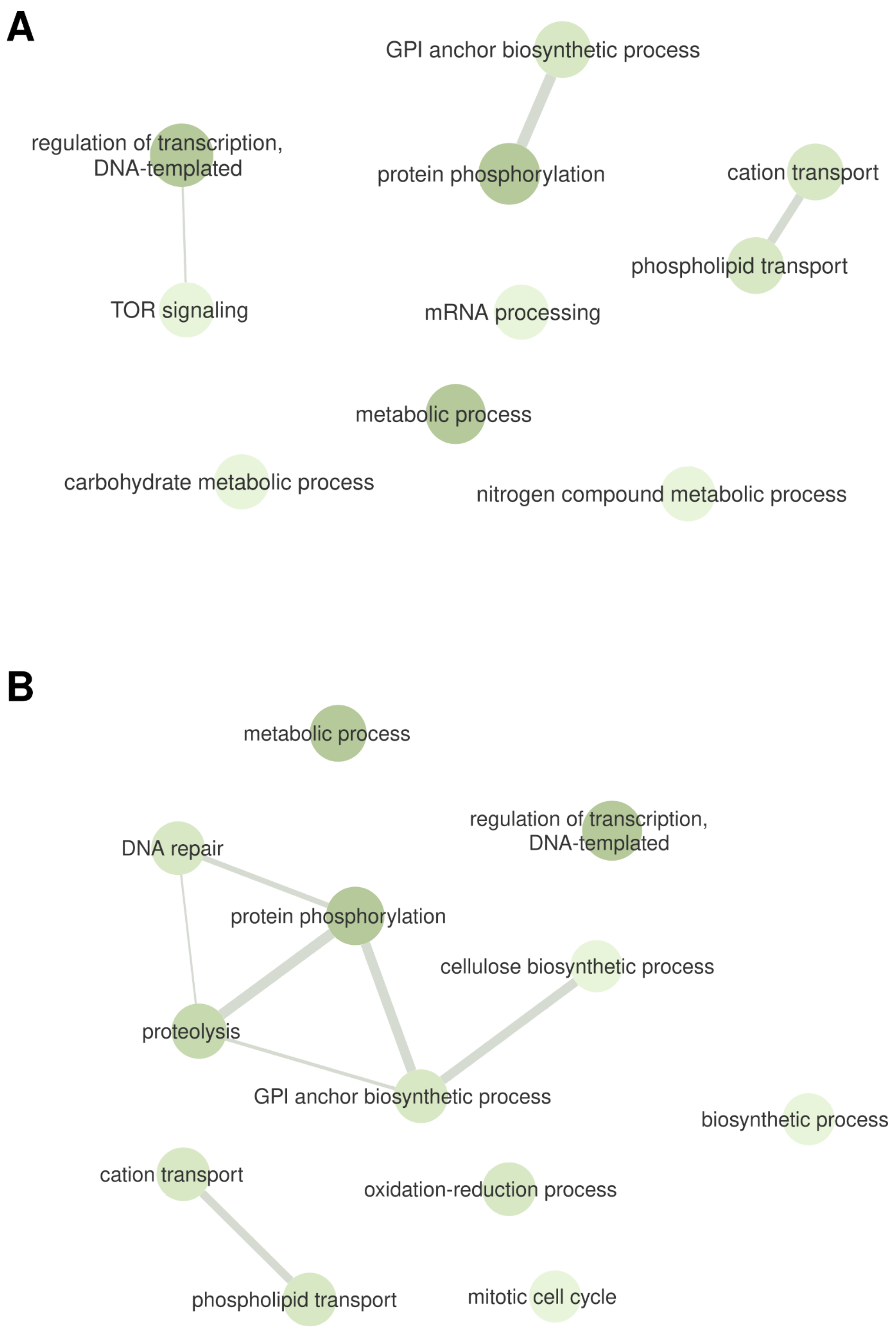


**Supplementary Fig. S25**. Gene ontology (GO) terms from the biological process category associated with the genomic regions containing the markers selected by the intersection of the

results from at least two out of the three feature selection (FS) strategies for the *Urochloa decumbens* leaf stem ratio (LSR). These markers were selected through (**B**) classification algorithms and (**B**) regression models.

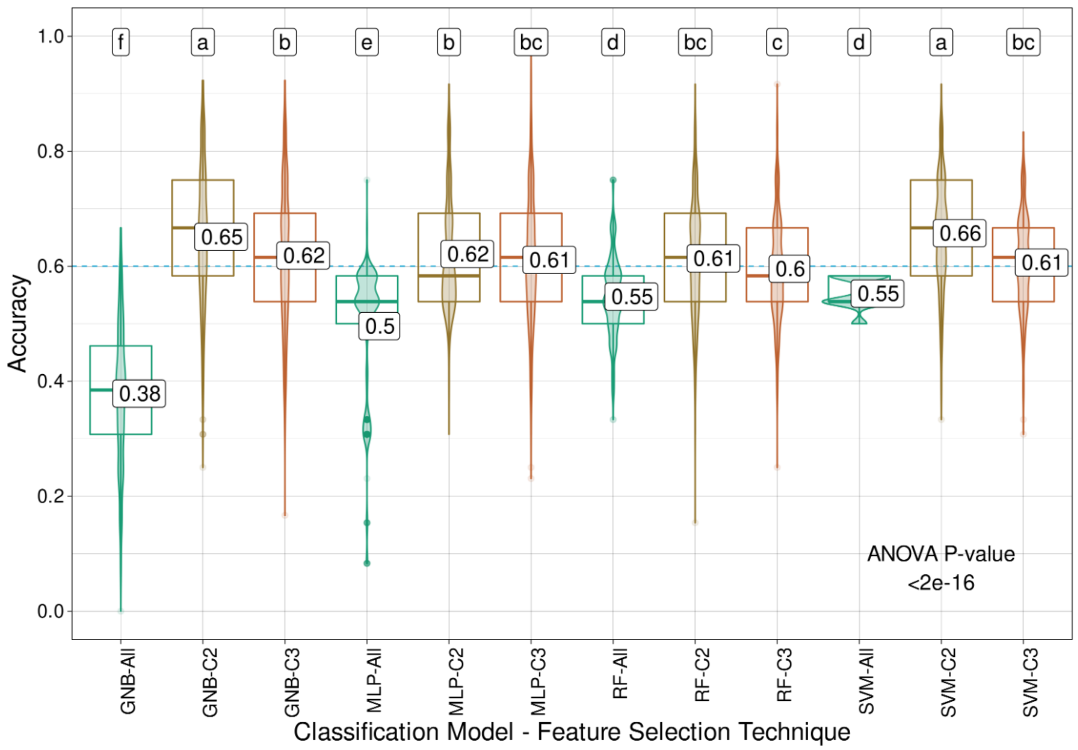
f a b e b bc d bc c d a bc
0.38
0.65
0.62
0.5
0.62
0.61
0.55
0.61
0.6
0.55
0.66
0.61
ANOVA P-value
<2e-16
Accuracy
1.0
0.8
0.6
0.4
0.2
0.0
GNB-All
GNB-C2
GNB-C3
MLP-All
MLP-C2
MLP-C3
RF-All
RF-C2
RF-C3
SVM-All
SVM-C2
SVM-C3
Classification Model - Feature Selection Technique

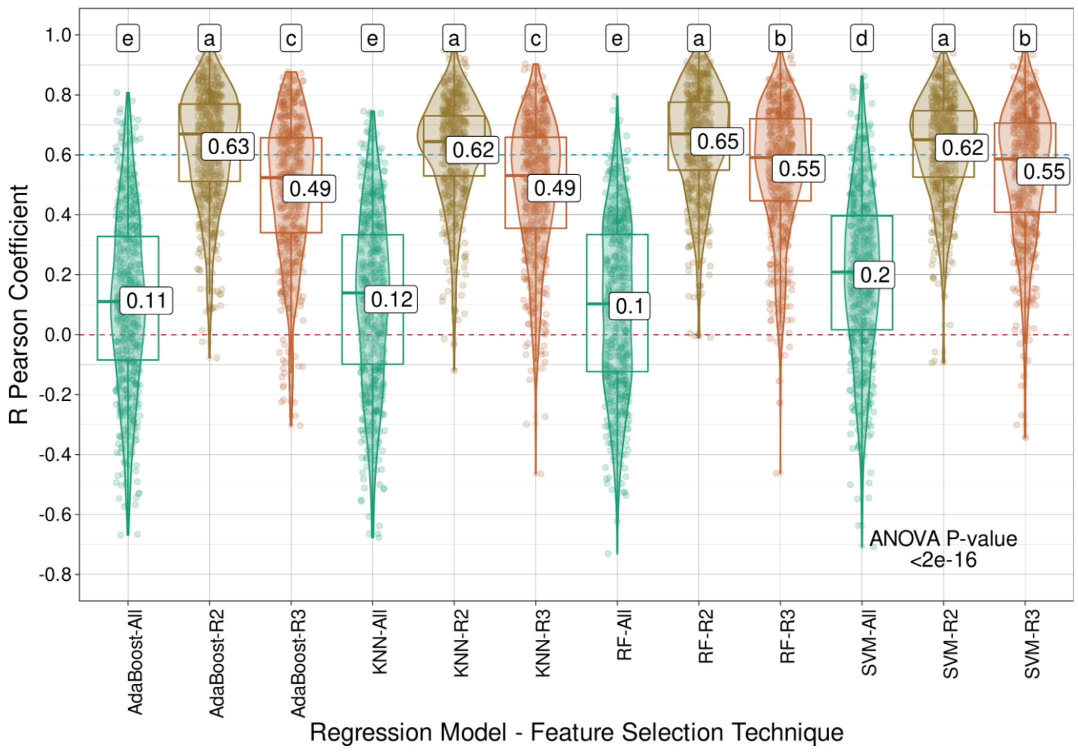
e a c e a c e a b d a b
0.11
0.63
0.49
0.12
0.62
0.49
0.1
0.65
0.55
0.2
0.62
0.55
ANOVA P-value
<2e-16
R Pearson Coefficient
1.0
0.8
0.6
0.4
0.2
0.0
-0.2
-0.4
-0.6
-0.8
AdaBoost-All
AdaBoost-R2
AdaBoost-R3
KNN-All
KNN-R2
KNN-R3
RF-All
RF-R2
RF-R3
SVM-All
SVM-R2
SVM-R3
Regression Model - Feature Selection Technique

**Supplementary Fig. S26**. Model performance for *Urochloa decumbens* regrowth capacity (RC) using classification (C) algorithms (Gaussian naive Bayes (GNB), multilayer perceptron (MLP), random forest (RF), and support vector machine (SVM)) and regression (R) models (AdaBoost, k-nearest neighbors (KNN), RF and SVM), coupled with the intersection of the results from the three feature selection (FS) strategies tested (C3/I3), the intersection of at least two out of the three FS methods (C2/I2), or the entire set of markers (All). Mean values are presented in the boxplots, and the labels at the top of the plots represent the results from Tukey's multiple comparisons test (p value of 0.05).

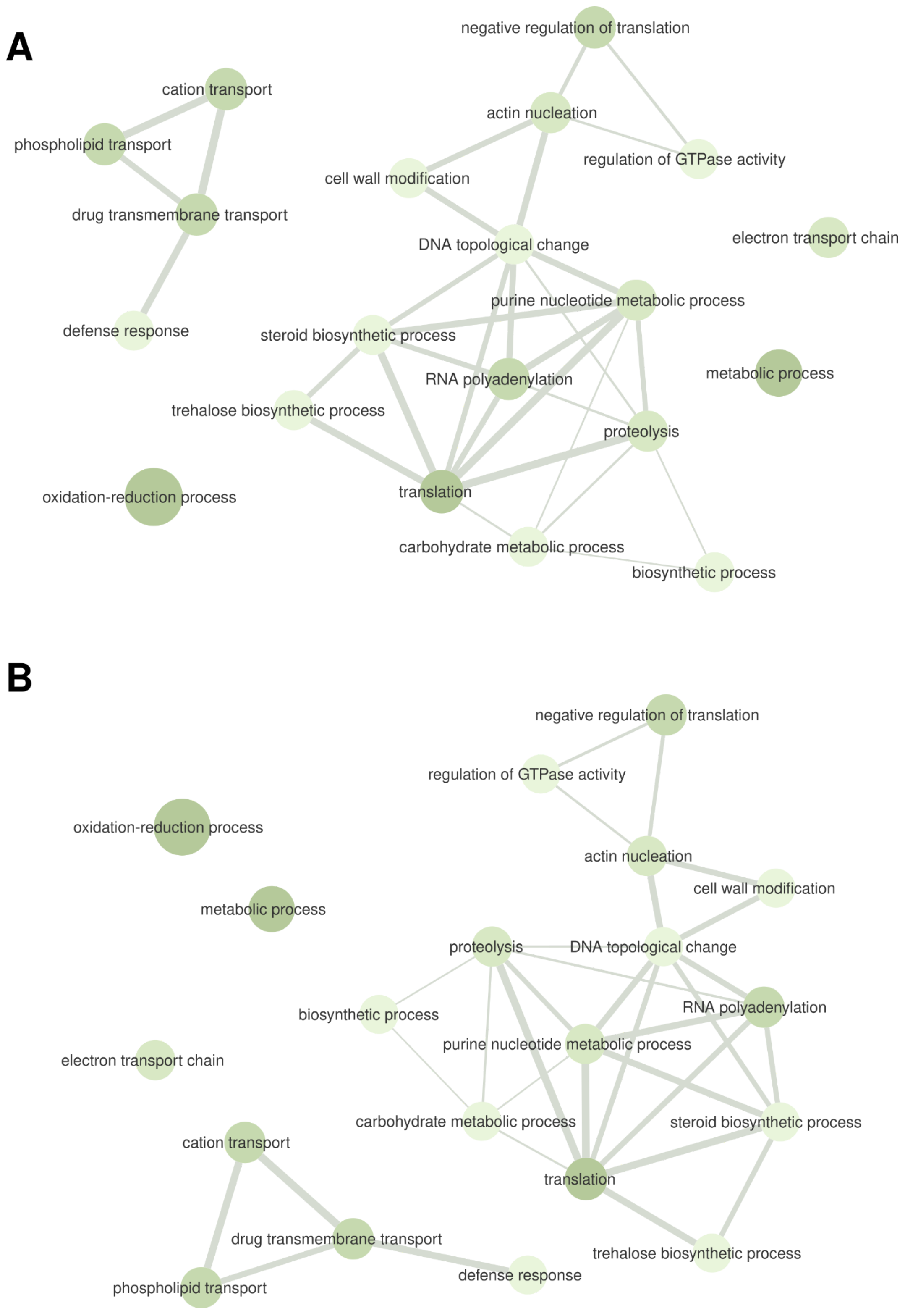
A
negative regulation of translation
cation transport
actin nucleation
phospholipid transport
regulation of GTPase activity
cell wall modification
drug transmembrane transport
electron transport chain
DNA topological change
purine nucleotide metabolic process
defense response
steroid biosynthetic process
metabolic process
RNA polyadenylation
trehalose biosynthetic process
proteolysis
oxidation-reduction process
translation
carbohydrate metabolic process
biosynthetic process
B
negative regulation of translation
regulation of GTPase activity
oxidation-reduction process
actin nucleation
cell wall modification
metabolic process
proteolysis
DNA topological change
RNA polyadenylation
biosynthetic process
purine nucleotide metabolic process
electron transport chain
carbohydrate metabolic process
steroid biosynthetic process
cation transport
translation
drug transmembrane transport
trehalose biosynthetic process
defense response
phospholipid transport

**Supplementary Fig. S27**. Gene ontology (GO) terms from the biological process category associated with the genomic regions containing the markers selected by the intersection of the results from at least two out of the three feature selection (FS) strategies for *Urochloa decumbens* regrowth capacity (RC). These markers were selected through (**A**) classification algorithms and (**B**) regression models.

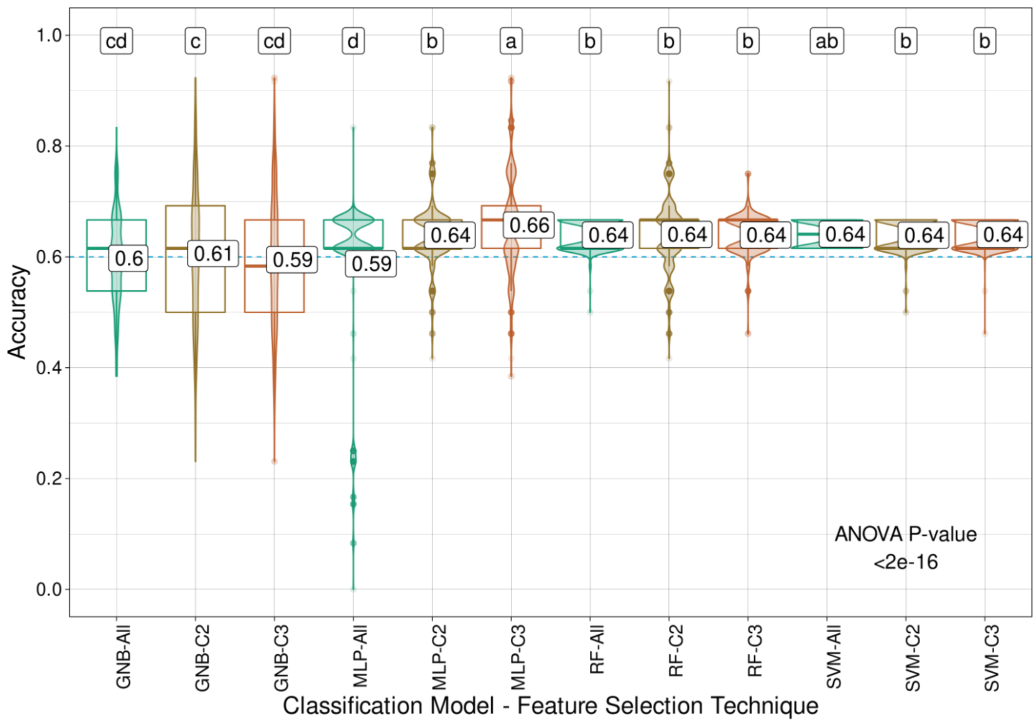


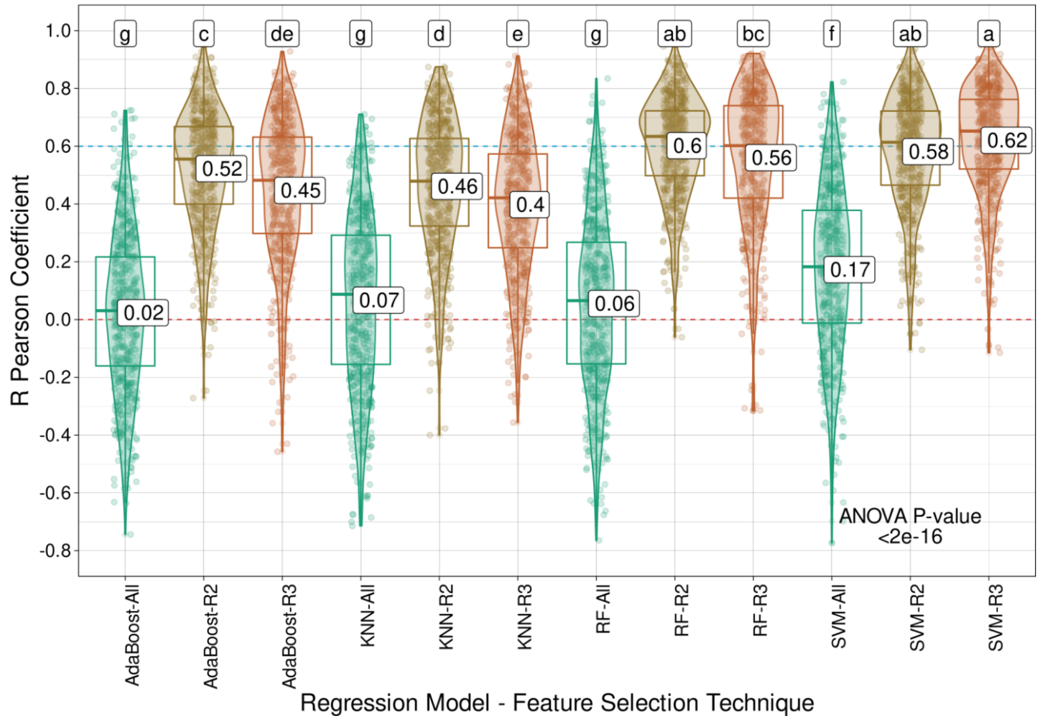


**Supplementary Fig. S28**. Model performance for *Urochloa decumbens* total dry matter (TDM) using classification (C) algorithms (Gaussian naive Bayes (GNB), multilayer perceptron (MLP),

random forest (RF), and support vector machine (SVM)) and regression (R) models (AdaBoost, k-nearest neighbors (KNN), RF and SVM), coupled with the intersection of the results from the three feature selection (FS) strategies tested (C3/I3), the intersection of at least two out of the three FS methods (C2/I2), or the entire set of markers (All). Mean values are presented in the boxplots, and the labels at the top of the plots represent the results from Tukey's multiple comparisons test (p value of 0.05).

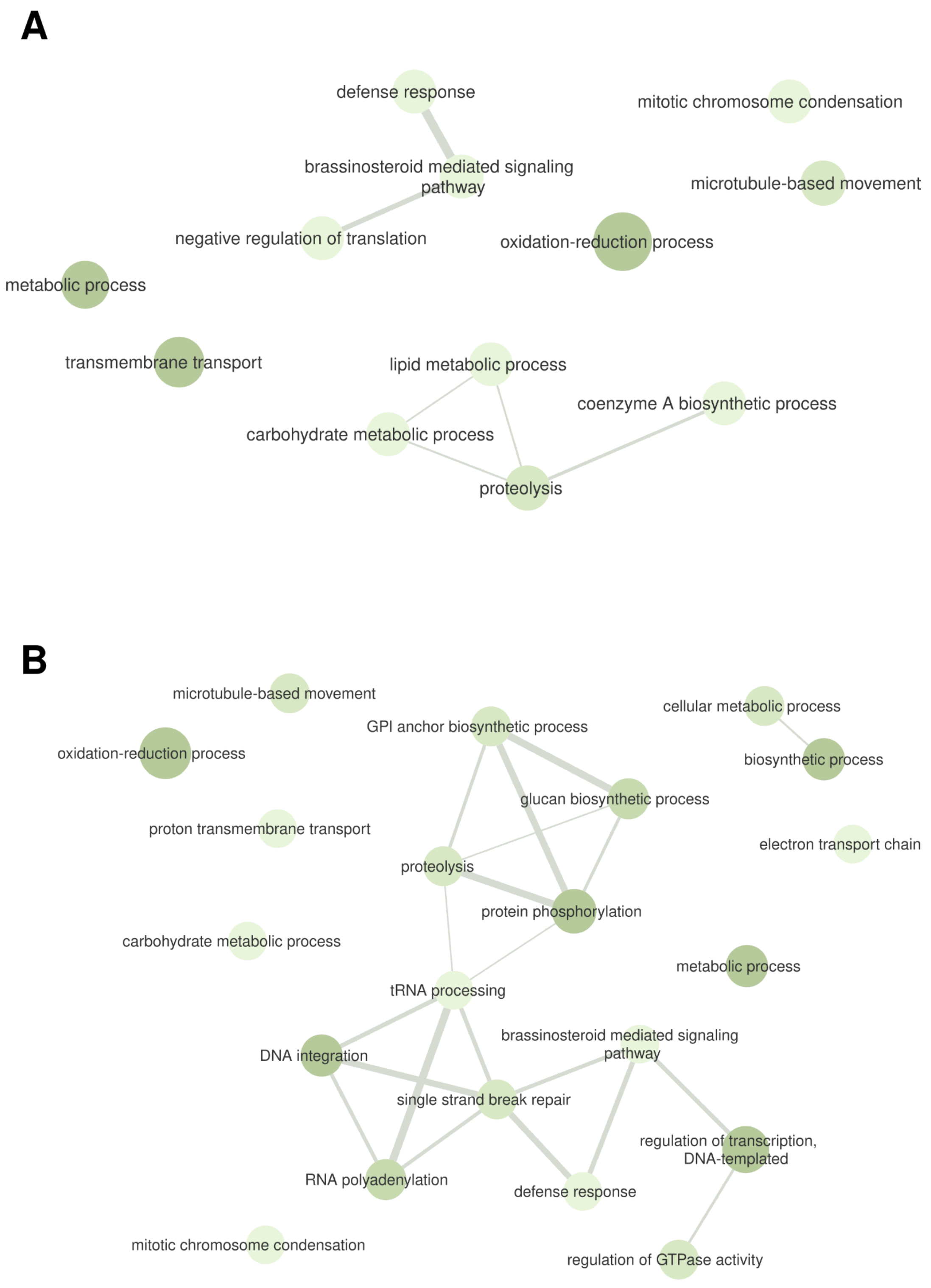


**Supplementary Fig. S29**. Gene ontology (GO) terms from the biological process category associated with the genomic regions containing the markers selected by the intersection of the

results from at least two out of the three feature selection (FS) strategies for *Urochloa decumbens* total dry matter (TDM). These markers were selected through (**A**) classification algorithms and (**B**) regression models.

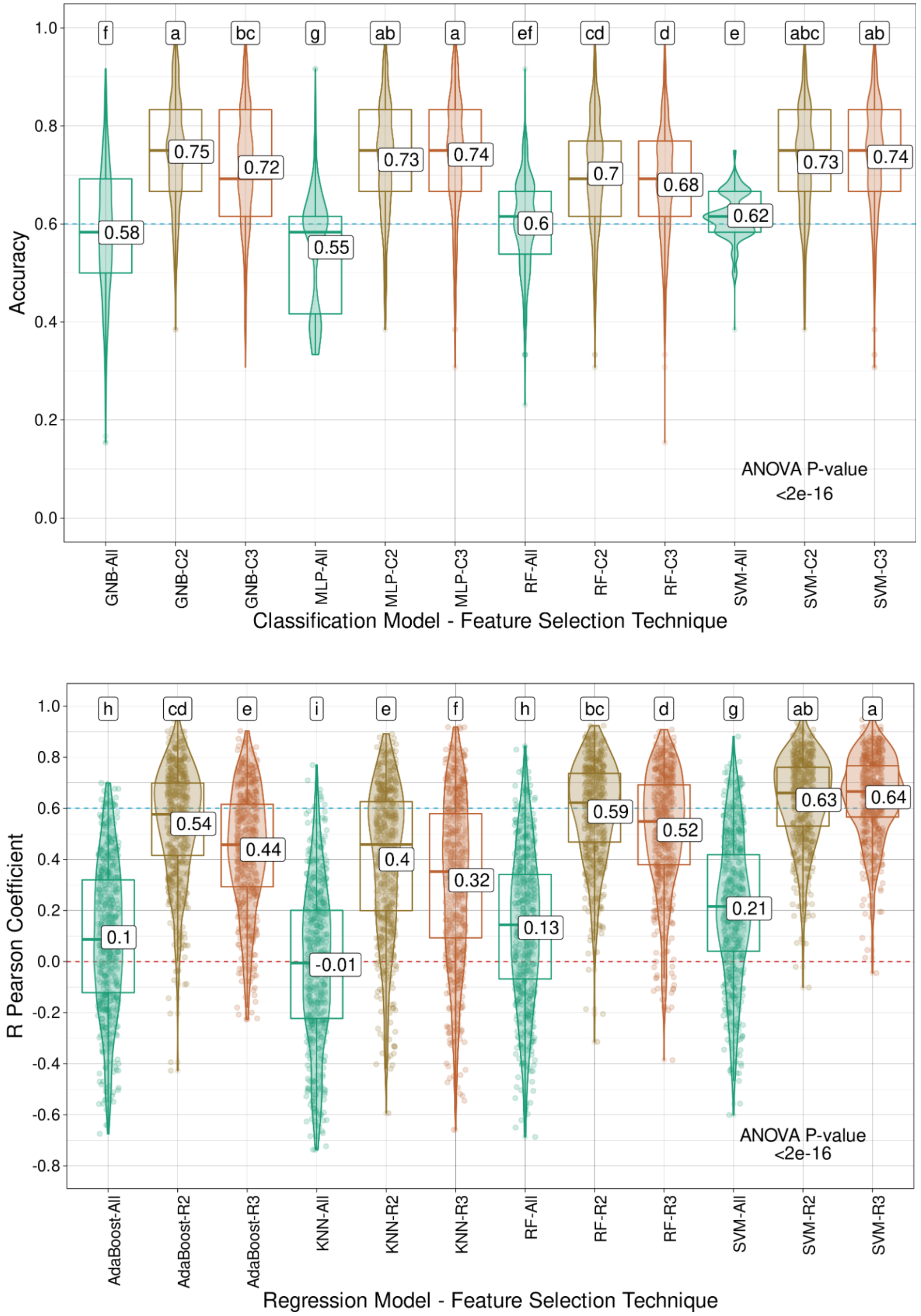


**Supplementary Fig. S30**. Model performance for *Megathyrsus maximus* green matter (GM) using classification (C) algorithms (Gaussian naive Bayes (GNB), multilayer perceptron (MLP),

random forest (RF), and support vector machine (SVM)) and regression (R) models (AdaBoost, k-nearest neighbors (KNN), RF and SVM), coupled with the intersection of the results from the three feature selection (FS) strategies tested (C3/I3), the intersection of at least two out of the three FS methods (C2/I2), or the entire set of markers (All). Mean values are presented in the boxplots, and the labels at the top of the plots represent the results from Tukey's multiple comparisons test (p value of 0.05).

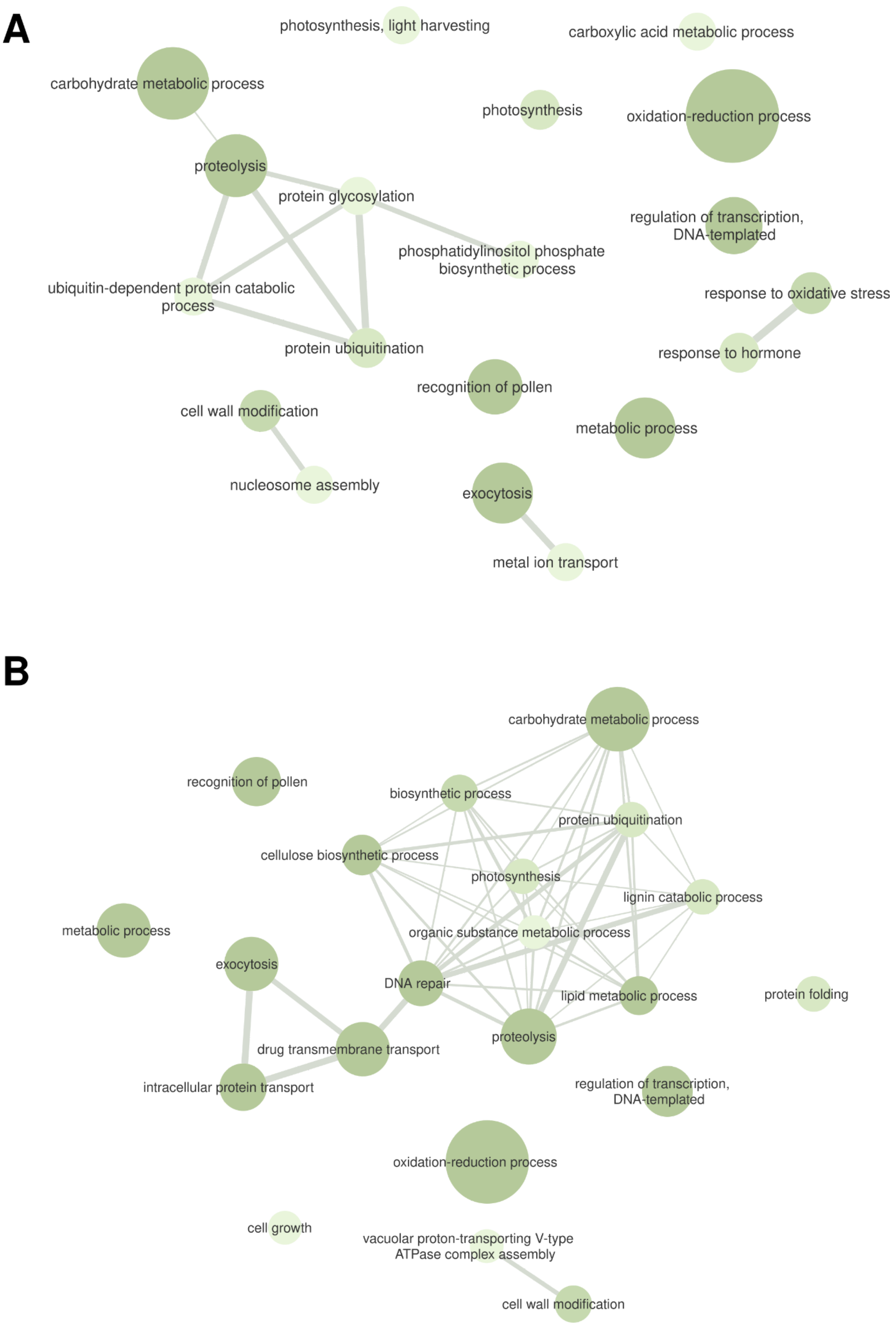
A
photosynthesis, light harvesting
carboxylic acid metabolic process
carbohydrate metabolic process
photosynthesis
oxidation-reduction process
proteolysis
protein glycosylation
regulation of transcription, DNA-templated
phosphatidylinositol phosphate biosynthetic process
ubiquitin-dependent protein catabolic process
response to oxidative stress
protein ubiquitination
response to hormone
recognition of pollen
cell wall modification
metabolic process
nucleosome assembly
exocytosis
metal ion transport
B
carbohydrate metabolic process
recognition of pollen
biosynthetic process
protein ubiquitination
cellulose biosynthetic process
photosynthesis
lignin catabolic process
metabolic process
organic substance metabolic process
exocytosis
DNA repair
lipid metabolic process
protein folding
proteolysis
drug transmembrane transport
intracellular protein transport
regulation of transcription, DNA-templated
oxidation-reduction process
cell growth
vacuolar proton-transporting V-type ATPase complex assembly
cell wall modification

**Supplementary Fig. S31**. Gene ontology (GO) terms from the biological process category associated with the genomic regions containing the markers selected by the intersection of the results from at least two out of the three feature selection (FS) strategies for *Megathyrsus maximus* green matter (GM). These markers were selected through (**A**) classification algorithms and (**B**) regression models.

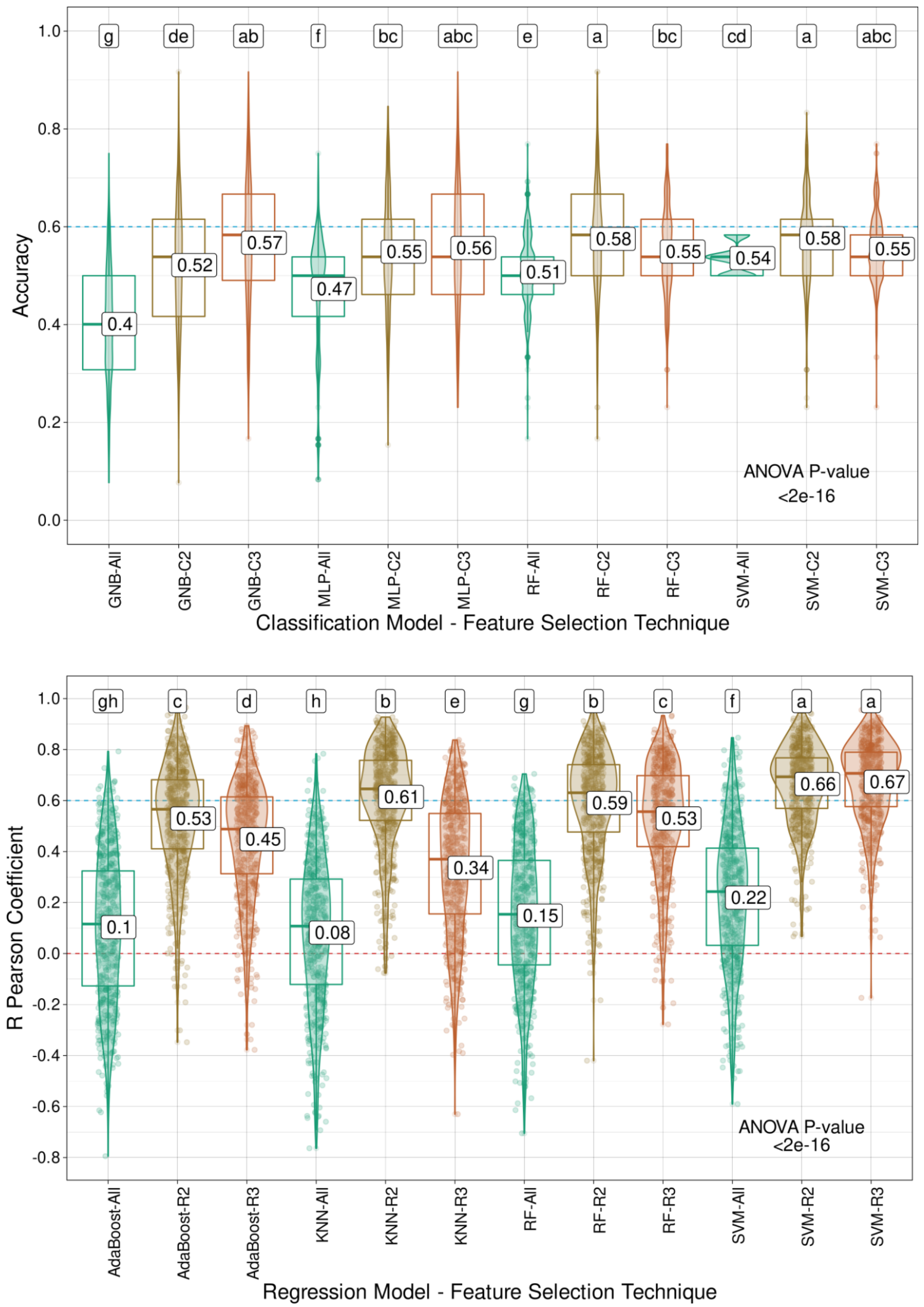


**Supplementary Fig. S32**. Model performance for *Megathyrsus maximus* leaf dry matter (LDM) using classification (C) algorithms (Gaussian naive Bayes (GNB), multilayer perceptron (MLP),

random forest (RF), and support vector machine (SVM)) and regression (R) models (AdaBoost, k-nearest neighbors (KNN), RF and SVM), coupled with the intersection of the results from the three feature selection (FS) strategies tested (C3/I3), the intersection of at least two out of the three FS methods (C2/I2), or the entire set of markers (All). Mean values are presented in the boxplots, and the labels at the top of the plots represent the results from Tukey's multiple comparisons test (p value of 0.05).

A

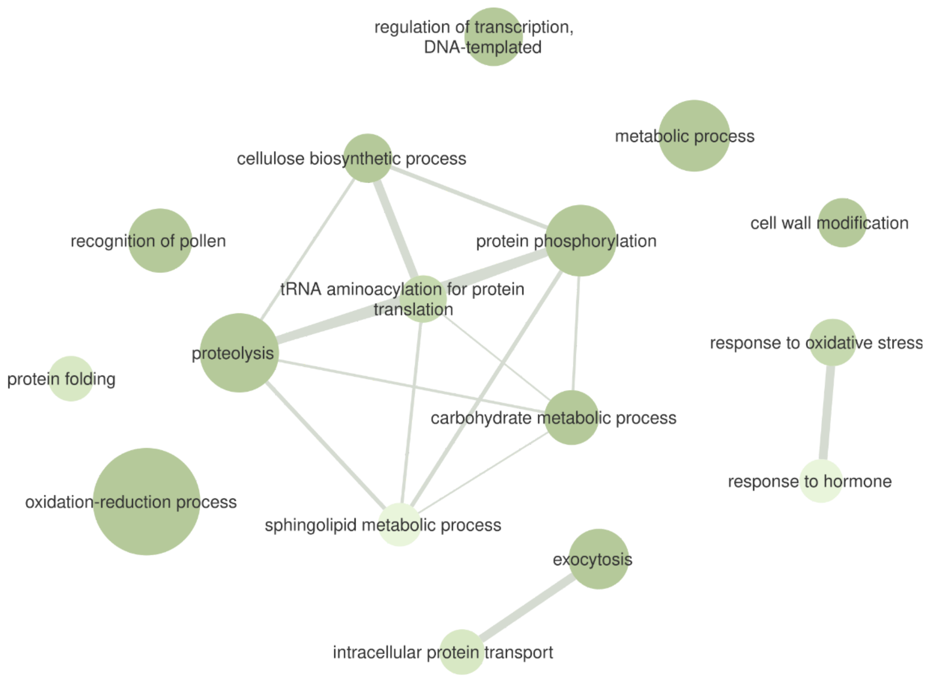
regulation of transcription, DNA-templated
metabolic process
cellulose biosynthetic process
recognition of pollen
protein phosphorylation
cell wall modification
tRNA aminoacylation for protein translation
proteolysis
response to oxidative stress
protein folding
carbohydrate metabolic process
response to hormone
oxidation-reduction process
sphingolipid metabolic process
exocytosis
intracellular protein transport

B

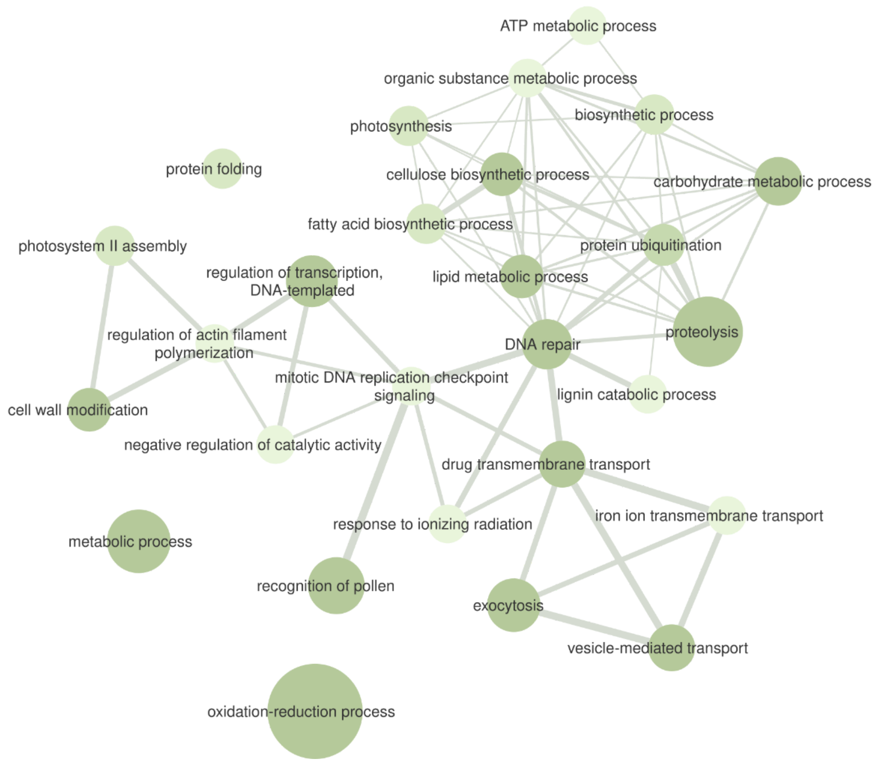
ATP metabolic process
organic substance metabolic process
photosynthesis
biosynthetic process
protein folding
cellulose biosynthetic process
carbohydrate metabolic process
fatty acid biosynthetic process
photosystem II assembly
protein ubiquitination
regulation of transcription, DNA-templated
lipid metabolic process
regulation of actin filament polymerization
proteolysis
DNA repair
mitotic DNA replication checkpoint signaling
lignin catabolic process
cell wall modification
negative regulation of catalytic activity
drug transmembrane transport
iron ion transmembrane transport
response to ionizing radiation
metabolic process
recognition of pollen
exocytosis
vesicle-mediated transport
oxidation-reduction process

**Supplementary Fig. S33**. Gene ontology (GO) terms from the biological process category associated with the genomic regions containing the markers selected by the intersection of the results from at least two out of the three feature selection (FS) strategies for *Megathyrsus maximus* leaf dry matter (LDM). These markers were selected through (**A**) classification algorithms and (**B**) regression models.

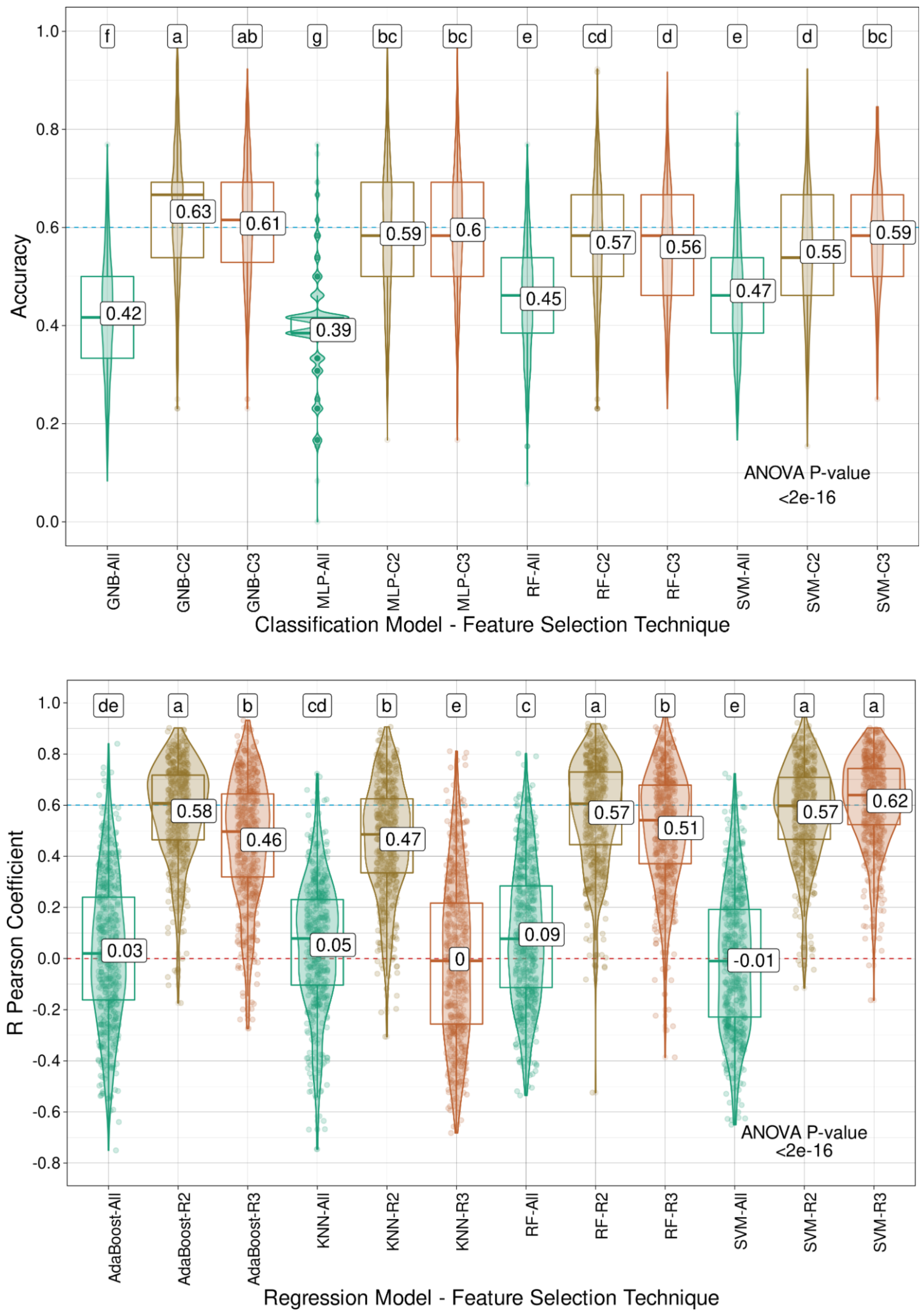


**Supplementary Fig. S34**. Model performance for *Megathyrsus maximus* percentage of leaf blade (PLB) using classification (C) algorithms (Gaussian naive Bayes (GNB), multilayer perceptron (MLP), random forest (RF), and support vector machine (SVM)) and regression (R)

models (AdaBoost, k-nearest neighbors (KNN), RF and SVM), coupled with the intersection of the results from the three feature selection (FS) strategies tested (C3/I3), the intersection of at least two out of the three FS methods (C2/I2), or the entire set of markers (All). Mean values are presented in the boxplots, and the labels at the top of the plots represent the results from Tukey's multiple comparisons test (p value of 0.05).

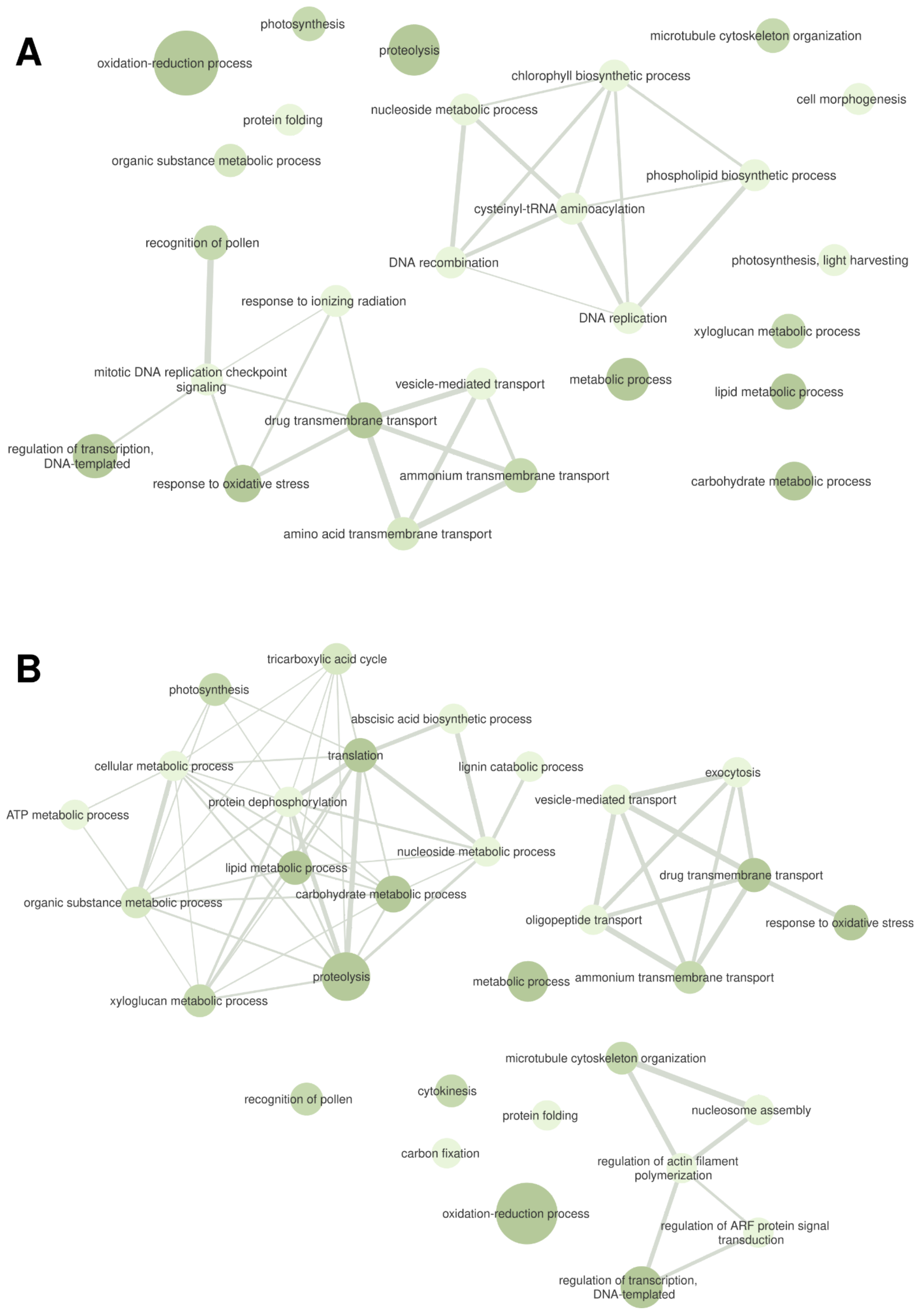


**Supplementary Fig. S35**. Gene ontology (GO) terms from the biological process category associated with the genomic regions containing the markers selected by the intersection of the

results from at least two out of the three feature selection (FS) strategies for *Megathyrsus maximus* percentage of leaf blade (PLB). These markers were selected through (**A**) classification algorithms and (**B**) regression models.

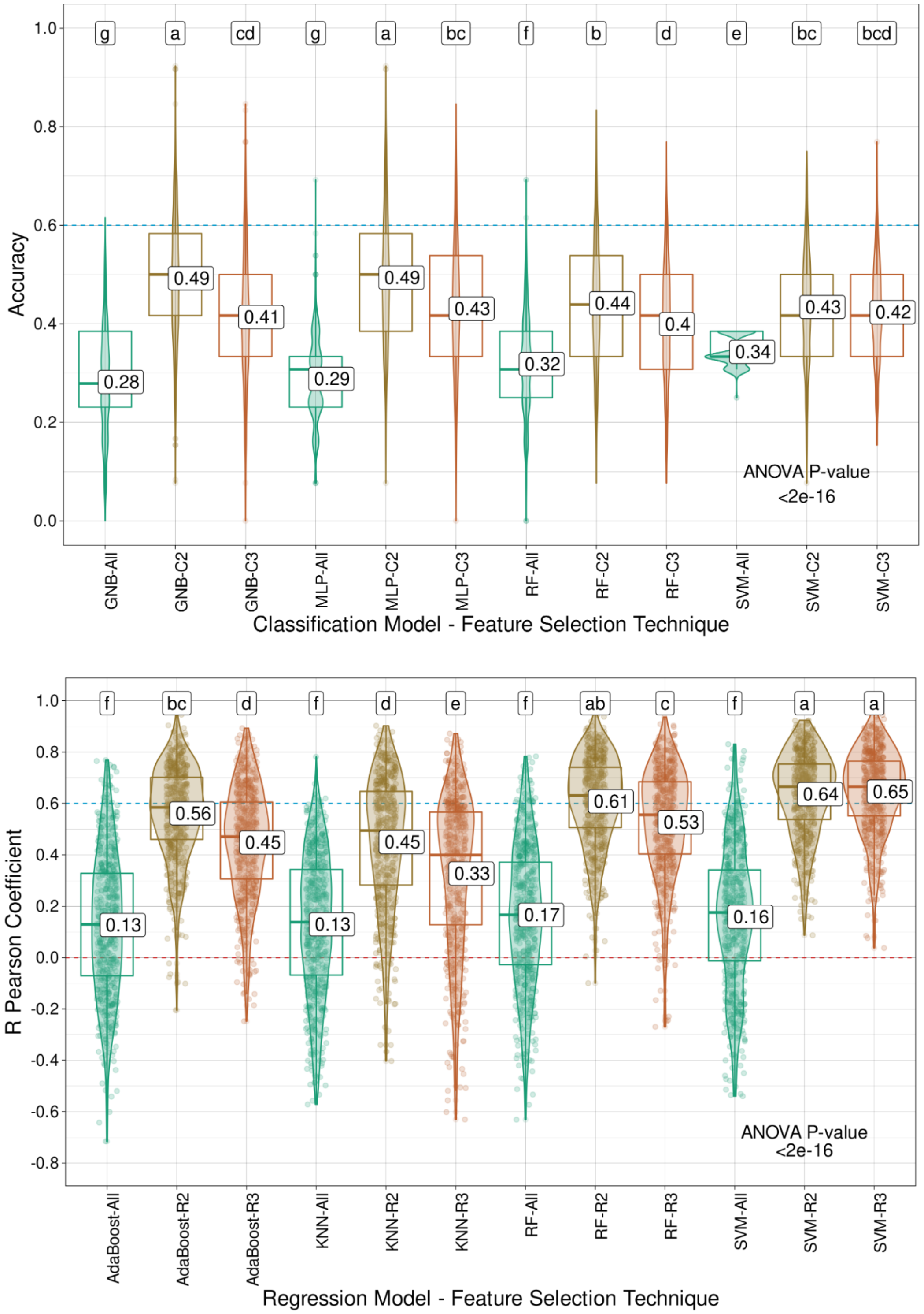


**Supplementary Fig. S36**. Model performance for *Megathyrsus maximus* regrowth capacity (RC) using classification (C) algorithms (Gaussian naive Bayes (GNB), multilayer perceptron (MLP),

random forest (RF), and support vector machine (SVM)) and regression (R) models (AdaBoost, k-nearest neighbors (KNN), RF and SVM), coupled with the intersection of the results from the three feature selection (FS) strategies tested (C3/I3), the intersection of at least two out of the three FS methods (C2/I2), or the entire set of markers (All). Mean values are presented in the boxplots, and the labels at the top of the plots represent the results from Tukey's multiple comparisons test (p value of 0.05).

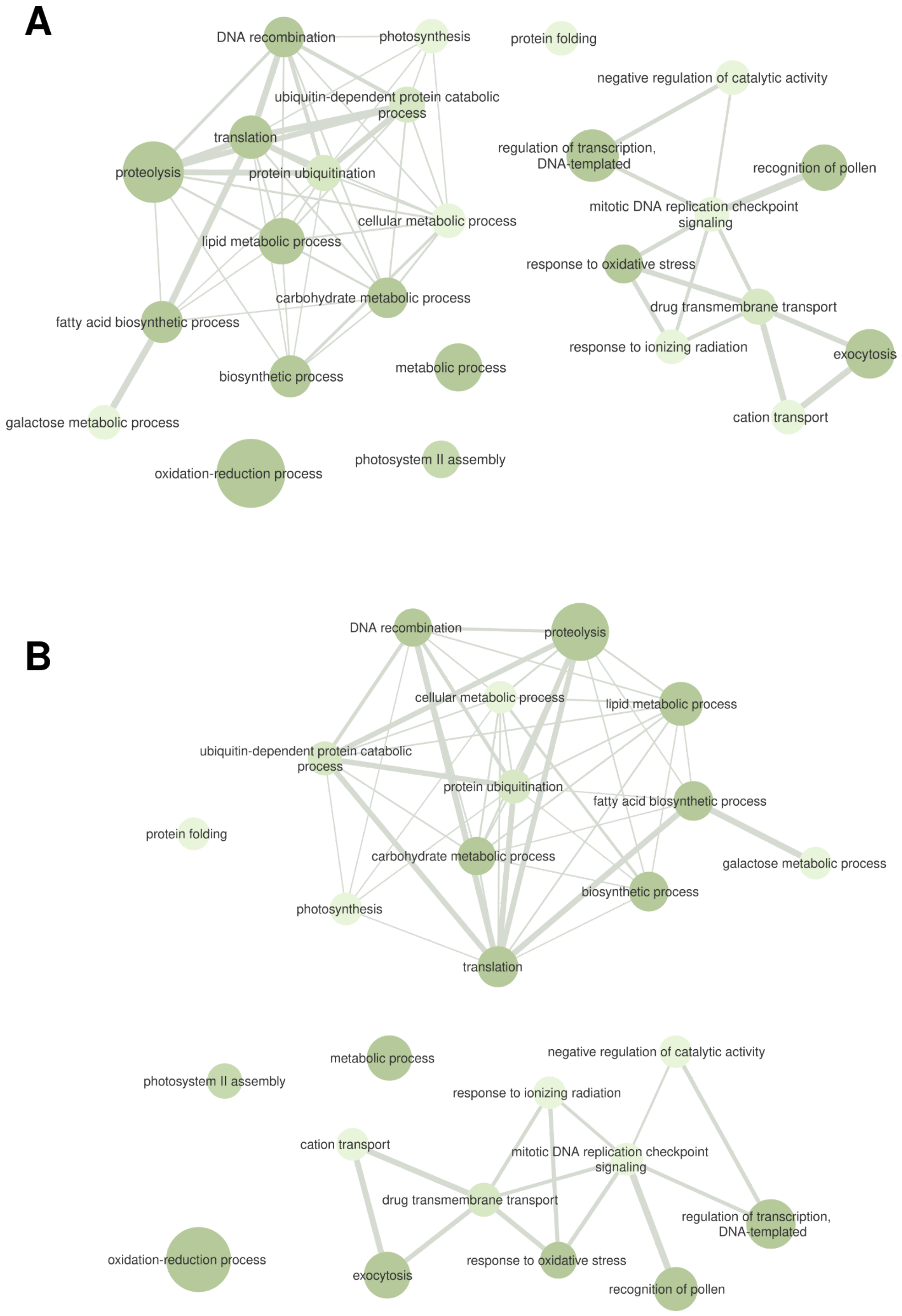
A
DNA recombination
photosynthesis
protein folding
negative regulation of catalytic activity
ubiquitin-dependent protein catabolic process
translation
regulation of transcription, DNA-templated
proteolysis
protein ubiquitination
recognition of pollen
mitotic DNA replication checkpoint signaling
cellular metabolic process
lipid metabolic process
response to oxidative stress
carbohydrate metabolic process
drug transmembrane transport
fatty acid biosynthetic process
response to ionizing radiation
exocytosis
biosynthetic process
metabolic process
cation transport
galactose metabolic process
oxidation-reduction process
photosystem II assembly
B
DNA recombination
proteolysis
cellular metabolic process
lipid metabolic process
ubiquitin-dependent protein catabolic process
protein ubiquitination
fatty acid biosynthetic process
protein folding
carbohydrate metabolic process
galactose metabolic process
biosynthetic process
photosynthesis
translation
metabolic process
negative regulation of catalytic activity
photosystem II assembly
response to ionizing radiation
cation transport
mitotic DNA replication checkpoint signaling
drug transmembrane transport
regulation of transcription, DNA-templated
oxidation-reduction process
exocytosis
response to oxidative stress
recognition of pollen

**Supplementary Fig. S37**. Gene ontology (GO) terms from the biological process category associated with the genomic regions containing the markers selected by the intersection of the results from at least two out of the three feature selection (FS) strategies for *Megathyrsus maximus* regrowth capacity (RC). These markers were selected through (**A**) classification algorithms and (**B**) regression models.

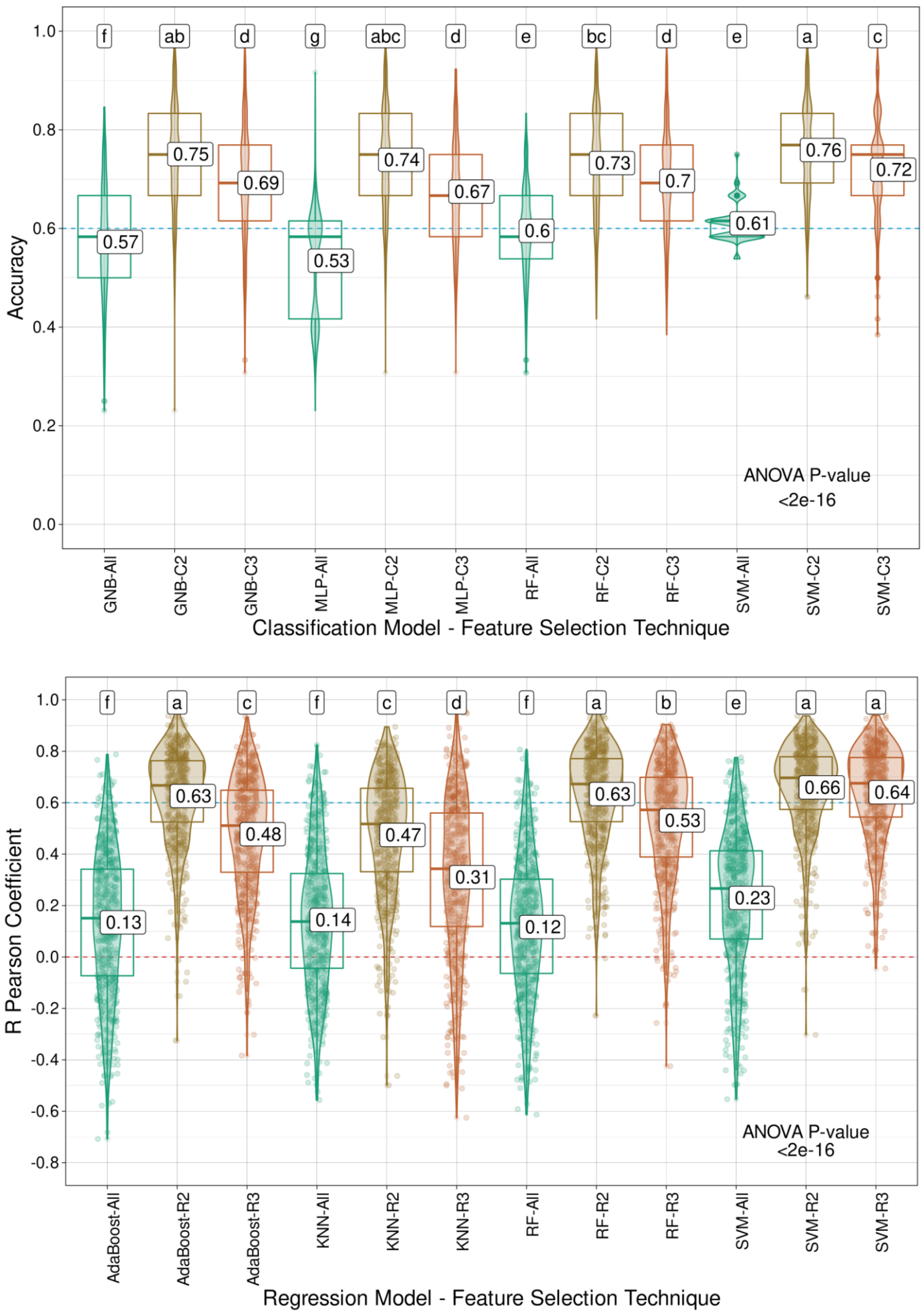


**Supplementary Fig. S38**. Model performance for *Megathyrsus maximus* stem dry matter (SDM) using classification (C) algorithms (Gaussian naive Bayes (GNB), multilayer perceptron (MLP),

random forest (RF), and support vector machine (SVM)) and regression (R) models (AdaBoost, k-nearest neighbors (KNN), RF and SVM), coupled with the intersection of the results from the three feature selection (FS) strategies tested (C3/I3), the intersection of at least two out of the three FS methods (C2/I2), or the entire set of markers (All). Mean values are presented in the boxplots, and the labels at the top of the plots represent the results from Tukey's multiple comparisons test (p value of 0.05).

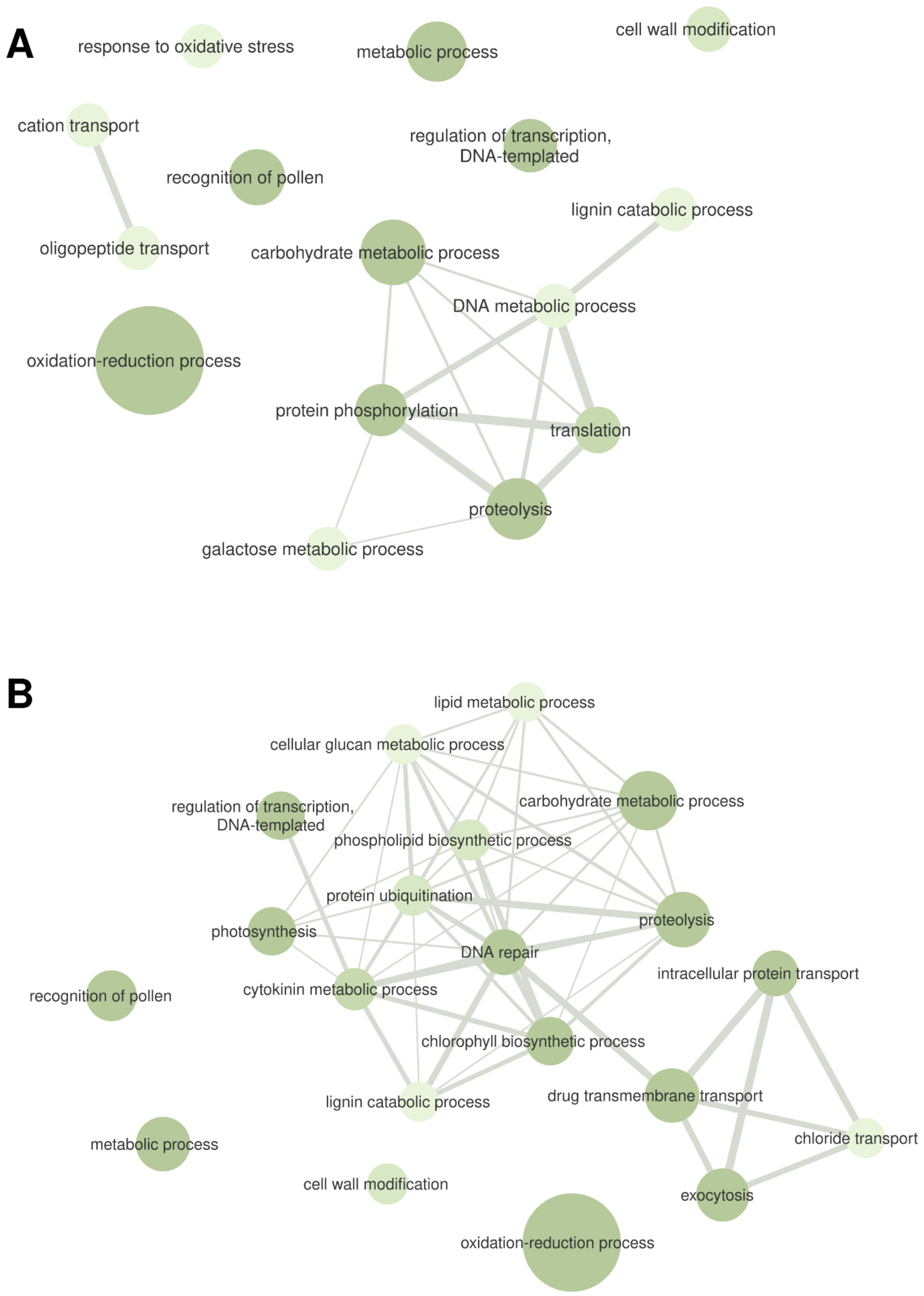


**Supplementary Fig. S39**. Gene ontology (GO) terms from the biological process category associated with the genomic regions containing the markers selected by the intersection of the

results from at least two out of the three feature selection (FS) strategies for *Megathyrsus maximus* stem dry matter (SDM). These markers were selected through (**A**) classification algorithms and (**B**) regression models.

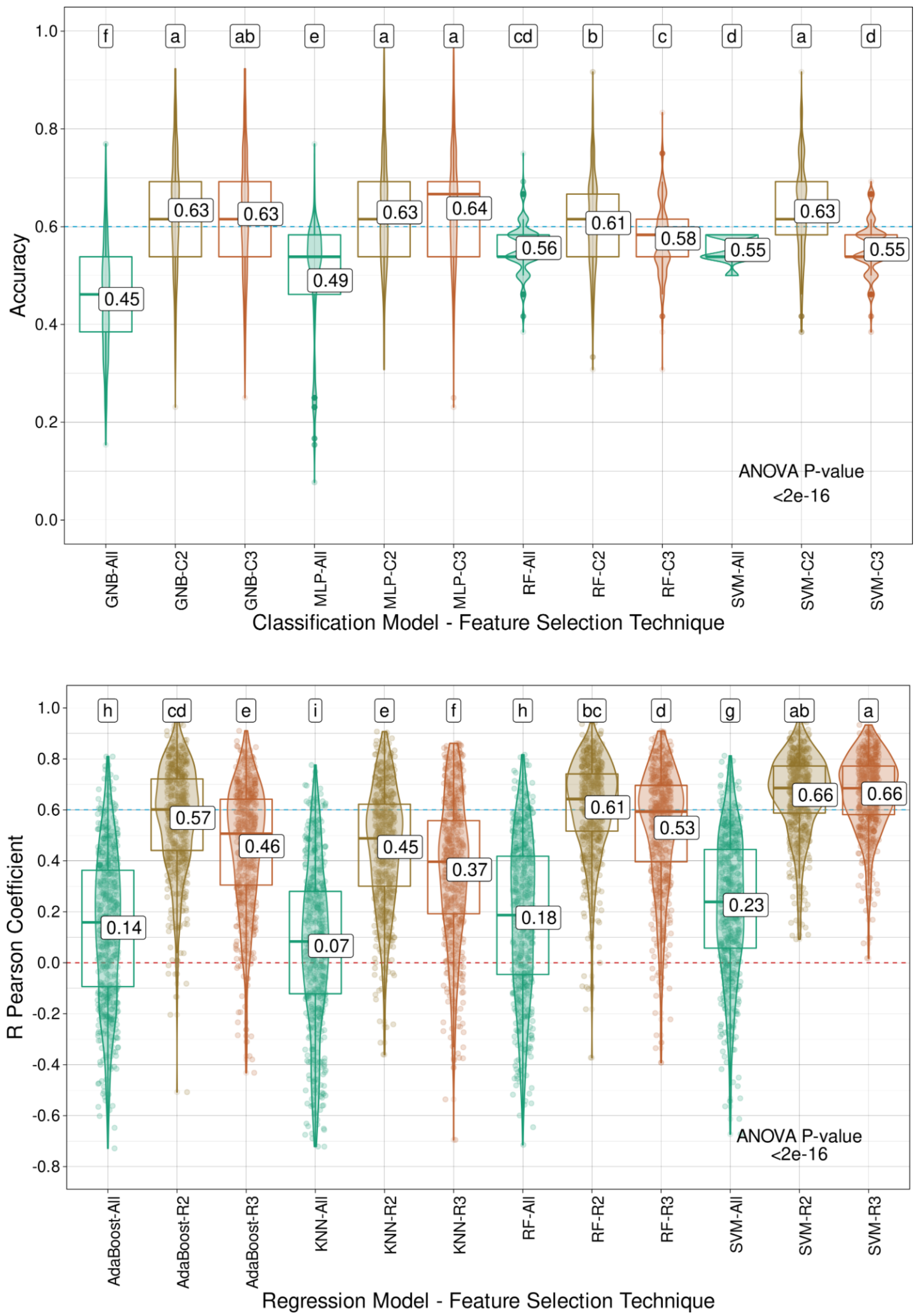


**Supplementary Fig. S40**. Model performance for *Megathyrsus maximus* total dry matter (TDM) using classification (C) algorithms (Gaussian naive Bayes (GNB), multilayer perceptron (MLP),

random forest (RF), and support vector machine (SVM)) and regression (R) models (AdaBoost, k-nearest neighbors (KNN), RF and SVM), coupled with the intersection of the results from the three feature selection (FS) strategies tested (C3/I3), the intersection of at least two out of the three FS methods (C2/I2), or the entire set of markers (All). Mean values are presented in the boxplots, and the labels at the top of the plots represent the results from Tukey's multiple comparisons test (p value of 0.05).

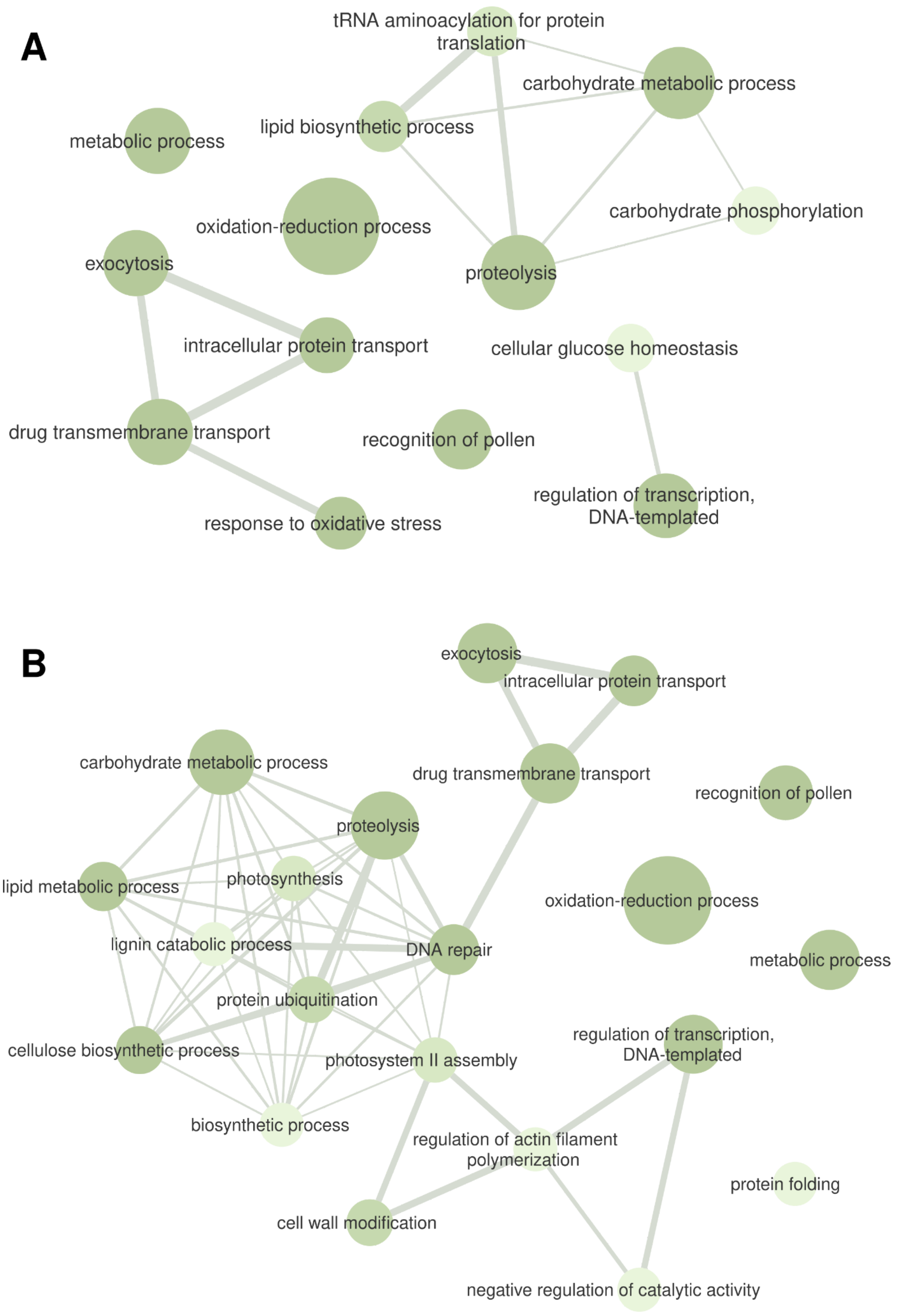
A
tRNA aminoacylation for protein translation
carbohydrate metabolic process
metabolic process
lipid biosynthetic process
oxidation-reduction process
carbohydrate phosphorylation
exocytosis
proteolysis
intracellular protein transport
cellular glucose homeostasis
drug transmembrane transport
recognition of pollen
regulation of transcription, DNA-templated
response to oxidative stress
B
exocytosis
intracellular protein transport
carbohydrate metabolic process
drug transmembrane transport
recognition of pollen
proteolysis
lipid metabolic process
photosynthesis
oxidation-reduction process
lignin catabolic process
DNA repair
metabolic process
protein ubiquitination
regulation of transcription, DNA-templated
cellulose biosynthetic process
photosystem II assembly
biosynthetic process
regulation of actin filament polymerization
protein folding
cell wall modification
negative regulation of catalytic activity

**Supplementary Fig. S41**. Gene ontology (GO) terms from the biological process category associated with the genomic regions containing the markers selected by the intersection of the results from at least two of the three feature selection (FS) strategies for *Megathyrsus maximus* total dry matter (TDM). These markers were selected through (**A**) classification algorithms and (**B**) regression models.

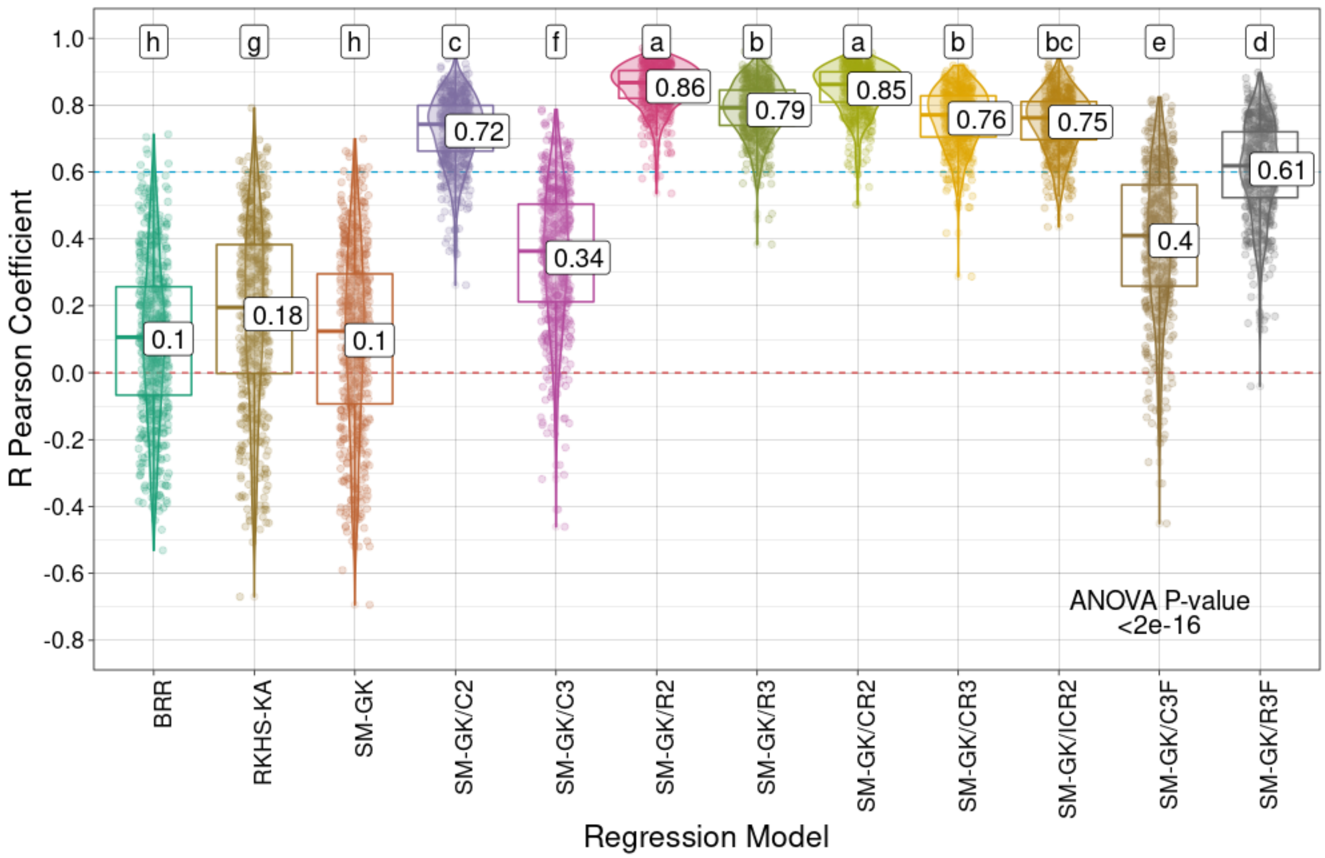

R Pearson Coefficient
1.0
0.8
0.6
0.4
0.2
0.0
-0.2
-0.4
-0.6
-0.8
h
g
h
c
f
a
b
a
b
bc
e
d
0.1
0.18
0.1
0.72
0.34
0.86
0.79
0.85
0.76
0.75
0.4
0.61
ANOVA P-value
<2e-16
BRR
RKHS-KA
SM-GK
SM-GK/C2
SM-GK/C3
SM-GK/R2
SM-GK/R3
SM-GK/CR2
SM-GK/CR3
SM-GK/ICR2
SM-GK/C3F
SM-GK/R3F
Regression Model

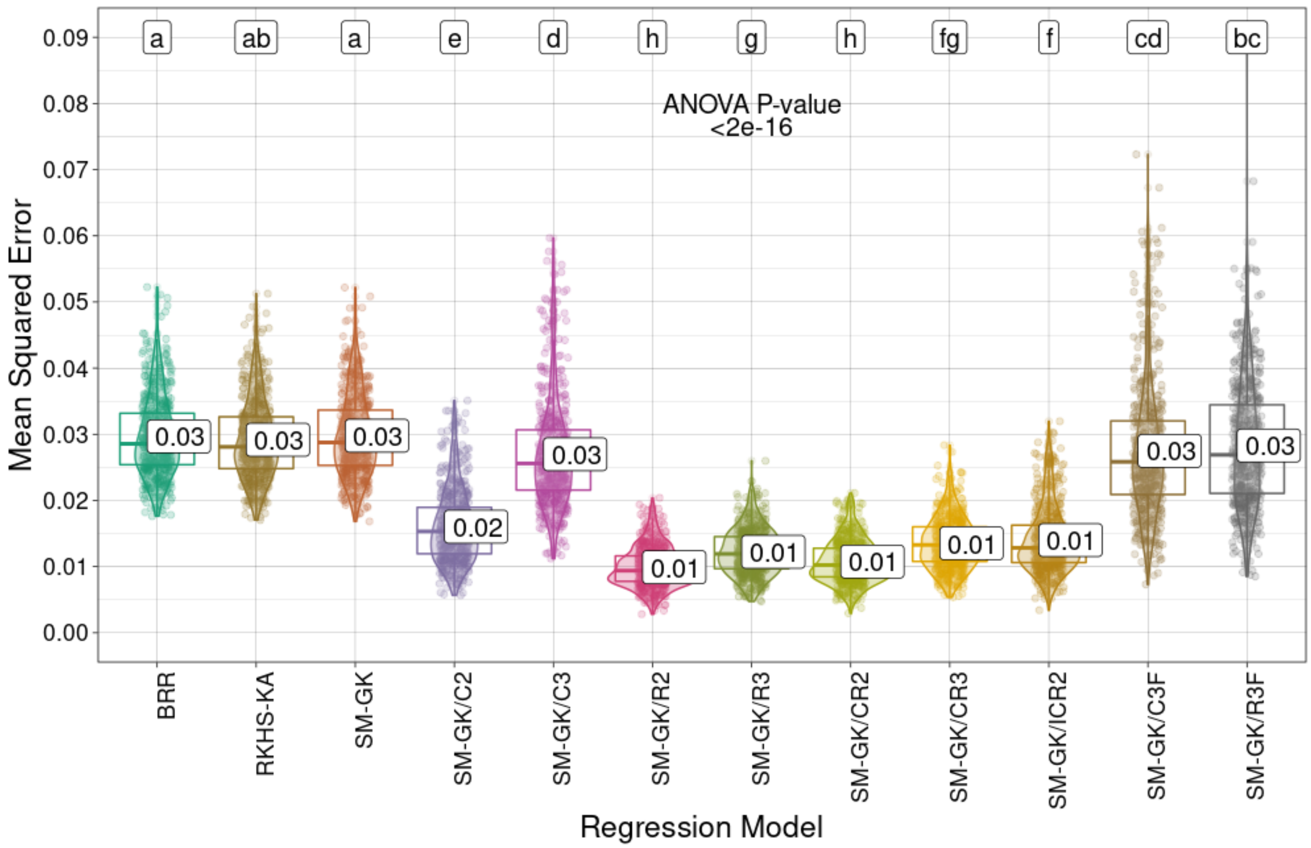


**Supplementary Fig. S42**. Prediction performances (R Pearson correlation coefficients and mean squared errors) for sugarcane stem diameter (SD) using the approaches employed for regression (Bayesian ridge regression (BRR), reproducing kernel Hilbert space with kernel averaging (RKHS-KA) model, and single-environment, main genotypic effect model with a Gaussian kernel (SM-GK)) and the markers selected through feature selection (FS) methods considering classification (C) and regression (R) algorithms and the intersection of the three tested techniques (C3/R3) or at least two of them (C2/R2). The combinations (i) union of C2 and R2 (CR2), (ii) union of C3 and R3 (CR3), and (iii) intersection of C2 and R2 (ICR2) were also evaluated, as well as the addition of C3 and R3 as fixed effects (C3F and R3F). Mean values are

presented in the boxplots, and the labels at the top of the plots represent the results from Tukey's multiple comparisons test (p value of 0.05).

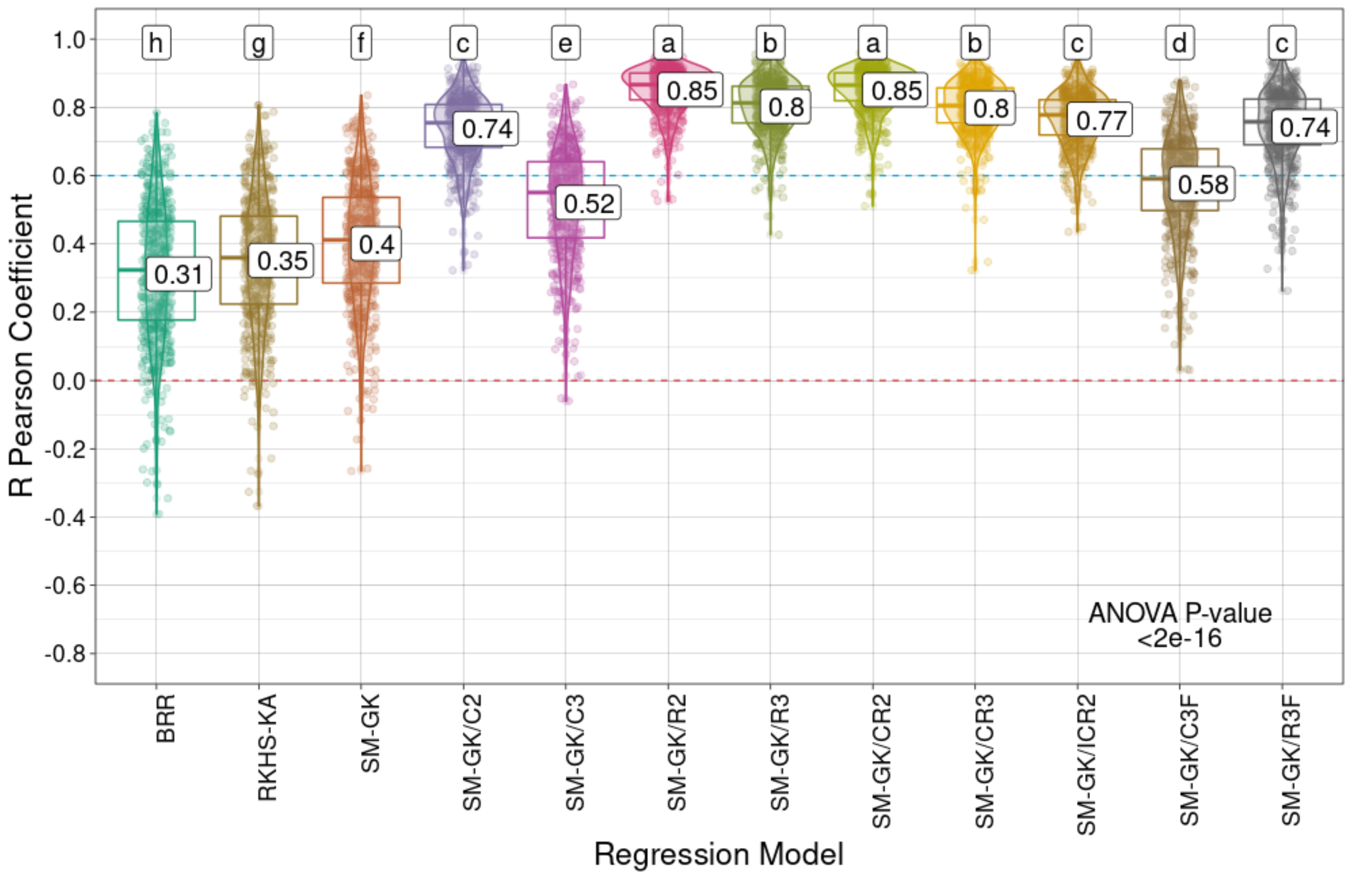

h
g
f
c
e
a
b
a
b
c
d
c
0.31
0.35
0.4
0.74
0.52
0.85
0.8
0.85
0.8
0.77
0.58
0.74
1.0
0.8
0.6
0.4
0.2
0.0
-0.2
-0.4
-0.6
-0.8
R Pearson Coefficient
ANOVA P-value
<2e-16
BRR
RKHS-KA
SM-GK
SM-GK/C2
SM-GK/C3
SM-GK/R2
SM-GK/R3
SM-GK/CR2
SM-GK/CR3
SM-GK/ICR2
SM-GK/C3F
SM-GK/R3F
Regression Model

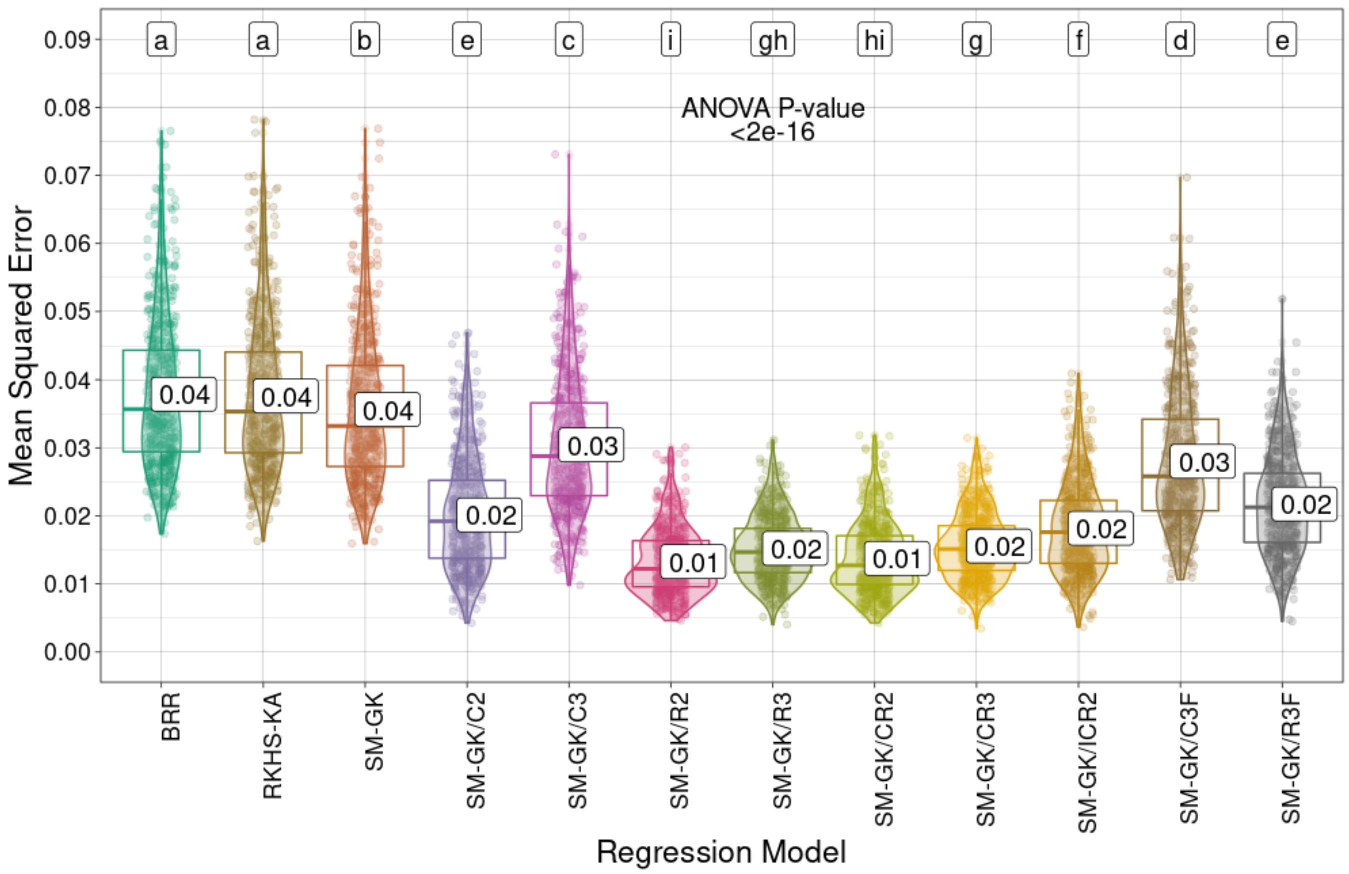


**Supplementary Fig. S43**. Prediction performances (R Pearson correlation coefficients and mean squared errors) for sugarcane stem height (SH) using the approaches employed for regression (Bayesian ridge regression (BRR), reproducing kernel Hilbert space with kernel averaging (RKHS-KA) model, and single-environment, main genotypic effect model with a Gaussian kernel (SM-GK)) and the markers selected through feature selection (FS) methods considering classification (C) and regression (R) algorithms and the intersection of the three tested techniques (C3/R3) or at least two of them (C2/R2). The combinations (i) union of C2 and R2 (CR2), (ii) union of C3 and R3 (CR3), and (iii) intersection of C2 and R2 (ICR2) were also evaluated, as well as the addition of C3 and R3 as fixed effects (C3F and R3F). Mean values are

presented in the boxplots, and the labels at the top of the plots represent the results from Tukey's multiple comparisons test (p value of 0.05).

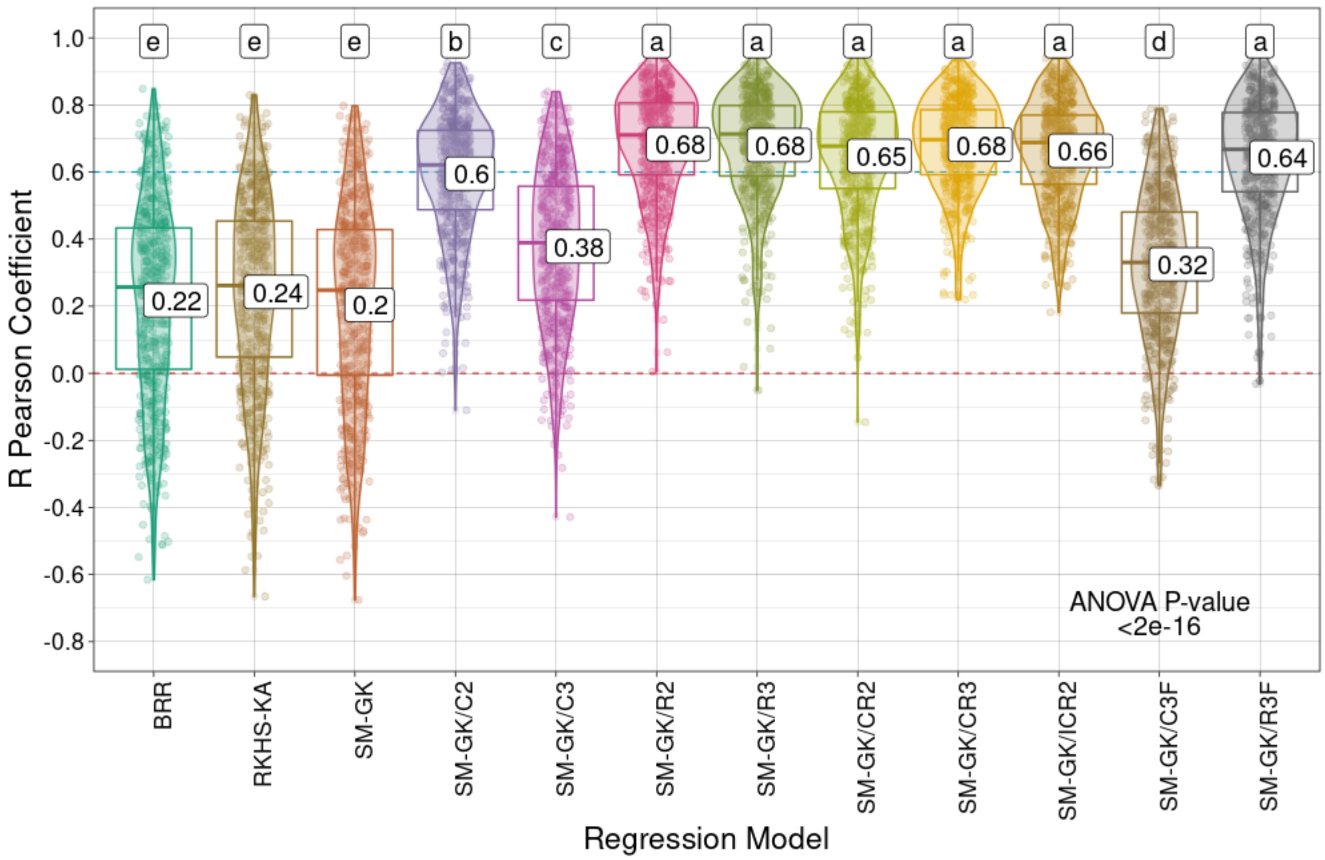

R Pearson Coefficient
1.0
0.8
0.6
0.4
0.2
0.0
-0.2
-0.4
-0.6
-0.8
e
e
e
b
c
a
a
a
a
a
d
a
0.22
0.24
0.2
0.6
0.38
0.68
0.68
0.65
0.68
0.66
0.32
0.64
ANOVA P-value
<2e-16
BRR
RKHS-KA
SM-GK
SM-GK/C2
SM-GK/C3
SM-GK/R2
SM-GK/R3
SM-GK/CR2
SM-GK/CR3
SM-GK/ICR2
SM-GK/C3F
SM-GK/R3F
Regression Model

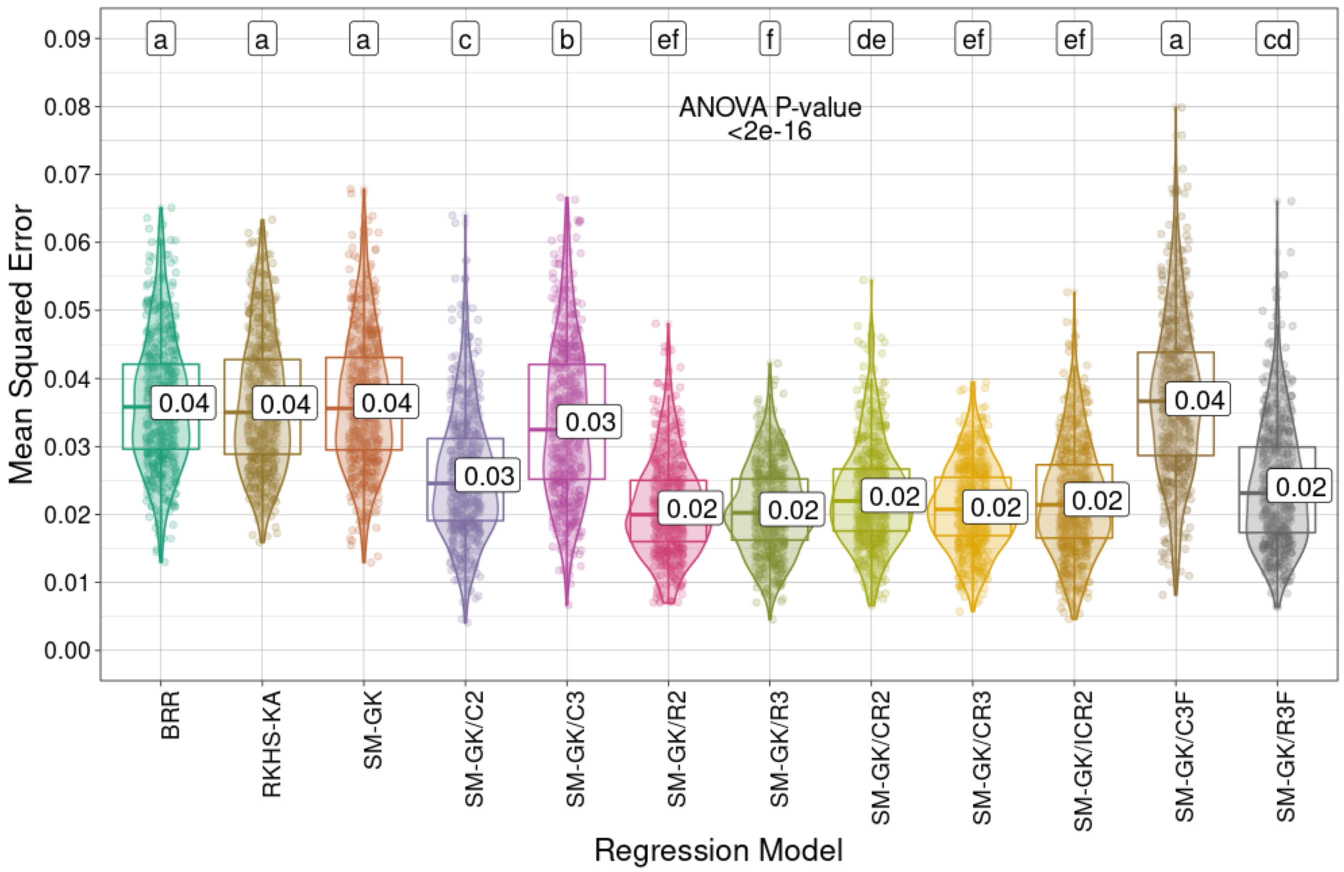


**Supplementary Fig. S44**. Prediction performances (R Pearson correlation coefficients and mean squared errors) for *Urochloa decumbens* field green weight (FGW) using the approaches employed for regression (Bayesian ridge regression (BRR), reproducing kernel Hilbert space with kernel averaging (RKHS-KA) model, and single-environment, main genotypic effect model with a Gaussian kernel (SM-GK)) and the markers selected through feature selection (FS) methods considering classification (C) and regression (R) algorithms and the intersection of the three tested techniques (C3/R3) or at least two of them (C2/R2). The combinations (i) union of C2 and R2 (CR2), (ii) union of C3 and R3 (CR3), and (iii) intersection of C2 and R2 (ICR2) were also evaluated, as well as the addition of C3 and R3 as fixed effects (C3F and R3F). Mean

values are presented in the boxplots, and the labels at the top of the plots represent the results from Tukey's multiple comparisons test (p value of 0.05).

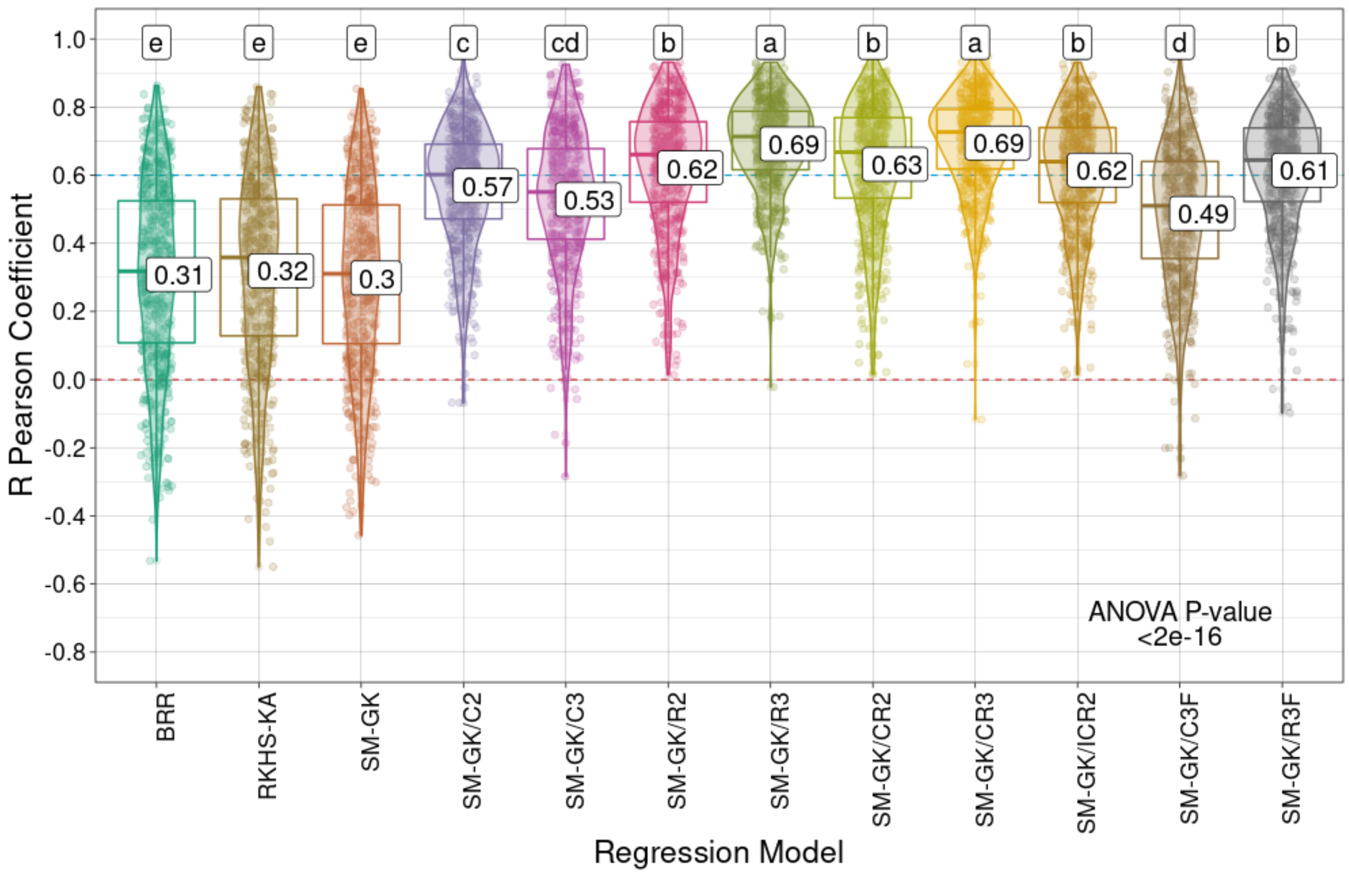

R Pearson Coefficient
1.0
0.8
0.6
0.4
0.2
0.0
-0.2
-0.4
-0.6
-0.8
e
e
e
c
cd
b
a
b
a
b
d
b
0.31
0.32
0.3
0.57
0.53
0.62
0.69
0.63
0.69
0.62
0.49
0.61
ANOVA P-value
<2e-16
BRR
RKHS-KA
SM-GK
SM-GK/C2
SM-GK/C3
SM-GK/R2
SM-GK/R3
SM-GK/CR2
SM-GK/CR3
SM-GK/ICR2
SM-GK/C3F
SM-GK/R3F
Regression Model

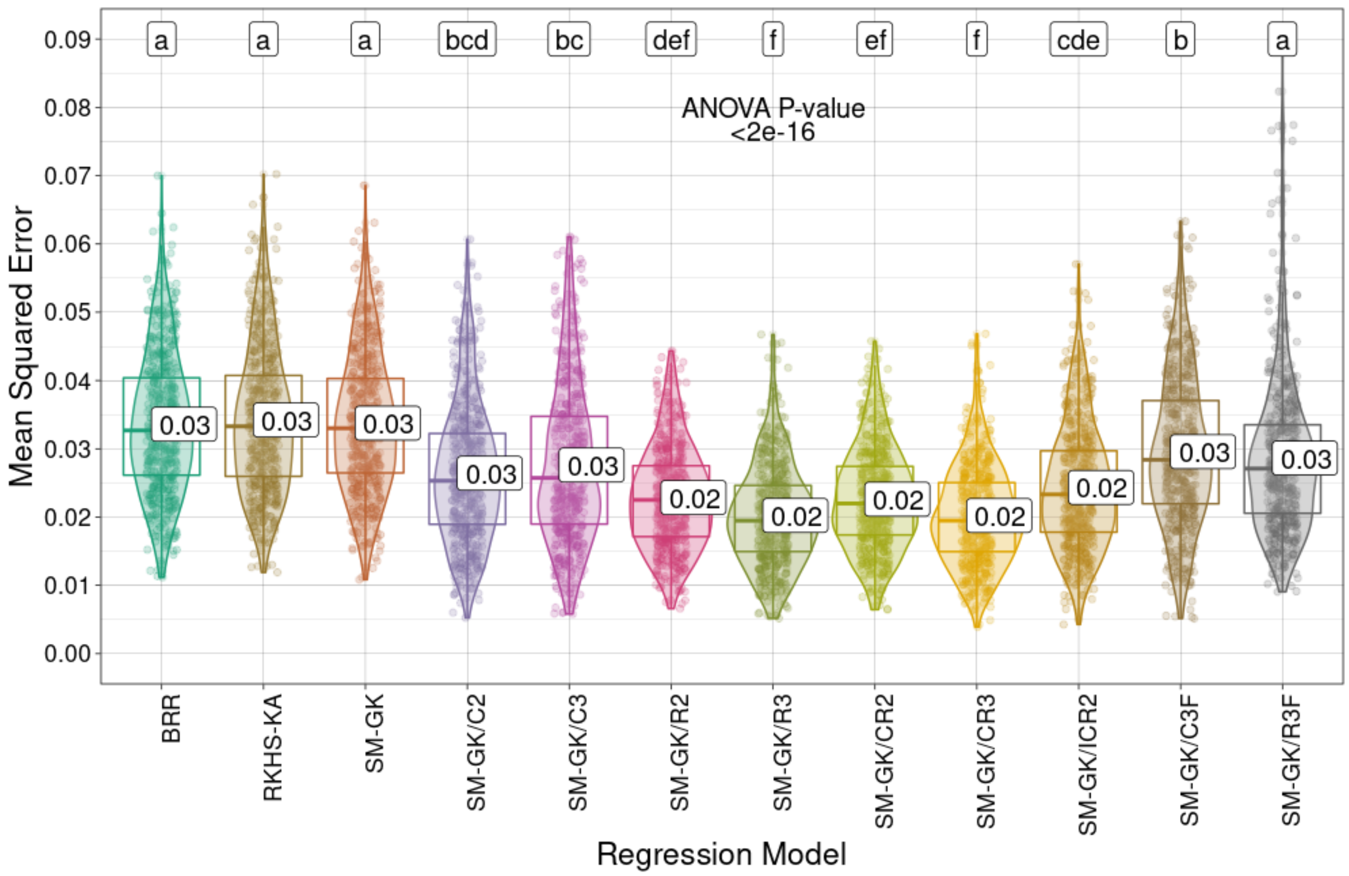


**Supplementary Fig. S45**. Prediction performances (R Pearson correlation coefficients and mean squared errors) for *Urochloa decumbens* leaf dry matter (LDM) using the approaches employed for regression (Bayesian ridge regression (BRR), reproducing kernel Hilbert space with kernel averaging (RKHS-KA) model, and single-environment, main genotypic effect model with a Gaussian kernel (SM-GK)) and the markers selected through feature selection (FS) methods considering classification (C) and regression (R) algorithms and the intersection of the three tested techniques (C3/R3) or at least two of them (C2/R2). The combinations (i) union of C2 and R2 (CR2), (ii) union of C3 and R3 (CR3), and (iii) intersection of C2 and R2 (ICR2) were also evaluated, as well as the addition of C3 and R3 as fixed effects (C3F and R3F). Mean values are

presented in the boxplots, and the labels at the top of the plots represent the results from Tukey's multiple comparisons test (p value of 0.05).

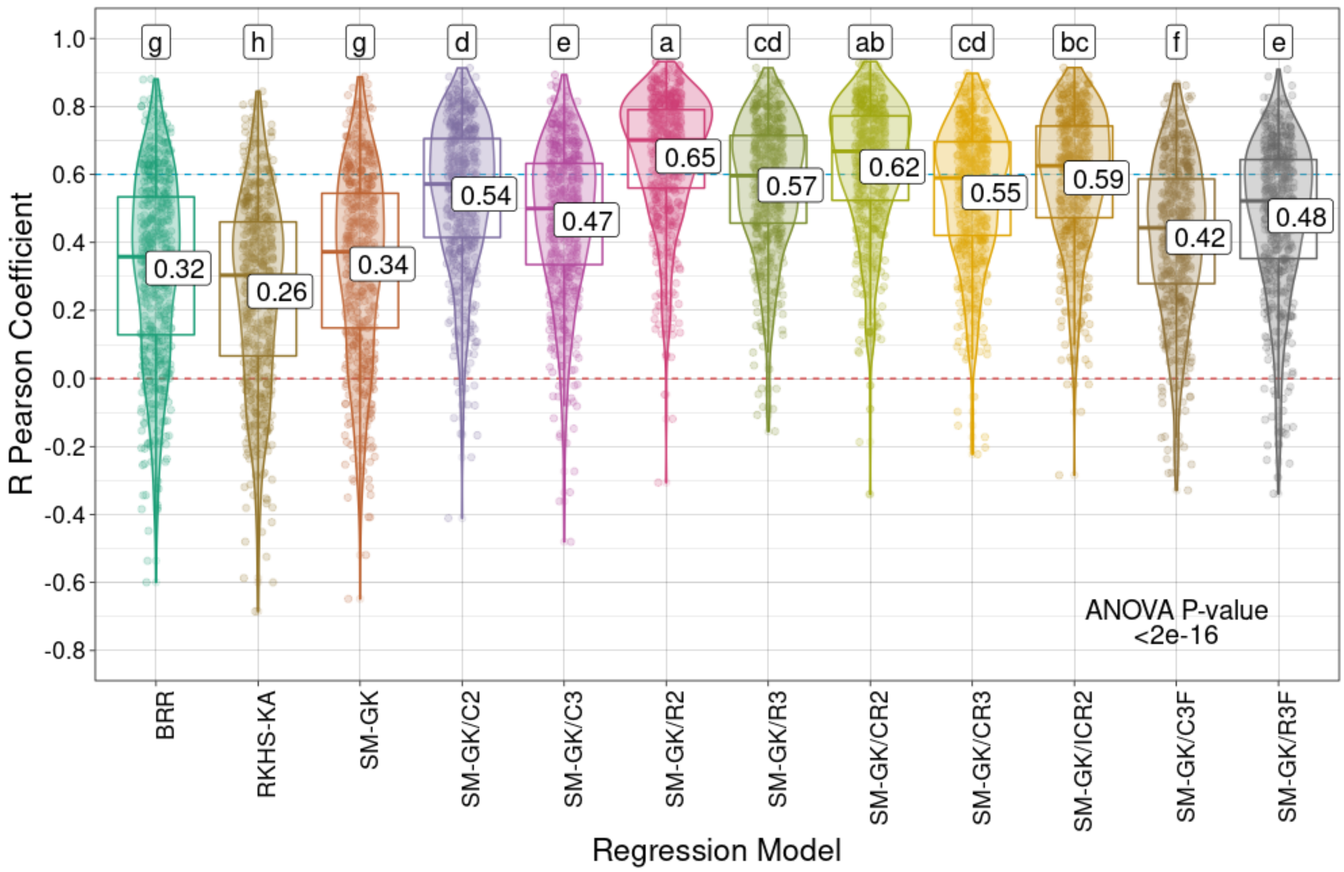
R Pearson Coefficient
1.0
0.8
0.6
0.4
0.2
0.0
-0.2
-0.4
-0.6
-0.8
g
h
g
d
e
a
cd
ab
cd
bc
f
e
0.32
0.26
0.34
0.54
0.47
0.65
0.57
0.62
0.55
0.59
0.42
0.48
ANOVA P-value
<2e-16
BRR
RKHS-KA
SM-GK
SM-GK/C2
SM-GK/C3
SM-GK/R2
SM-GK/R3
SM-GK/CR2
SM-GK/CR3
SM-GK/ICR2
SM-GK/C3F
SM-GK/R3F
Regression Model

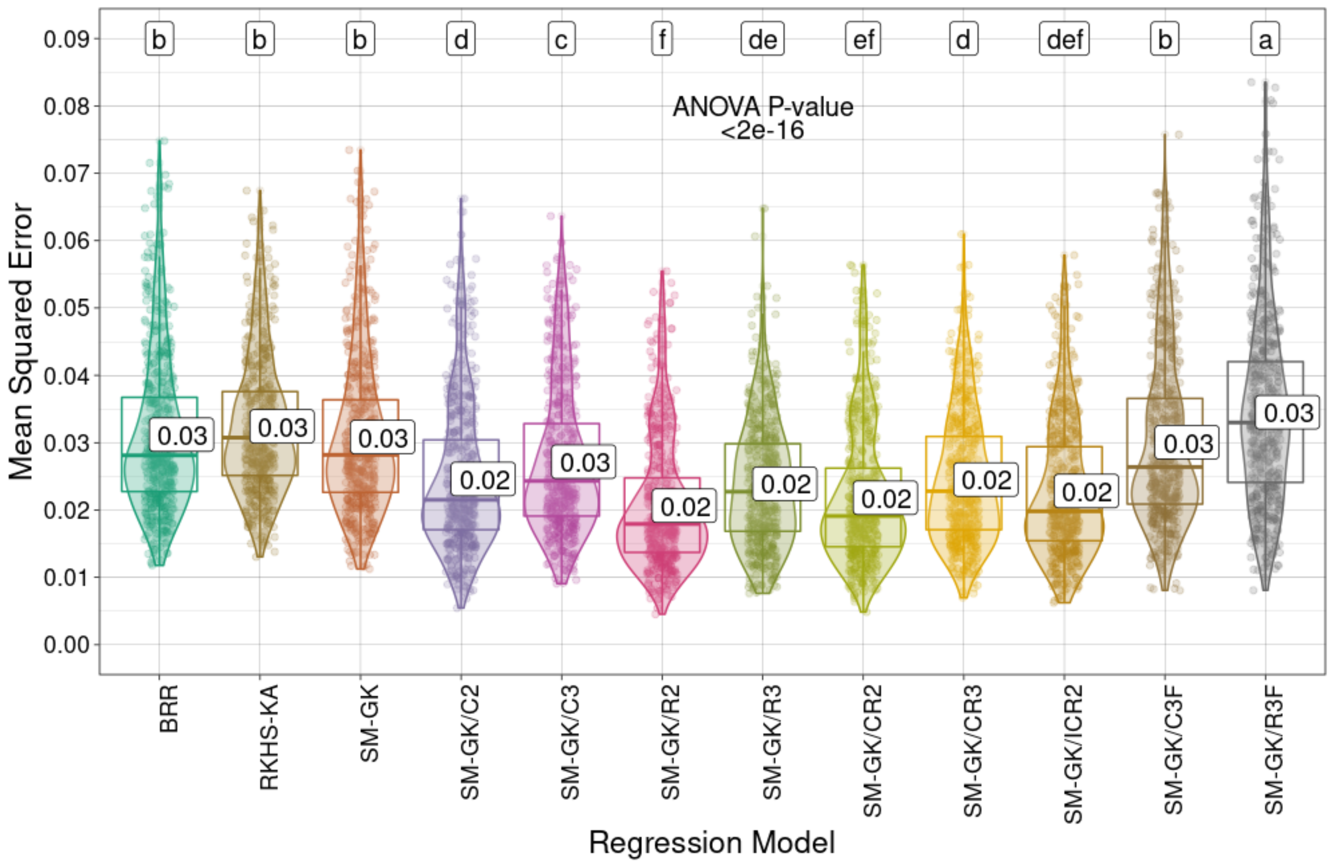


**Supplementary Fig. S46**. Prediction performances (R Pearson correlation coefficients and mean squared errors) for *Urochloa decumbens* leaf percentage (LP) using the approaches employed for regression (Bayesian ridge regression (BRR), reproducing kernel Hilbert space with kernel averaging (RKHS-KA) model, and single-environment, main genotypic effect model with a Gaussian kernel (SM-GK)) and the markers selected through feature selection (FS) methods considering classification (C) and regression (R) algorithms and the intersection of the three tested techniques (C3/R3) or at least two of them (C2/R2). The combinations (i) union of C2 and R2 (CR2), (ii) union of C3 and R3 (CR3), and (iii) intersection of C2 and R2 (ICR2) were also evaluated, as well as the addition of C3 and R3 as fixed effects (C3F and R3F). Mean values are

presented in the boxplots, and the labels at the top of the plots represent the results from Tukey's multiple comparisons test (p value of 0.05).

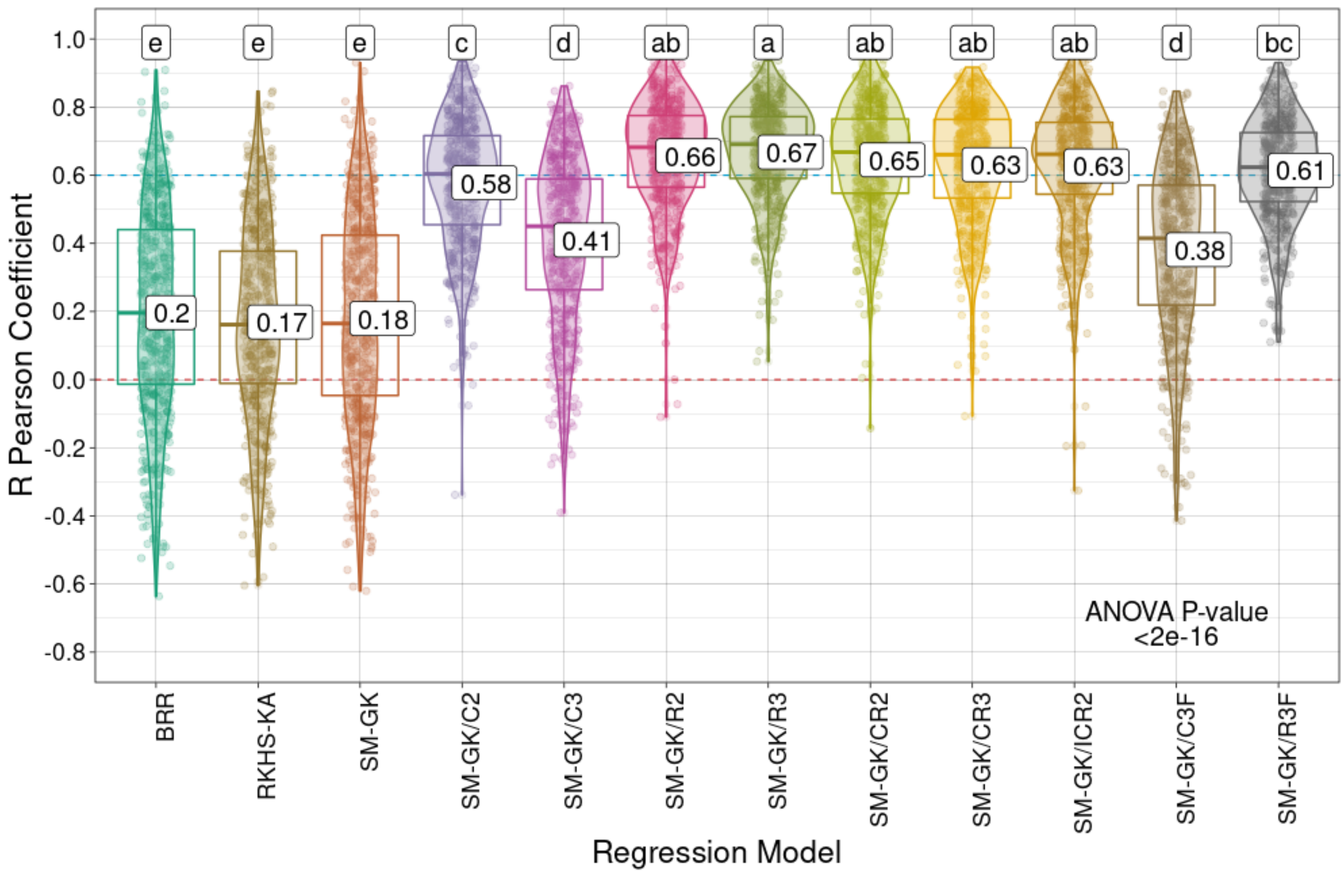

R Pearson Coefficient
Regression Model
1.0
0.8
0.6
0.4
0.2
0.0
-0.2
-0.4
-0.6
-0.8
e
e
e
c
d
ab
a
ab
ab
ab
d
bc
0.2
0.17
0.18
0.58
0.41
0.66
0.67
0.65
0.63
0.63
0.38
0.61
ANOVA P-value
<2e-16
BRR
RKHS-KA
SM-GK
SM-GK/C2
SM-GK/C3
SM-GK/R2
SM-GK/R3
SM-GK/CR2
SM-GK/CR3
SM-GK/ICR2
SM-GK/C3F
SM-GK/R3F

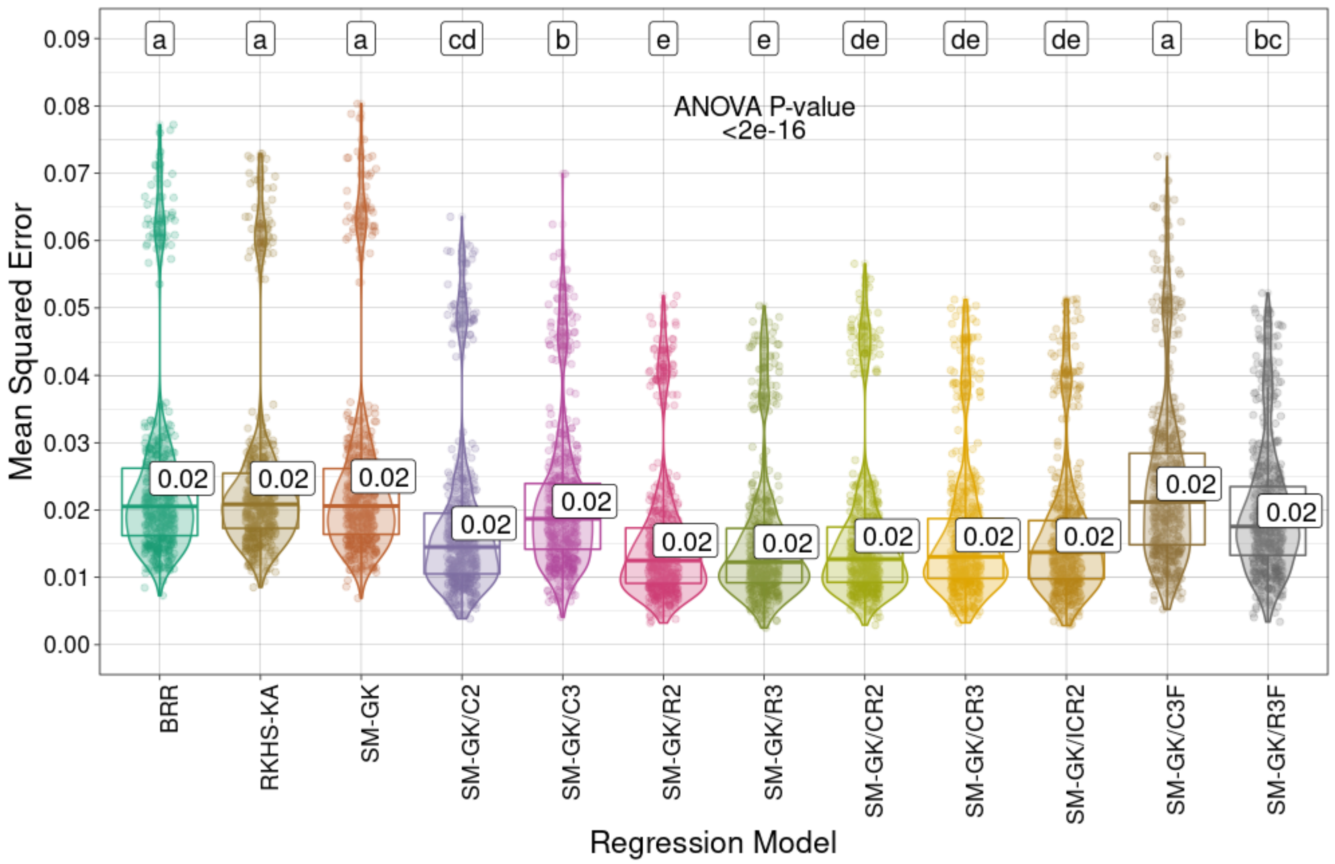


**Supplementary Fig. S47**. Prediction performances (R Pearson correlation coefficients and mean squared errors) for the *Urochloa decumbens* leaf stem ratio (LSR) using the approaches employed for regression (Bayesian ridge regression (BRR), reproducing kernel Hilbert space with kernel averaging (RKHS-KA) model, and single-environment, main genotypic effect model with a Gaussian kernel (SM-GK)) and the markers selected through feature selection (FS) methods considering classification (C) and regression (R) algorithms and the intersection of the three tested techniques (C3/R3) or at least two of them (C2/R2). The combinations (i) union of C2 and R2 (CR2), (ii) union of C3 and R3 (CR3), and (iii) intersection of C2 and R2 (ICR2) were also evaluated, as well as the addition of C3 and R3 as fixed effects (C3F and R3F). Mean

values are presented in the boxplots, and the labels at the top of the plots represent the results from Tukey's multiple comparisons test (p value of 0.05).

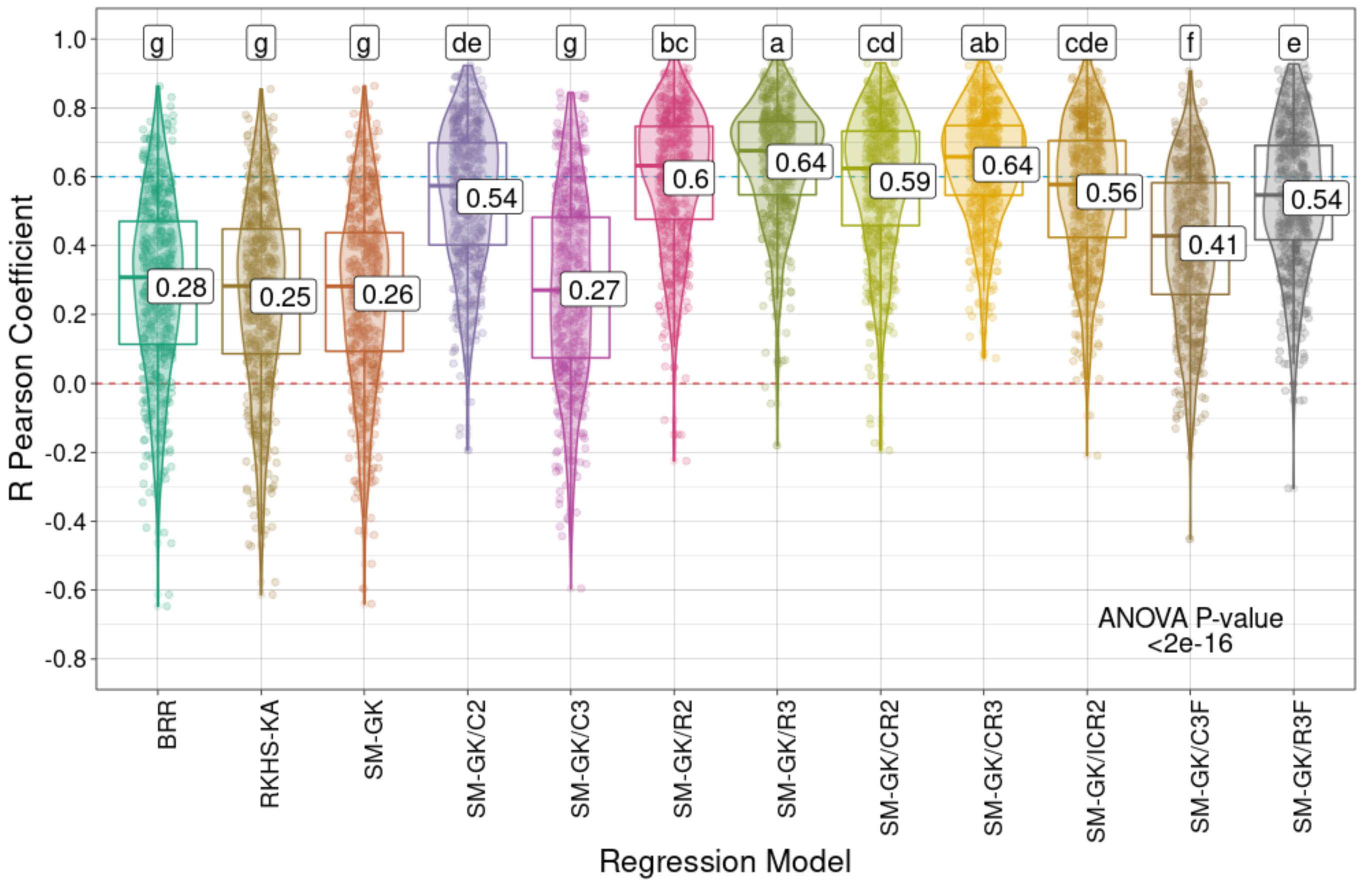
R Pearson Coefficient
Regression Model
g
g
g
de
g
bc
a
cd
ab
cde
f
e
0.28
0.25
0.26
0.54
0.27
0.6
0.64
0.59
0.64
0.56
0.41
0.54
ANOVA P-value
<2e-16
BRR
RKHS-KA
SM-GK
SM-GK/C2
SM-GK/C3
SM-GK/R2
SM-GK/R3
SM-GK/CR2
SM-GK/CR3
SM-GK/ICR2
SM-GK/C3F
SM-GK/R3F

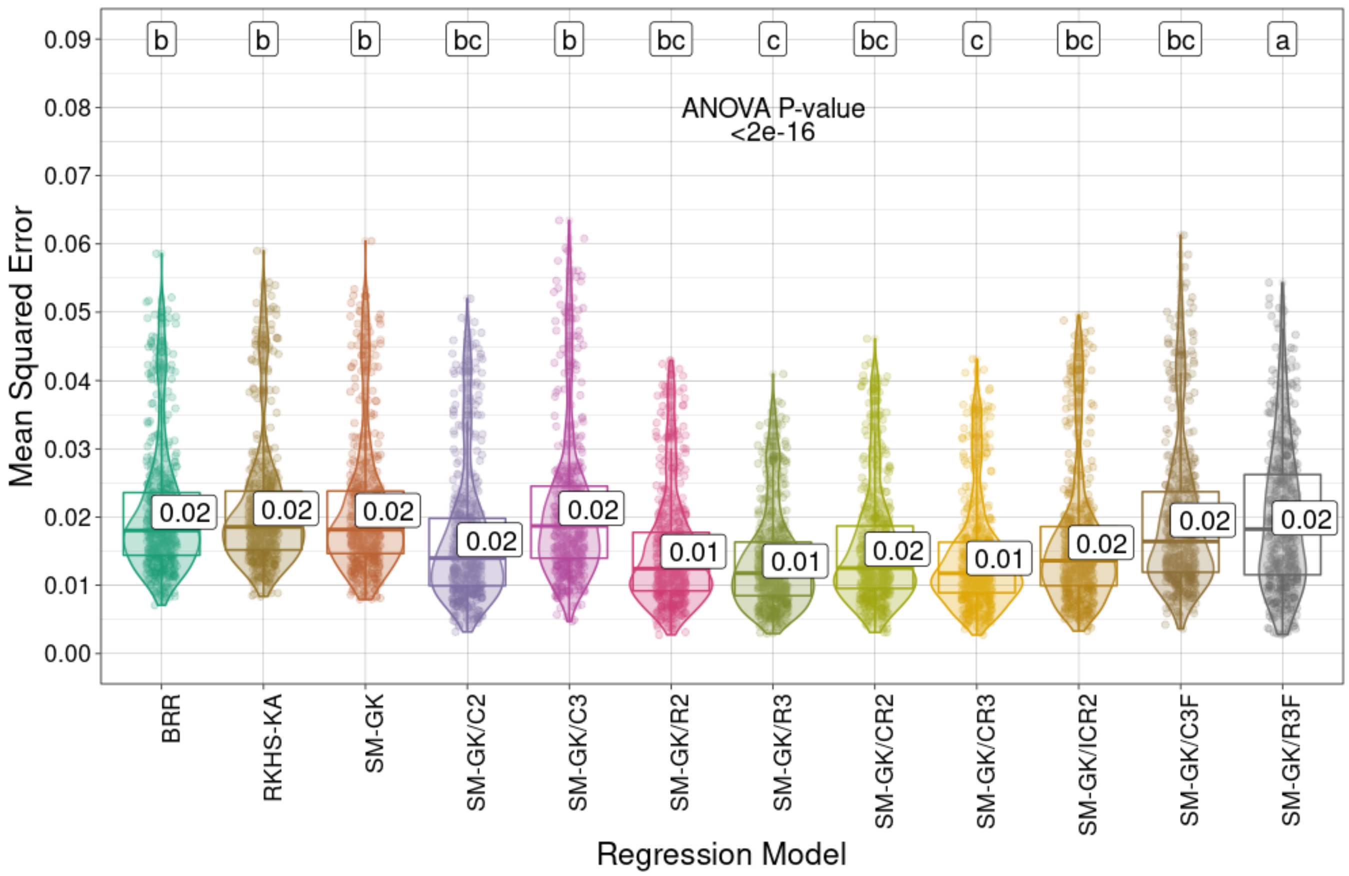


**Supplementary Fig. S48**. Prediction performances (R Pearson correlation coefficients and mean squared errors) for *Urochloa decumbens* regrowth capacity (RC) using the approaches employed for regression (Bayesian ridge regression (BRR), reproducing kernel Hilbert space with kernel averaging (RKHS-KA) model, and single-environment, main genotypic effect model with a Gaussian kernel (SM-GK)) and the markers selected through feature selection (FS) methods considering classification (C) and regression (R) algorithms and the intersection of the three tested techniques (C3/R3) or at least two of them (C2/R2). The combinations (i) union of C2 and R2 (CR2), (ii) union of C3 and R3 (CR3), and (iii) intersection of C2 and R2 (ICR2) were also evaluated, as well as the addition of C3 and R3 as fixed effects (C3F and R3F). Mean values are

presented in the boxplots, and the labels at the top of the plots represent the results from Tukey's multiple comparisons test (p value of 0.05).

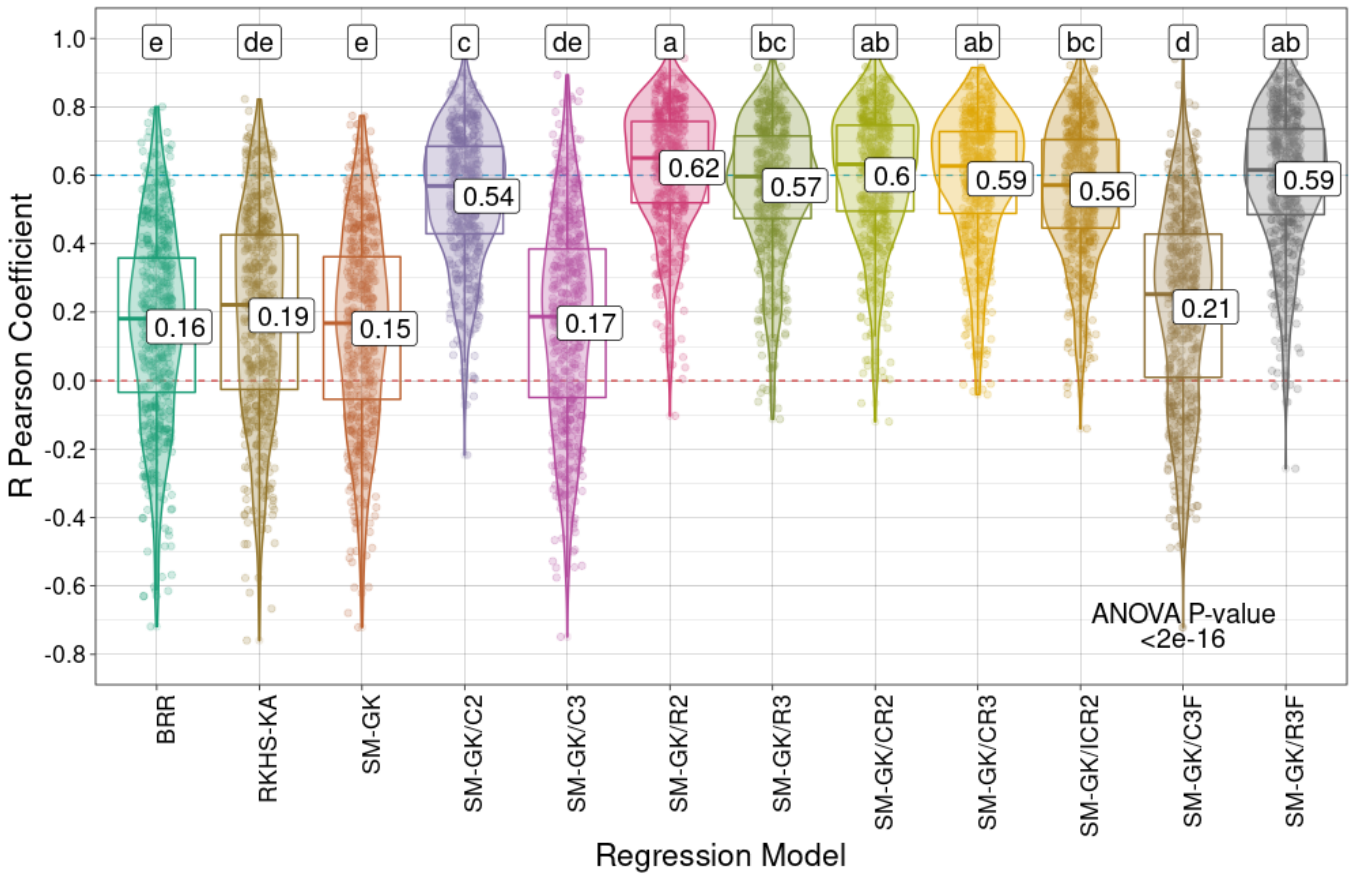

R Pearson Coefficient
1.0
0.8
0.6
0.4
0.2
0.0
-0.2
-0.4
-0.6
-0.8
e
de
e
c
de
a
bc
ab
ab
bc
d
ab
0.16
0.19
0.15
0.54
0.17
0.62
0.57
0.6
0.59
0.56
0.21
0.59
ANOVA P-value
<2e-16
BRR
RKHS-KA
SM-GK
SM-GK/C2
SM-GK/C3
SM-GK/R2
SM-GK/R3
SM-GK/CR2
SM-GK/CR3
SM-GK/ICR2
SM-GK/C3F
SM-GK/R3F
Regression Model

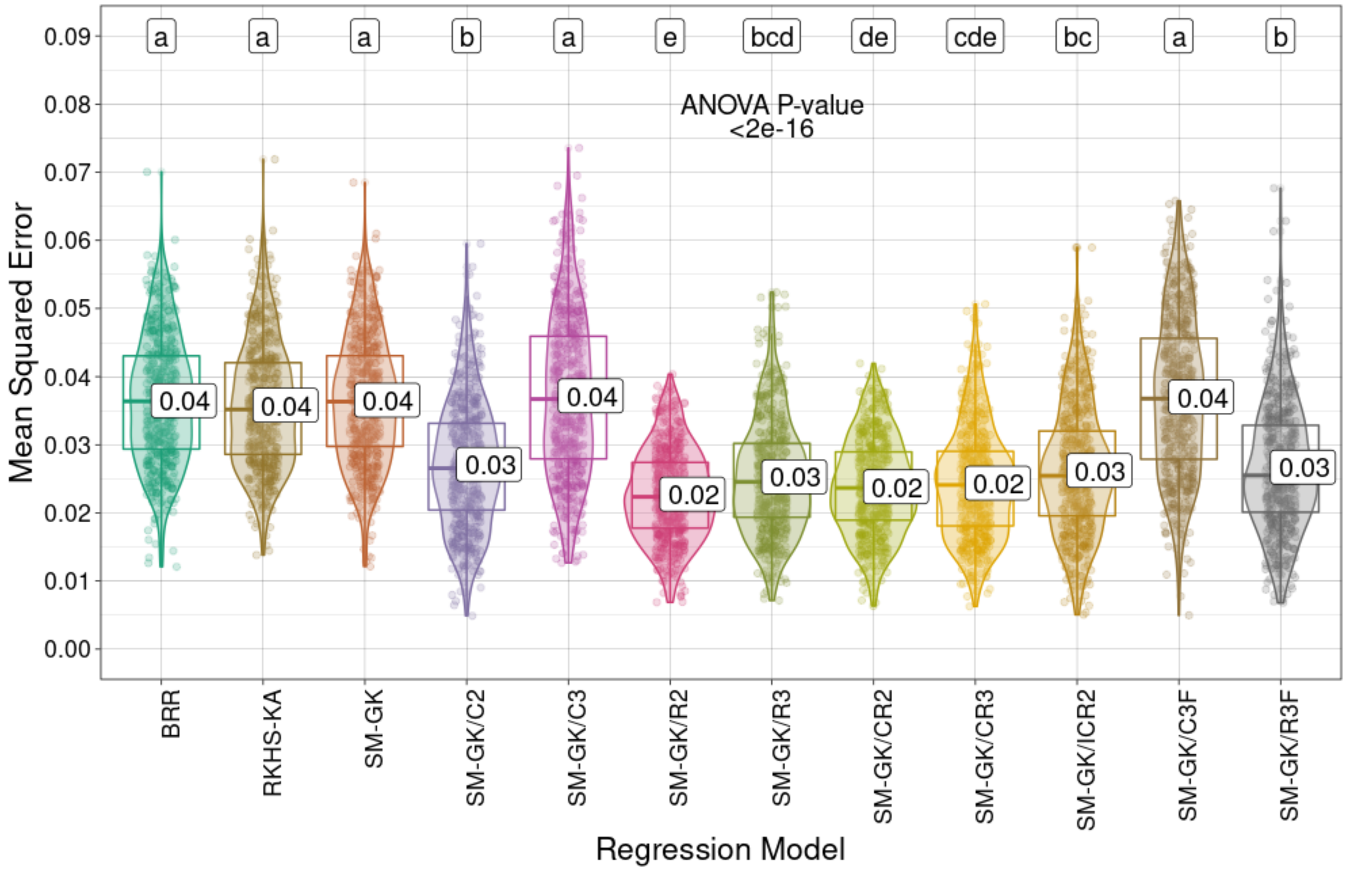


**Supplementary Fig. S49**. Prediction performances (R Pearson correlation coefficients and mean squared errors) for *Urochloa decumbens* total dry matter (TDM) using the approaches employed for regression (Bayesian ridge regression (BRR), reproducing kernel Hilbert space with kernel averaging (RKHS-KA) model, and single-environment, main genotypic effect model with a Gaussian kernel (SM-GK)) and the markers selected through feature selection (FS) methods considering classification (C) and regression (R) algorithms and the intersection of the three tested techniques (C3/R3) or at least two of them (C2/R2). The combinations (i) union of C2 and R2 (CR2), (ii) union of C3 and R3 (CR3), and (iii) intersection of C2 and R2 (ICR2) were also evaluated, as well as the addition of C3 and R3 as fixed effects (C3F and R3F). Mean values are

presented in the boxplots, and the labels at the top of the plots represent the results from Tukey's multiple comparisons test (p value of 0.05).

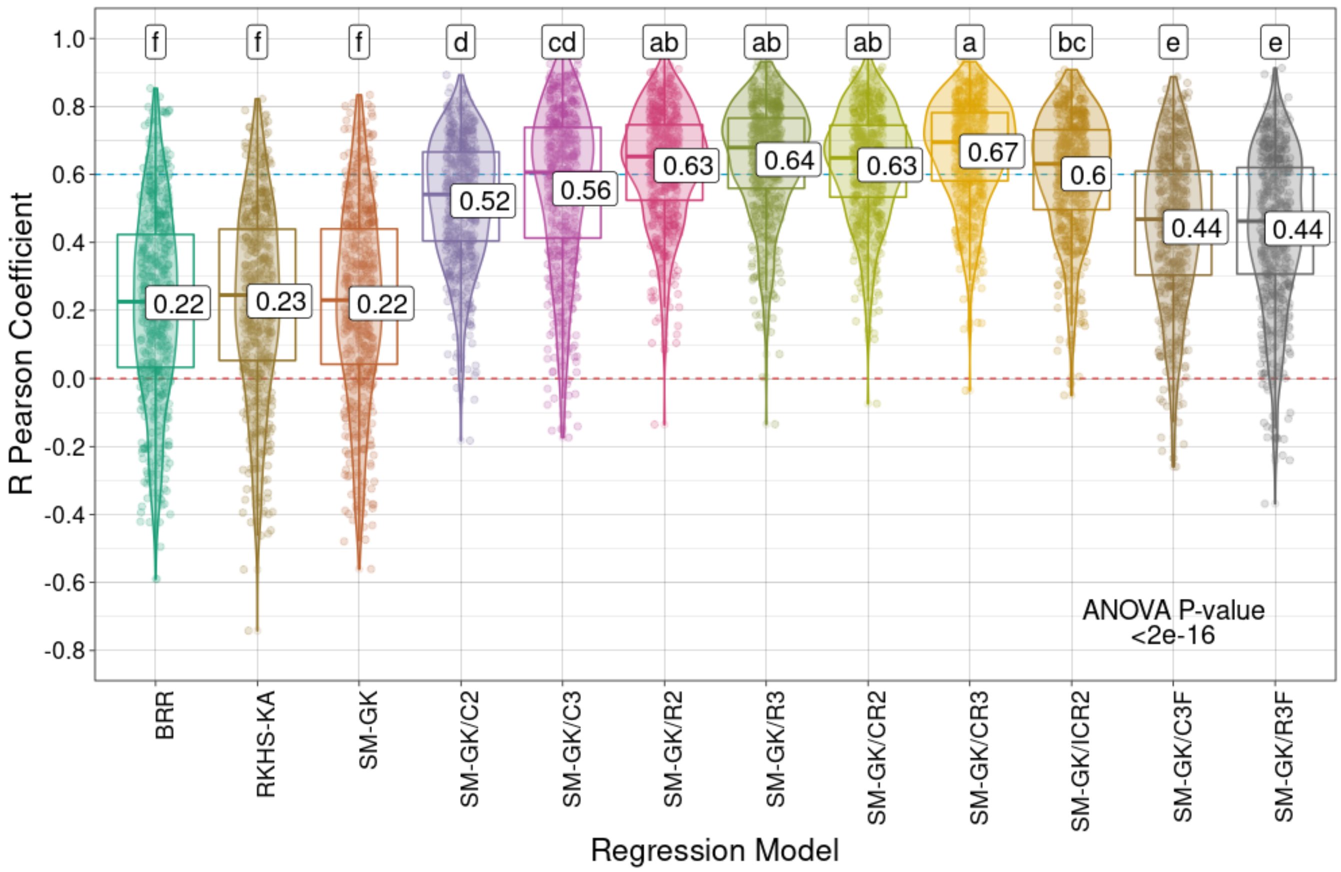

R Pearson Coefficient
1.0
0.8
0.6
0.4
0.2
0.0
-0.2
-0.4
-0.6
-0.8
f
f
f
d
cd
ab
ab
ab
a
bc
e
e
0.22
0.23
0.22
0.52
0.56
0.63
0.64
0.63
0.67
0.6
0.44
0.44
ANOVA P-value
<2e-16
BRR
RKHS-KA
SM-GK
SM-GK/C2
SM-GK/C3
SM-GK/R2
SM-GK/R3
SM-GK/CR2
SM-GK/CR3
SM-GK/ICR2
SM-GK/C3F
SM-GK/R3F
Regression Model

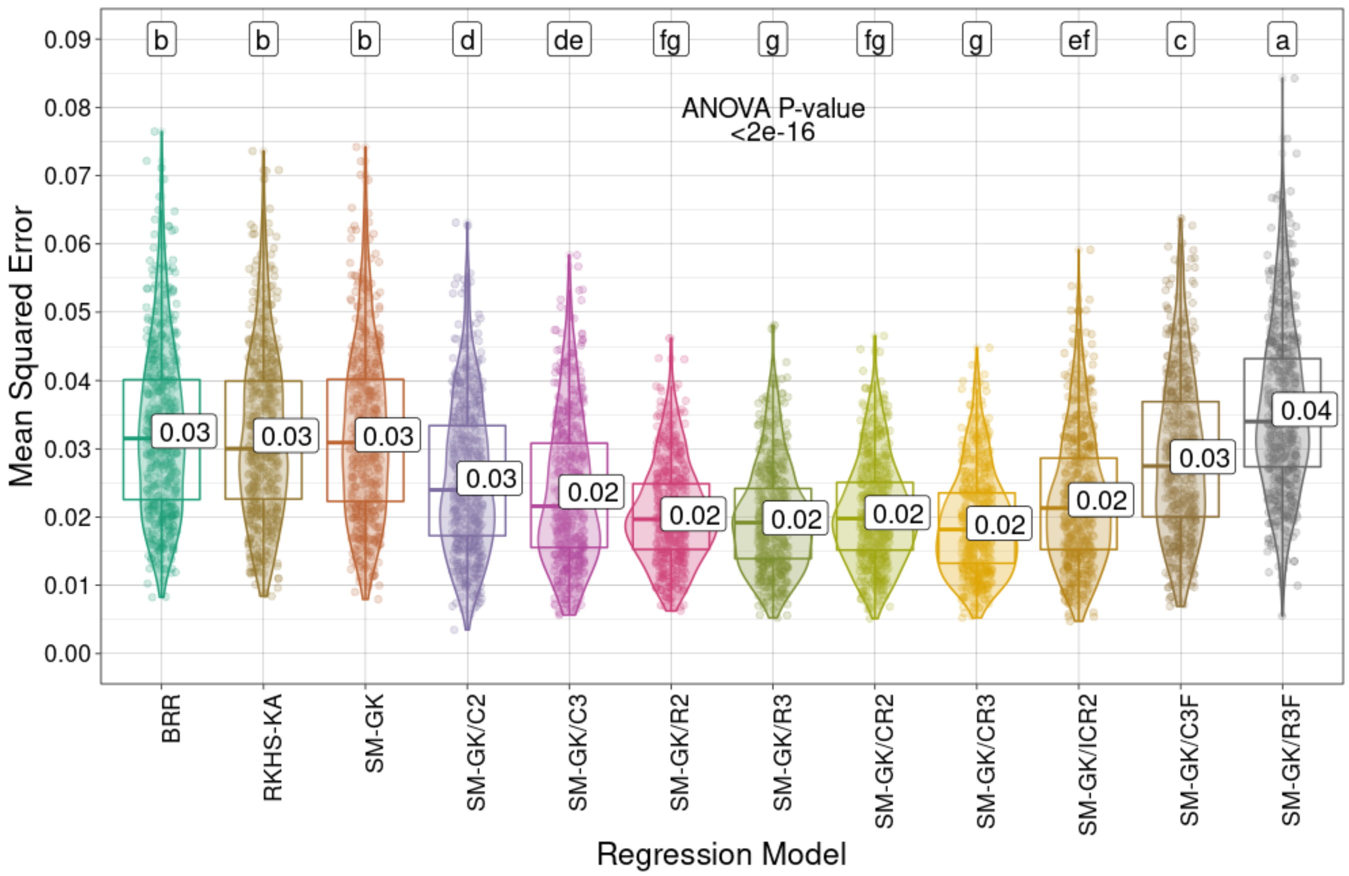


**Supplementary Fig. S50**. Prediction performances (R Pearson correlation coefficients and mean squared errors) for *Megathyrsus maximus* green matter (GM) using the approaches employed for regression (Bayesian ridge regression (BRR), reproducing kernel Hilbert space with kernel averaging (RKHS-KA) model, and single-environment, main genotypic effect model with a Gaussian kernel (SM-GK)) and the markers selected through feature selection (FS) methods considering classification (C) and regression (R) algorithms and the intersection of the three tested techniques (C3/R3) or at least two of them (C2/R2). The combinations (i) union of C2 and R2 (CR2), (ii) union of C3 and R3 (CR3), and (iii) intersection of C2 and R2 (ICR2) were also evaluated, as well as the addition of C3 and R3 as fixed effects (C3F and R3F). Mean values are

presented in the boxplots, and the labels at the top of the plots represent the results from Tukey's multiple comparisons test (p value of 0.05).

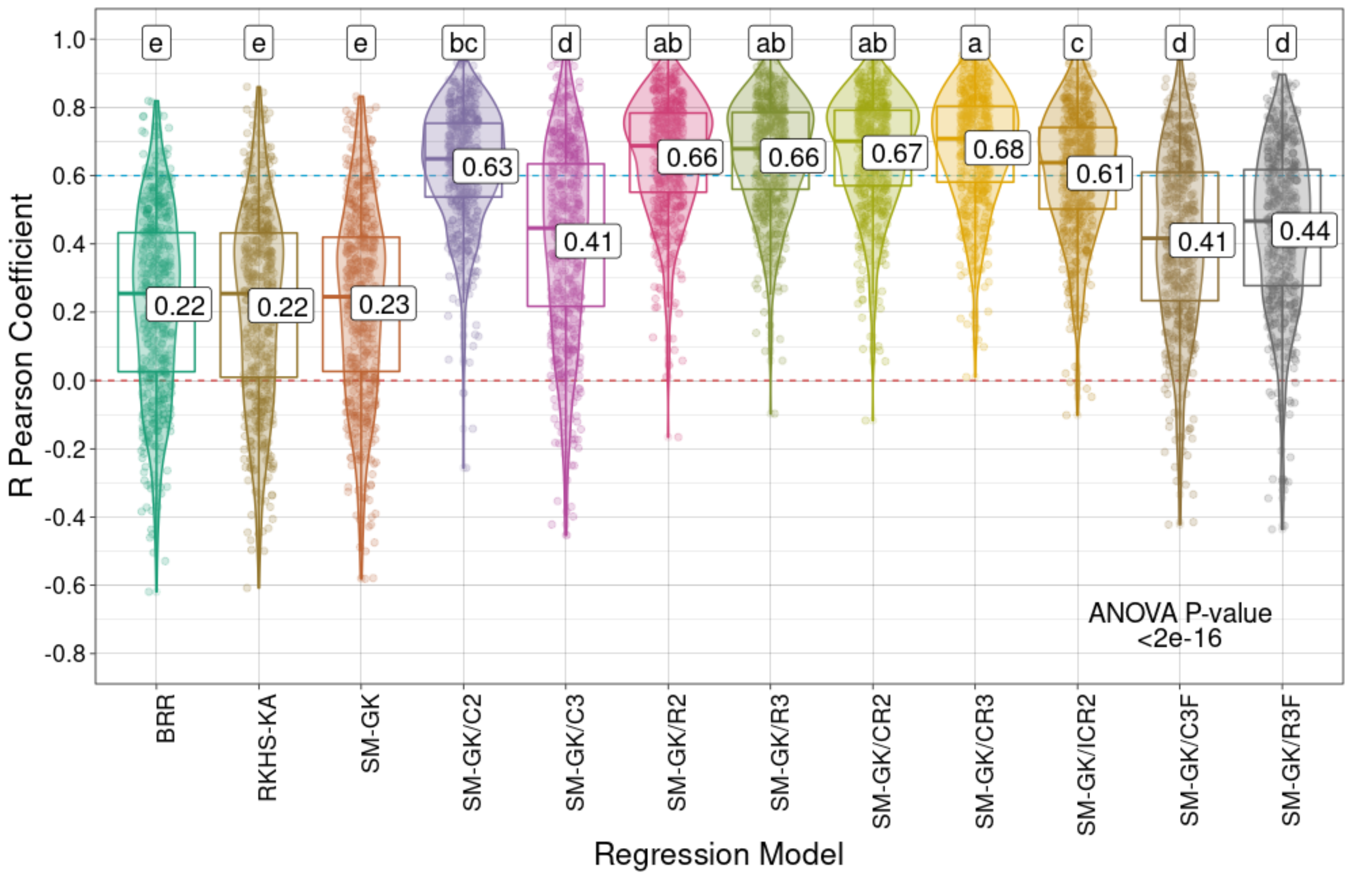
R Pearson Coefficient
1.0
0.8
0.6
0.4
0.2
0.0
-0.2
-0.4
-0.6
-0.8
e
e
e
bc
d
ab
ab
ab
a
c
d
d
0.22
0.22
0.23
0.63
0.41
0.66
0.66
0.67
0.68
0.61
0.41
0.44
ANOVA P-value
<2e-16
BRR
RKHS-KA
SM-GK
SM-GK/C2
SM-GK/C3
SM-GK/R2
SM-GK/R3
SM-GK/CR2
SM-GK/CR3
SM-GK/ICR2
SM-GK/C3F
SM-GK/R3F
Regression Model

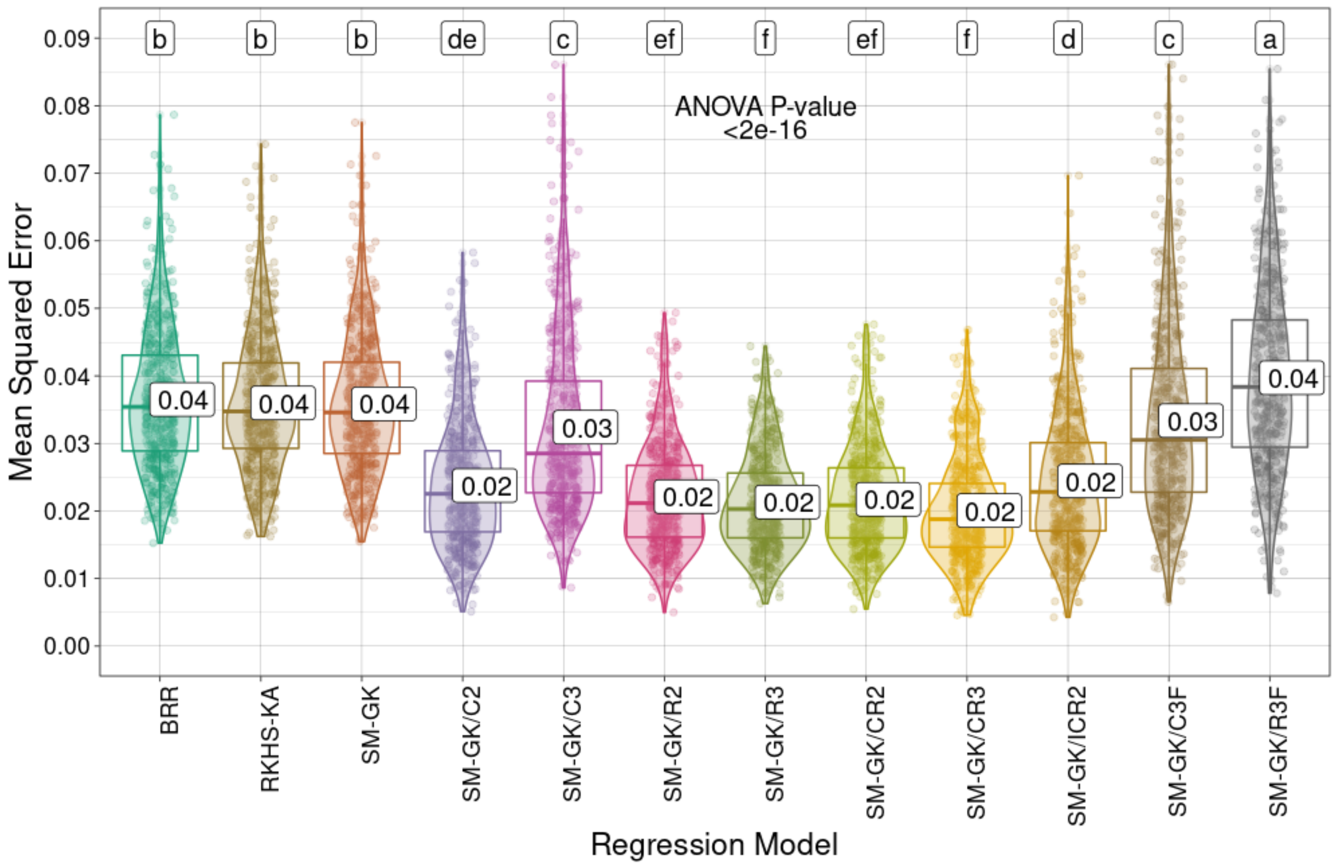


**Supplementary Fig. S51**. Prediction performances (R Pearson correlation coefficients and mean squared errors) for *Megathyrsus maximus* leaf dry matter (LDM) using the approaches employed for regression (Bayesian ridge regression (BRR), reproducing kernel Hilbert space with kernel averaging (RKHS-KA) model, and single-environment, main genotypic effect model with a Gaussian kernel (SM-GK)) and the markers selected through feature selection (FS) methods considering classification (C) and regression (R) algorithms and the intersection of the three tested techniques (C3/R3) or at least two of them (C2/R2). The combinations (i) union of C2 and R2 (CR2), (ii) union of C3 and R3 (CR3), and (iii) intersection of C2 and R2 (ICR2) were also evaluated, as well as the addition of C3 and R3 as fixed effects (C3F and R3F). Mean values are

presented in the boxplots, and the labels at the top of the plots represent the results from Tukey's multiple comparisons test (p value of 0.05).

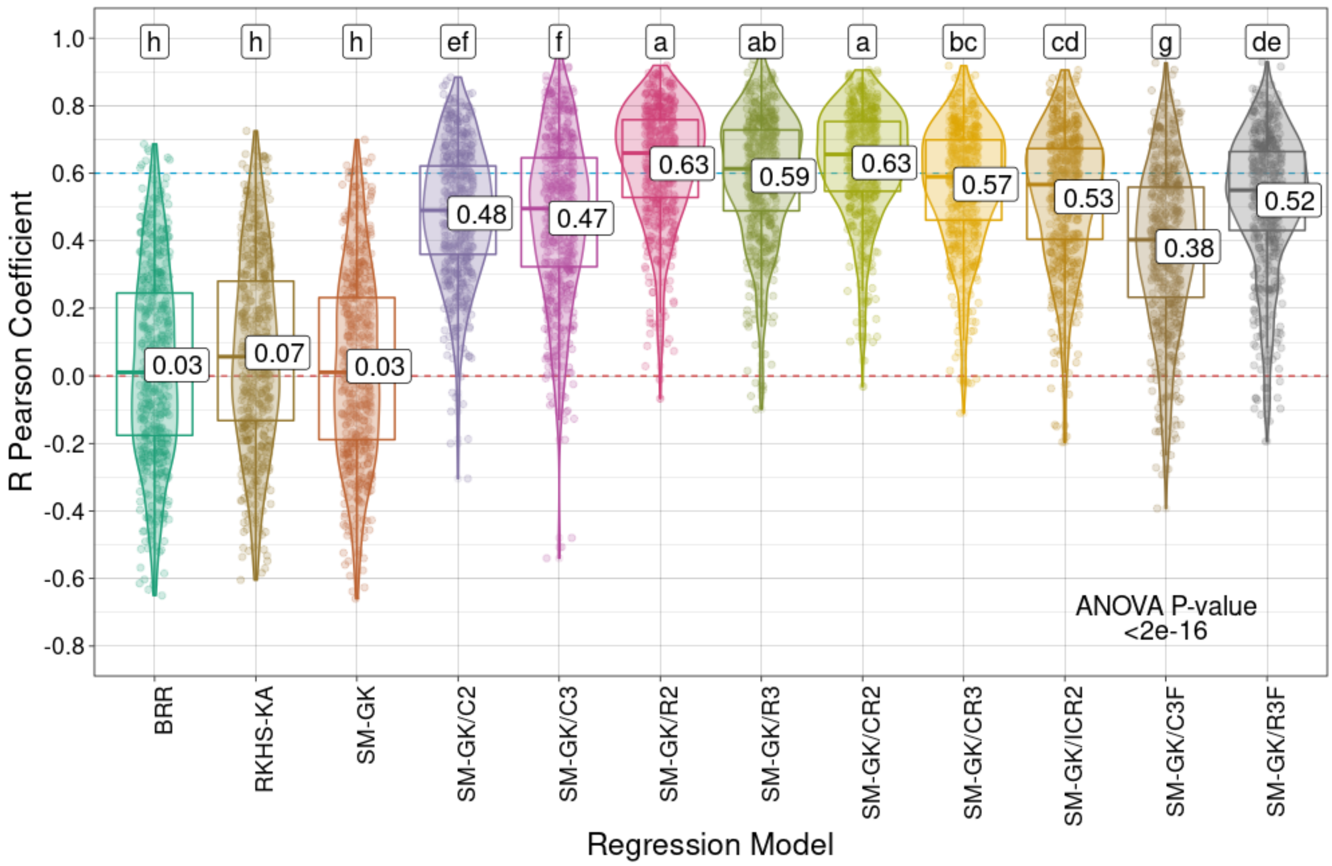
R Pearson Coefficient
Regression Model
BRR
RKHS-KA
SM-GK
SM-GK/C2
SM-GK/C3
SM-GK/R2
SM-GK/R3
SM-GK/CR2
SM-GK/CR3
SM-GK/ICR2
SM-GK/C3F
SM-GK/R3F
h
h
h
ef
f
a
ab
a
bc
cd
g
de
0.03
0.07
0.03
0.48
0.47
0.63
0.59
0.63
0.57
0.53
0.38
0.52
ANOVA P-value
<2e-16

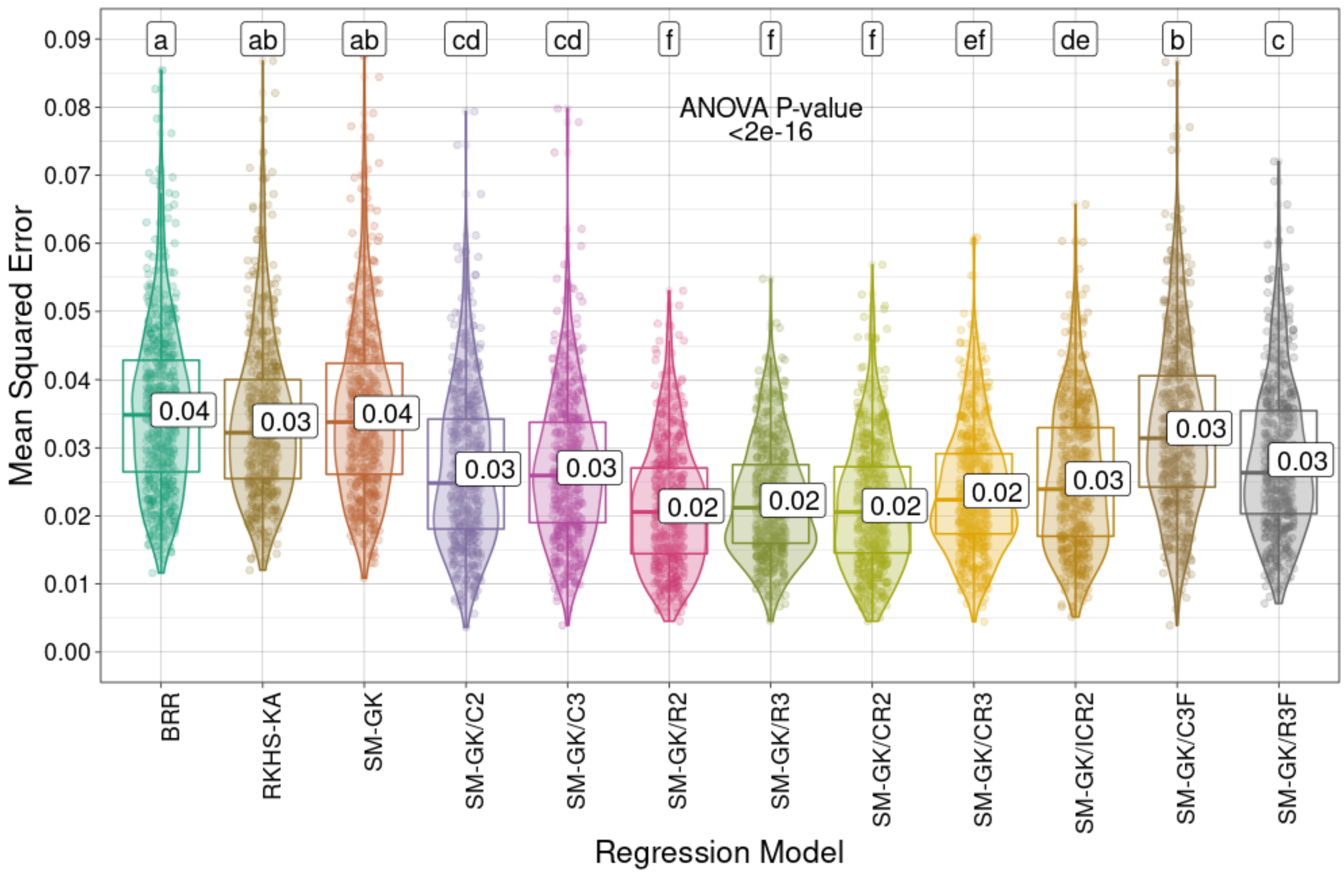


**Supplementary Fig. S52**. Prediction performances (R Pearson correlation coefficients and mean squared errors) for *Megathyrsus maximus* percentage of leaf blade (PLB) using the approaches employed for regression (Bayesian ridge regression (BRR), reproducing kernel Hilbert space with kernel averaging (RKHS-KA) model, and single-environment, main genotypic effect model with a Gaussian kernel (SM-GK)) and the markers selected through feature selection (FS) methods considering classification (C) and regression (R) algorithms and the intersection of the three tested techniques (C3/R3) or at least two of them (C2/R2). The combinations (i) union of C2 and R2 (CR2), (ii) union of C3 and R3 (CR3), and (iii) intersection of C2 and R2 (ICR2) were also evaluated, as well as the addition of C3 and R3 as fixed effects (C3F and R3F). Mean

values are presented in the boxplots, and the labels at the top of the plots represent the results from Tukey's multiple comparisons test (p value of 0.05).

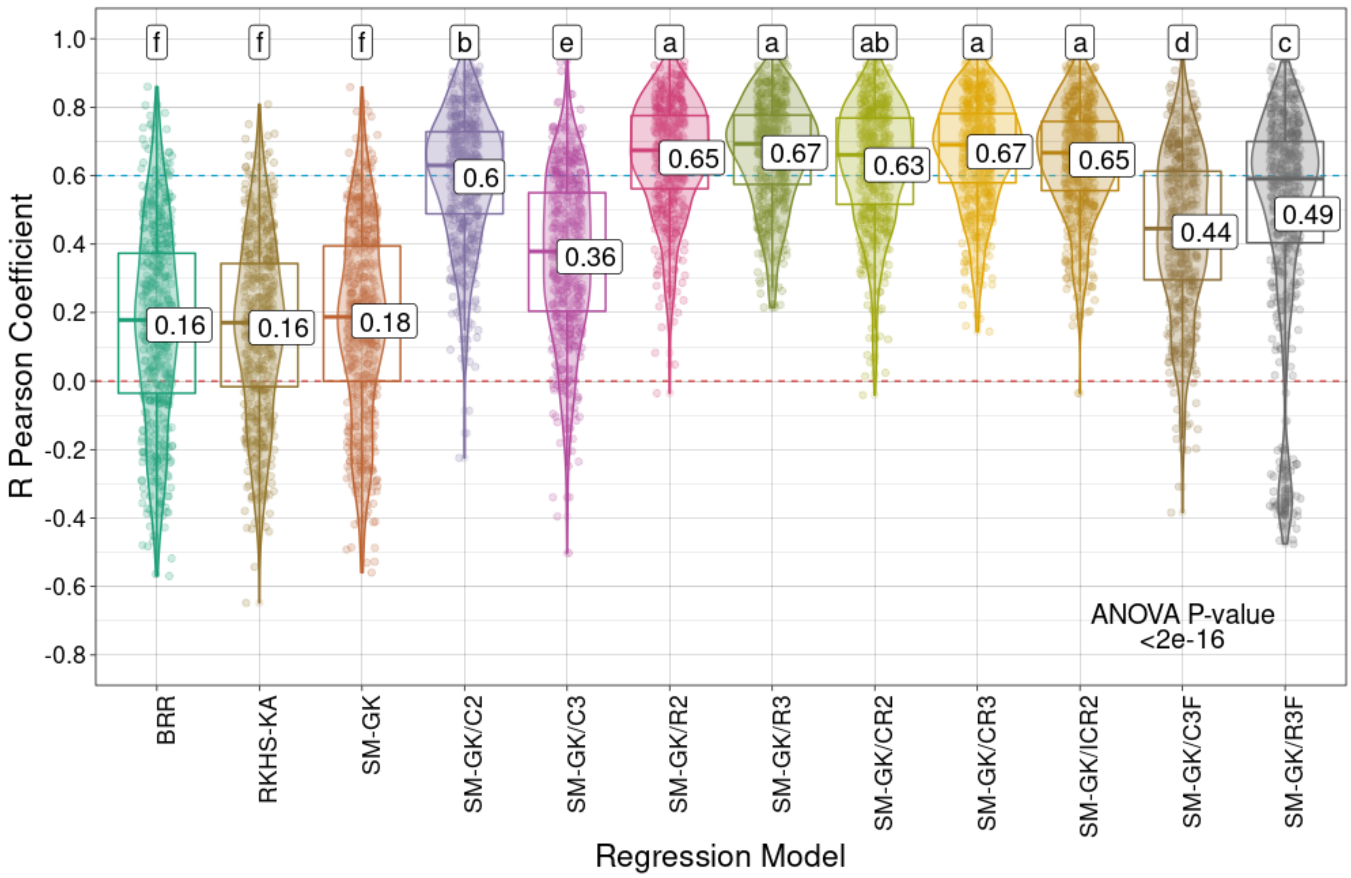
R Pearson Coefficient
Regression Model
1.0
0.8
0.6
0.4
0.2
0.0
-0.2
-0.4
-0.6
-0.8
f
f
f
b
e
a
a
ab
a
a
d
c
0.16
0.16
0.18
0.6
0.36
0.65
0.67
0.63
0.67
0.65
0.44
0.49
ANOVA P-value
<2e-16
BRR
RKHS-KA
SM-GK
SM-GK/C2
SM-GK/C3
SM-GK/R2
SM-GK/R3
SM-GK/CR2
SM-GK/CR3
SM-GK/ICR2
SM-GK/C3F
SM-GK/R3F

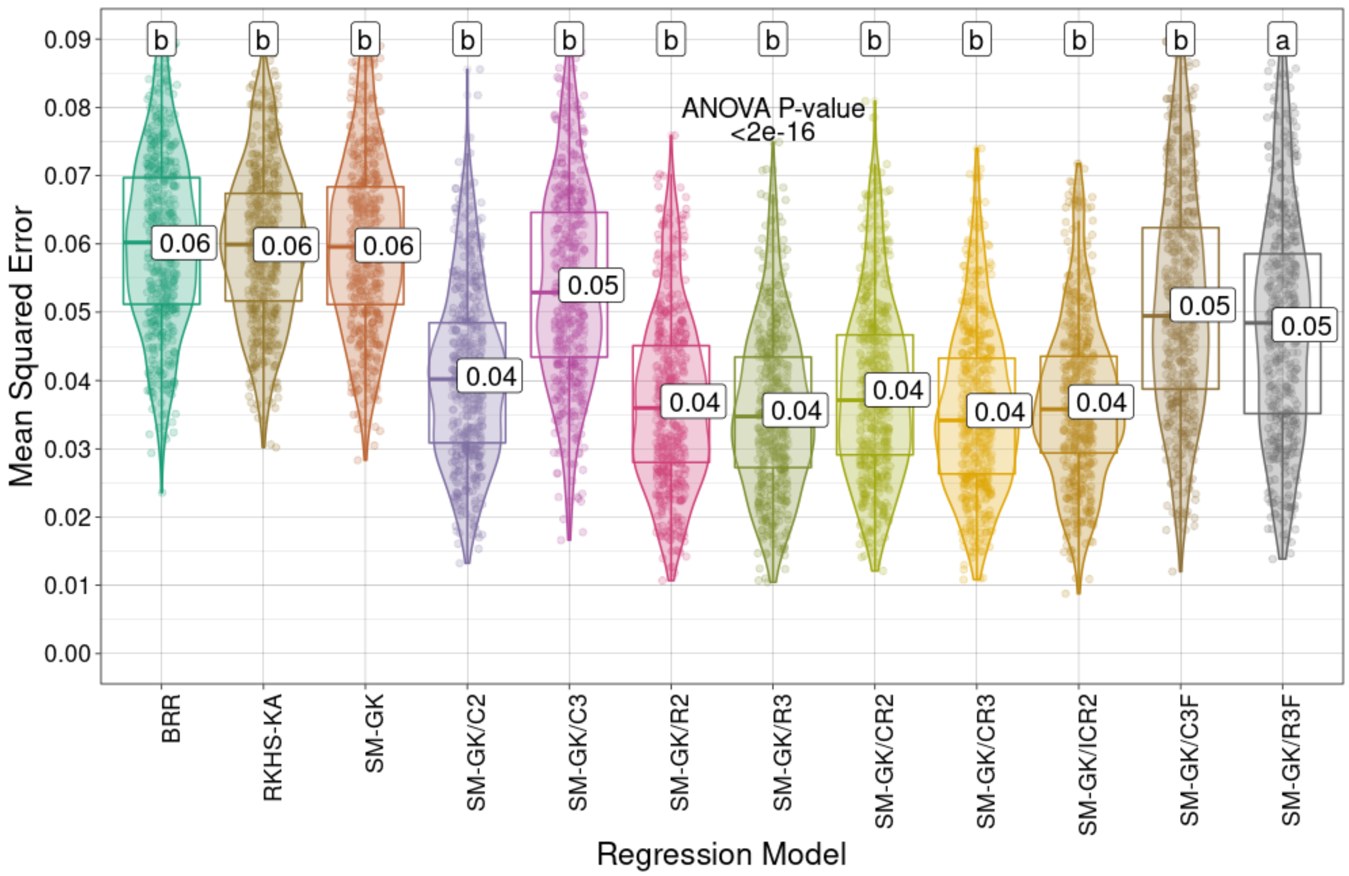


**Supplementary Fig. S53**. Prediction performances (R Pearson correlation coefficients and mean squared errors) for *Megathyrsus maximus* regrowth capacity (RC) using the approaches employed for regression (Bayesian ridge regression (BRR), reproducing kernel Hilbert space with kernel averaging (RKHS-KA) model, and single-environment, main genotypic effect model with a Gaussian kernel (SM-GK)) and the markers selected through feature selection (FS) methods considering classification (C) and regression (R) algorithms and the intersection of the three tested techniques (C3/R3) or at least two of them (C2/R2). The combinations (i) union of C2 and R2 (CR2), (ii) union of C3 and R3 (CR3), and (iii) intersection of C2 and R2 (ICR2) were also evaluated, as well as the addition of C3 and R3 as fixed effects (C3F and R3F). Mean

values are presented in the boxplots, and the labels at the top of the plots represent the results from Tukey's multiple comparisons test (p value of 0.05).

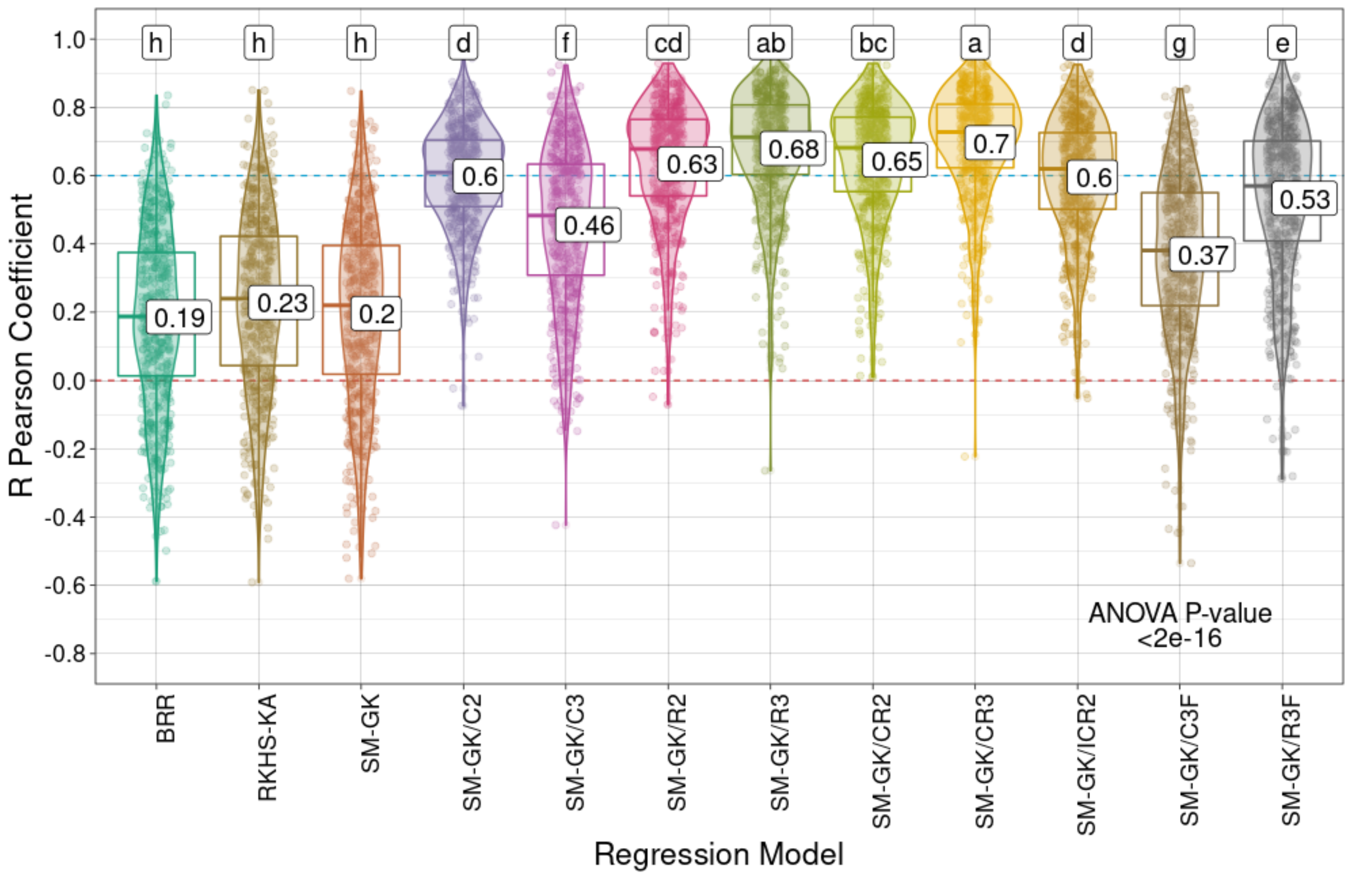

R Pearson Coefficient
Regression Model
h
h
h
d
f
cd
ab
bc
a
d
g
e
0.19
0.23
0.2
0.6
0.46
0.63
0.68
0.65
0.7
0.6
0.37
0.53
ANOVA P-value
<2e-16
BRR
RKHS-KA
SM-GK
SM-GK/C2
SM-GK/C3
SM-GK/R2
SM-GK/R3
SM-GK/CR2
SM-GK/CR3
SM-GK/ICR2
SM-GK/C3F
SM-GK/R3F

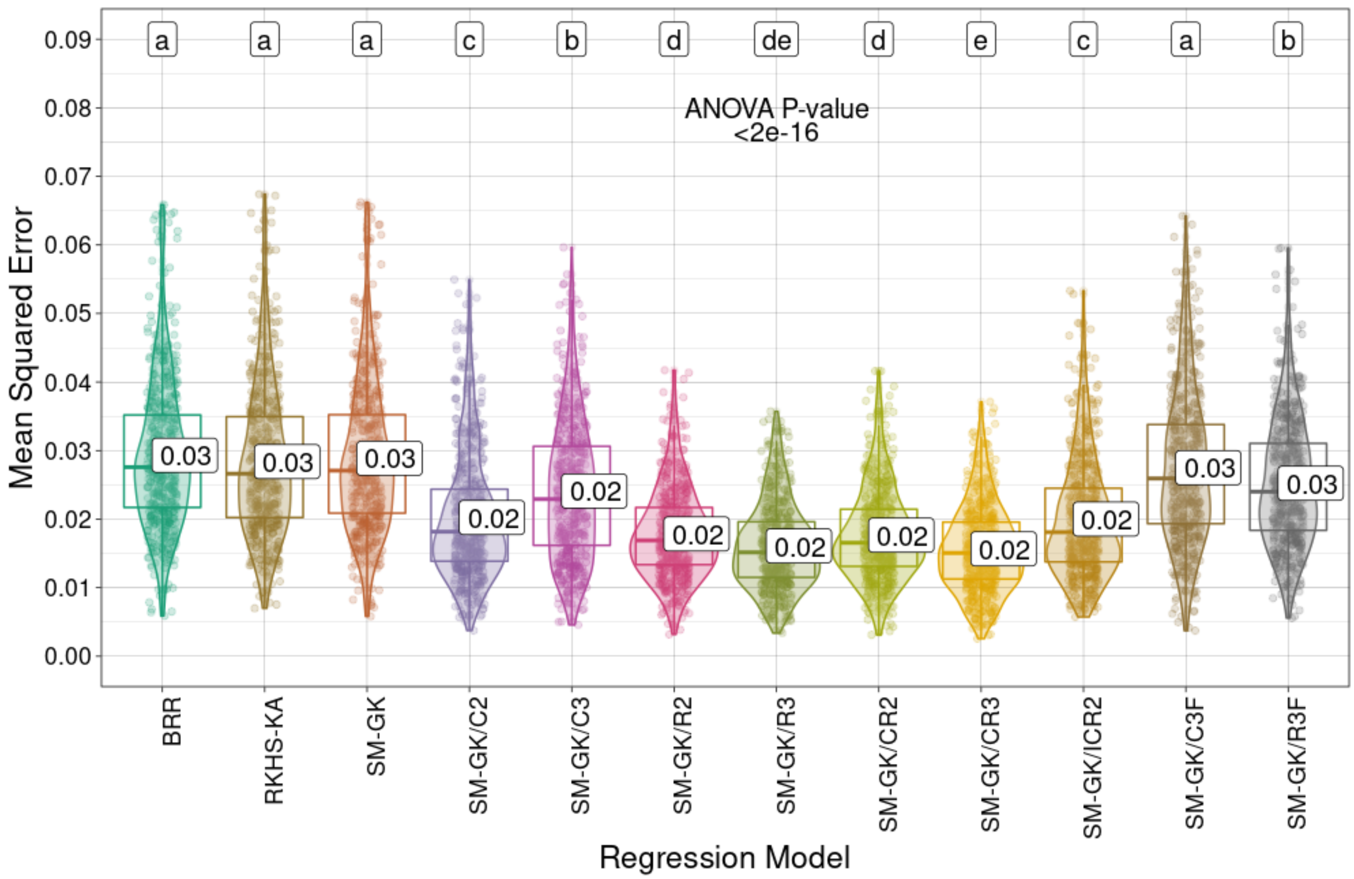


**Supplementary Fig. S54**. Prediction performances (R Pearson correlation coefficients and mean squared errors) for *Megathyrsus maximus* stem dry matter (SDM) using the approaches employed for regression (Bayesian ridge regression (BRR), reproducing kernel Hilbert space with kernel averaging (RKHS-KA) model, and single-environment, main genotypic effect model with a Gaussian kernel (SM-GK)) and the markers selected through feature selection (FS) methods considering classification (C) and regression (R) algorithms and the intersection of the three tested techniques (C3/R3) or at least two of them (C2/R2). The combinations (i) union of C2 and R2 (CR2), (ii) union of C3 and R3 (CR3), and (iii) intersection of C2 and R2 (ICR2) were also evaluated, as well as the addition of C3 and R3 as fixed effects (C3F and R3F). Mean

values are presented in the boxplots, and the labels at the top of the plots represent the results from Tukey's multiple comparisons test (p value of 0.05).

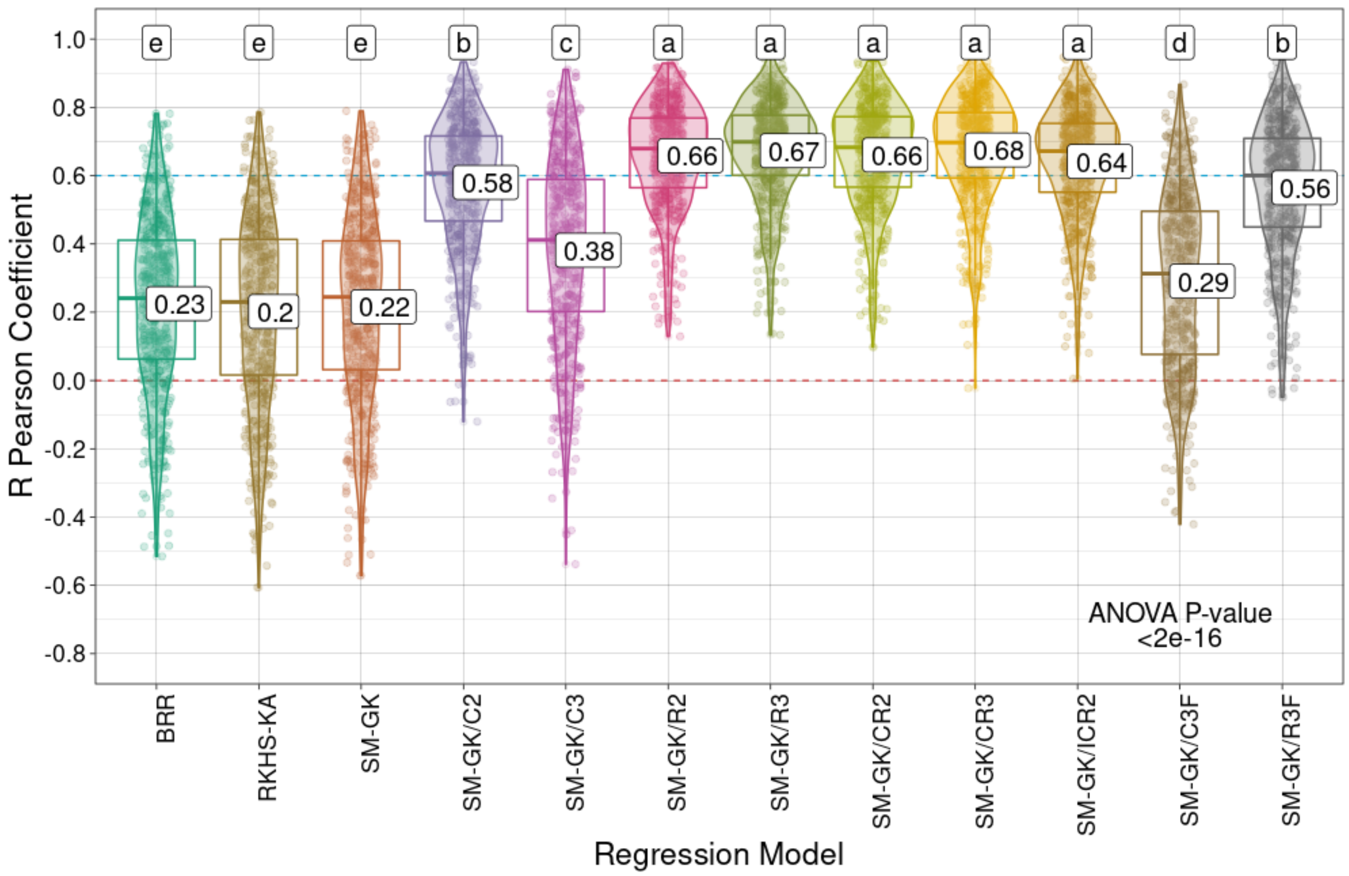

R Pearson Coefficient
1.0
0.8
0.6
0.4
0.2
0.0
-0.2
-0.4
-0.6
-0.8
e
e
e
b
c
a
a
a
a
a
d
b
0.23
0.2
0.22
0.58
0.38
0.66
0.67
0.66
0.68
0.64
0.29
0.56
ANOVA P-value
<2e-16
BRR
RKHS-KA
SM-GK
SM-GK/C2
SM-GK/C3
SM-GK/R2
SM-GK/R3
SM-GK/CR2
SM-GK/CR3
SM-GK/ICR2
SM-GK/C3F
SM-GK/R3F
Regression Model

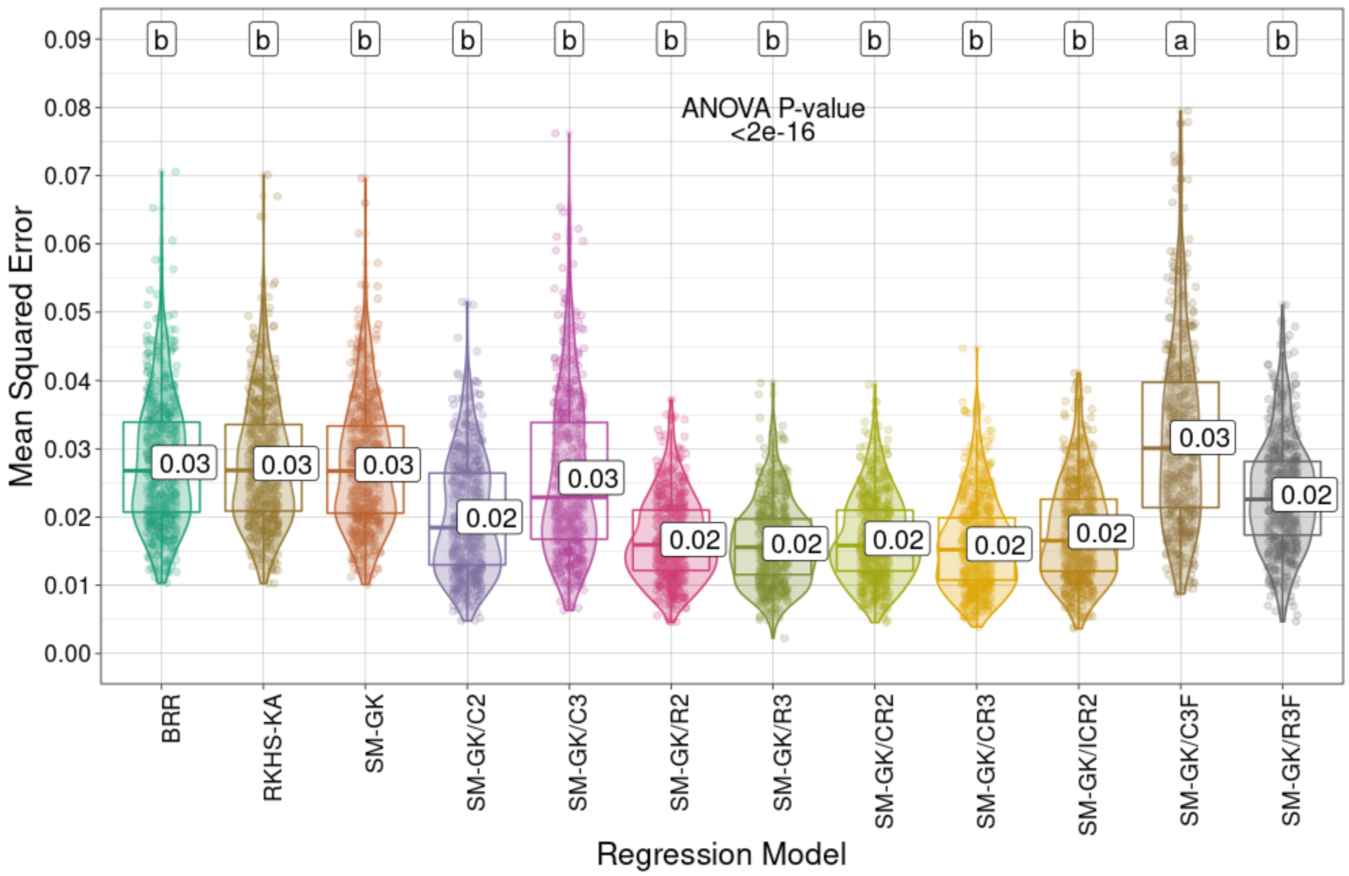


**Supplementary Fig. S55**. Prediction performances (R Pearson correlation coefficients and mean squared errors) for *Megathyrsus maximus* total dry matter (TDM) using the approaches employed for regression (Bayesian ridge regression (BRR), reproducing kernel Hilbert space with kernel averaging (RKHS-KA) model, and single-environment, main genotypic effect model with a Gaussian kernel (SM-GK)) and the markers selected through feature selection (FS) methods considering classification (C) and regression (R) algorithms and the intersection of the three tested techniques (C3/R3) or at least two of them (C2/R2). The combinations (i) union of C2 and R2 (CR2), (ii) union of C3 and R3 (CR3), and (iii) intersection of C2 and R2 (ICR2) were also evaluated, as well as the addition of C3 and R3 as fixed effects (C3F and R3F). Mean

values are presented in the boxplots, and the labels at the top of the plots represent the results from Tukey's multiple comparisons test (p value of 0.05).
